# Rethinking pain threshold as a zone of uncertainty

**DOI:** 10.1101/521302

**Authors:** Victoria J Madden, Peter R Kamerman, Mark J Catley, Valeria Bellan, Leslie N Russek, Danny Camfferman, G Lorimer Moseley

**Affiliations:** Department of Anaesthesia and Perioperative Medicine, Department of Psychiatry and Mental Health, University of Cape Town, Cape Town, Western Cape, South Africa; Sansom Institute for Health Research, University of South Australia, Adelaide, South Australia, Australia; School of Physiology, Faculty of Health Sciences, University of the Witwatersrand, Johannesburg, Gauteng, South Africa; School of Pharmacy and Biomedical Sciences, Faculty of Health Sciences, Curtin University, Perth, Western Australia, Australia; Department of Physical Therapy, Clarkson University, Potsdam, New York, USA

## Abstract

**Background:** The pain threshold is traditionally conceptualised as a boundary that lies between painful and non-painful events, suggesting a reasonably stable relationship between stimulus and response. In two previous experiments, participants received laser stimuli of various intensities and rated each stimulus on the Sensation and Pain Rating Scale (SPARS), which includes ranges for rating painful and non-painful events and clearly defines the presumed boundary between them. In the second experiment, participants also provided ratings on the conventional 0-100 Numerical Rating Scale for pain (NRS) and a new rating scale for non-painful events. Those data showed the SPARS to have a curvilinear stimulus-response relationship, reflecting that several different intensities may be rated as painful and non-painful in different trials. This suggests that participants were uncertain about painfulness over a range of intensities and calls into question the idea of a boundary between non-painful and painful events. The current study aimed to determine the number of different stimulus intensities across which each participant provided ‘painful’ and ‘non-painful’ reports in different trials.

**Methods:** We undertook novel exploratory analyses on data from the aforementioned two experiments (n = 19, 11 female, 18-31 years old; n = 7, 5 female, 21-30 years old). We used the binomial test to formally determine the width of this ‘zone of uncertainty’ about painfulness, using ratings on the SPARS and the comparator scales, and data visualisation to assess whether trial-to-trial change in stimulus intensity influences ratings.

**Results:** We found that the width of the zone of uncertainty varied notably between individuals and that the zone was non-continuous for most participants. Plots of group-level data concealed the inter-individual variability apparent in the individual plots, but still showed a wide zone of uncertainty on both the SPARS and the NRS, but a narrow zone on the scale for non-painful events. There was no evidence that trial-to-trial change in stimulus intensity influenced ratings.

**Conclusions:** The variability revealed by this study has important design implications for experiments that include initial calibration of repeatedly delivered stimuli. The variability also stands to inflate the size of sample that is required for adequate statistical powering of experiments, and provides rationale for the use of statistical approaches that account for individual variability in studies of pain. Finally, the high variability implies that, if experimental stimuli are to be used in clinical phenotyping, many trials may be required to obtain results that represent a single patient’s actual response profile.

## Introduction

The notion of pain threshold is familiar as the minimum stimulus required to evoke pain. In general, we tend to think we ‘have’ a pain threshold that is unique to us and reasonably stable. This position seems intuitive, yet we regularly encounter situations in which our pain threshold seems to have moved or indeed vanished completely. In the scientific literature, pain threshold is conceptualised as a fixed point (or very narrow range) in the stimulus-response relationship, ^1–3^ and the position of this point (or narrow range) on the stimulus-response curve can be shifted by biological, psychological and social factors (see ^4^ for an accessible review).

What is the system we use to understand and assess pain threshold? Self-report is the reference standard for assessing pain intensity in humans ^5^, and there are various numerical and visual pain rating scales, typically anchored at the extremes with ‘no pain’ and something equivalent to ‘worst pain imaginable’ ^6^. Early studies of the characteristics of these scales demonstrated that, at the group level, the scales have favourable measurement properties and good utility^6^, which has made them popular tools in routine clinical practice and clinical research, but for fundamental research on the psychophysics of pain perception, their utility is limited. For example, in experimental studies, participants are frequently exposed to a range of non-painful and painful stimuli, but scales that are anchored to ‘no pain’ at the lower end of the range do not allow participants to express the full breadth of the non-painful percept. It is also impossible to clearly identify the transition from a non-painful to a painful percept (pain threshold) with a scale that is truncated in this manner.

To address the range limitations of conventional pain rating scales, we previously developed and tested a new scale – the Sensation and Pain Rating Scale^7^ (SPARS, previously called the FEST^8^) – that spans the perceptual range including painful and non-painful percepts, and explicitly identifies the point of transition between stimuli perceived as painful and non-painful. The SPARS consists of a line extending from an anchor point of −50 (labelled: ‘no sensation’) to an anchor point of +50 (labelled: ‘most intense pain you can imagine’), with pain threshold—the intuitively neutral point—represented by 0 (labelled: ‘the exact point at which what you feel transitions to pain’). In addition, the range from −50 to 0 is labelled ‘non-painful’ while the range from 0 to +50 is labelled ‘painful’ (Fig. 1). In practice, participants are advised to first make a decision about whether a stimulus is ‘painful’ or ‘non-painful’ before selecting a point on the appropriate side of the scale that matches the intensity of their experience.

**Figure 1:**
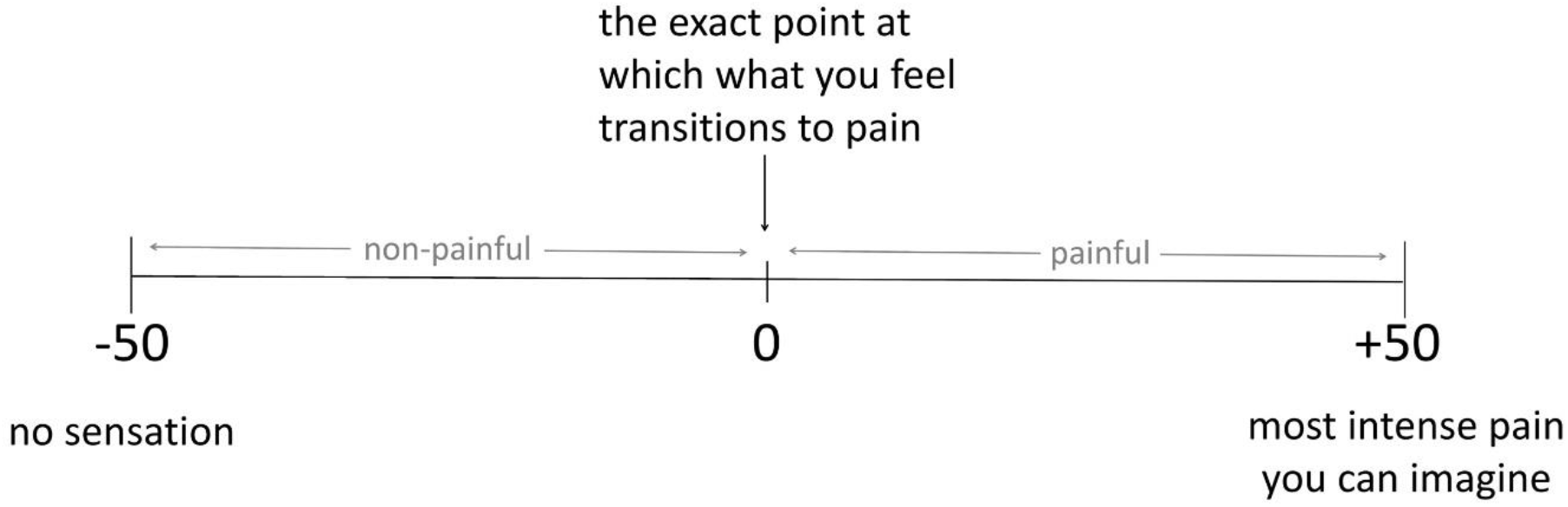
Structure of the Sensation and Pain Rating Scale (SPARS).

We previously tested the SPARS in two experiments involving the application of laser heat stimuli of a range of intensities to the lower back of healthy volunteers^9^. Results of the planned primary analysis have been reported at Madden, et al. ^7^. In the first experiment, participants used only the SPARS to rate stimulus intensity. In the second experiment, participants rated the stimuli using the SPARS, the ‘Sensation Rating Scale’ (SRS) (anchored at 0 with ‘no sensation’ and 100 with ‘pain’), and the Numerical Pain Rating Scale (NRS) (anchored at 0 with ‘no pain’ and 100 with ‘most intense pain you can imagine’). Our analysis, which included a thorough description of individual and group-level response characteristics, showed that the SPARS has a fairly consistent curvilinear relationship to stimulus intensity and stable response characteristics across its range, and that the change in ratings between successive stimuli is not influenced by the direction of the change in stimulus intensity - i.e. there is no hysteresis.

Interestingly, however, our work raised doubt about the presence, and indeed the concept of pain threshold. That is, the curvilinear stimulus-response curve of the SPARS meant that the slope of the response curve flattened at stimulus intensities near the (presumed) pain threshold range, getting steeper as stimulus intensity was further increased or decreased. We found that the slope of the stimulus-response relationship for the NRS and SRS also flattened as stimulus intensities approached the pain threshold range, but ultimately ratings on both scales reached a floor or ceiling value, respectively (see Madden, et al. ^7^ for detailed discussion). Until now, pain assessment tools, by definition, presume a pain threshold, yet we propose that the flattening of the response curve in the middle stimulus range in the SPARS, and the lower end of the other scales, reflects a zone of uncertainty about whether a stimulus should be considered painful or non-painful.

Here, we aimed to clarify the characteristics of this apparent zone of uncertainty by undertaking novel exploratory analyses on data from each of two experiments ^7^ to formally assess the range of stimulus intensities across which the distribution of a participant’s repeated ratings of each intensity included pain threshold. We used SPARS data from both experiments, and NRS and SRS data from the second experiment to verify the SPARS findings.

## Materials & Methods

The two experiments that provided the data for the current analysis have previously been described in full in Madden, et al. ^7^. Here, we present only the essential details of the procedures and our analysis approach.

### Participants

We recruited convenience samples of healthy volunteers between 18 and 65 years of age. Volunteers were eligible if they: spoke English; had no history of sensory deficits or chronic pain (defined as daily pain for three months or longer^10^); did not have pain at the time of testing; were not using medication that might affect skin sensitivity, pain, or healing; did not have an elevated risk of laser-induced skin damage (e.g. thinned skin); and had not been diagnosed with diabetes mellitus, or a neurological, peripheral vascular, or psychiatric disease. Participants gave written informed consent and were compensated AU$20/hour. The study conformed to the Declaration of Helsinki, and was approved by the University of South Australia’s human research ethics committee (protocols 31880 and 36008).

### Stimulus

Cutaneous heat stimuli were applied in single trials using a neodymmium-doped yttrium aluminium perovskite laser (Nd:YAP; Stimul 1340, Deka, Italy), which is thought to selectively activate temperature-responsive neurons that terminate in the most superficial layers of the skin^11^. Stimulus location was slightly shifted for successive trials to prevent skin damage and progressive changes in sensitivity ^12,13^, consistent with recommendations on safe use of experimental laser stimuli ^14^.

### Procedures and Outcomes

In Experiment 1, participants received laser stimuli (spot diameter of 4mm, pulse duration 6ms, inter-stimulus interval ~15sec) to the skin of their backs whilst lying prone. They were shown a visual representation of the SPARS and provided verbal ratings for eight trials at each of 13 energy levels (1.00J to 4.00J, at 0.25J intervals) during four blocks of 26 pre-randomised trials (104 trials in total). Ratings were not restricted to whole numbers. For safety, trials of energy >3.50J were omitted from the sequence of trials when participants showed skin reddening or reported pain greater than +30, at the discretion of the investigator, as per ethical approval requirement.

In Experiment 2, participants received laser stimuli (spot diameter of 6mm, pulse duration 5ms, inter-stimulus interval ~30 sec) to the dorsal left forearm whilst sitting. This procedure was divided into three blocks, with participants using a different scale to provide their ratings for each block. The three scales were: the SPARS, a conventional NRS anchored at the extreme left with 0, ‘no pain’ and at the extreme right with 100, ‘most intense pain you can imagine’, and a numerical Sensation Rating Scale (SRS) anchored at extreme left with 0, ‘no sensation’ and at the extreme right with 100, ‘pain’. Scales were presented on a computer monitor and participants provided ratings using the computer mouse (Affect4.0 software ^15^). Note that the lower extreme of the NRS (‘no pain’) and the upper end of the SRS (‘pain’) overlap at the transition between non-painful and painful stimulus perception. Each block was divided into four stimulus runs, broken by short breaks, with three trials per intensity (3 x 9 intensities = 27 trials) delivered in randomised order during each run. Block order was counterbalanced across participants. Note that Experiment 2 differed from Experiment 1 in that the range of stimulus intensities delivered to each participant was individually determined using a calibration procedure (see Madden, et al. ^9^ for details), so the intensity range could span 1.5-3.5J, 1.75-3.75J, 2-4J, 2.25-4.25J or 2.50-4.50J with a 0.25J step between successive intensities. Note, too, that participants were not given criteria to use when judging a stimulus as painful or non-painful. Rather, they were allowed to freely determine what constituted ‘painful’ and ‘non-painful’ for them, in order to maximise ecological validity.

### Data handling and analysis

Data tidying and analyses were done in R v3.5.1 (The R Foundation for Statistical Computing), in RStudio v1.1.453 (RStudio Inc). We used the packages magrittr ^16^, tidyverse ^17^, patchwork ^18^, boot ^19,20^ and skimr ^21^. The scripts for data cleaning and analysis and their outputs are publicly available at https://zivahub.uct.ac.za/s/c6c12136c0a632698495 and as supplementary materials for this manuscript (Scripts S1 to S4). Data files are available on request only because participants did not consent to public release of their data. Requests for access to the data will be considered on a case-by-case basis and data made available subject to Human Research Ethics Committee approval. Requests can be directed to VJM. All analyses are exploratory in nature.

For consistency across scales during data analysis, we shifted the SRS scale that ranged from 0 (‘no sensation’) to 100 (‘pain’) so that it ranged from −100 (‘no sensation’) to 0 (‘pain’). Consequently, for all three scales, 0 identified the theoretical value of threshold. However, it should be noted that only the SPARS data directly address the width of the pain threshold, whereas data from the NRS and SRS shed light only on the range of intensities that are not clearly painful (NRS) or likely to have been painful (SRS). Participants in Experiment 2 had received stimuli across a range of different absolute intensities; therefore, intensity was recoded into a rank (of 1-9) to facilitate comparison between participants.

#### Analysis for question 1: How wide is the zone of uncertainty?

Initial visualisation of individual trial-by-trial SPARS, NRS and SRS ratings at each intensity suggested a reasonably wide zone of uncertainty for most participants on all scales (Supplement 1 and 2: Data at the level of the individual - Scatter plots). To explore the range this uncertainty, we visualised the uncertainty in intensity ratings by generating bootstrapped 95% confidence intervals (95% CI; resamples: 10000, interval type: basic) of the Tukey Trimean of ratings for each individual, at each stimulus intensity. The Tukey trimean is a robust measure of central tendency that, unlike a median, takes the spread of the data into account. It was chosen because it had been found to be the most suitable measure of central tendency for the current data ^7, 22^. We plotted these intervals to examine whether the uncertainty estimate for each individual, at each stimulus intensity, spanned 0 (hypothetical threshold) for each of the three scales. To explore the effect of examining data at the level of the group, rather than at the level of individual participants, we calculated the Tukey trimean for each individual, at each stimulus intensity, and then calculated the mean and bootstrapped 95% CI for the group at each stimulus intensity. As with the data for individual participants, the group-level 95% CI were then plotted over the stimulus range to examine whether the uncertainty estimates spanned 0 for each scale. For the individual and group data, we estimated the width of the zone of rating uncertainty by examining the range of stimulus intensities for which the 95% CI included 0.

To formalise our interpretation of the 95% CI plots, we used the binomial test to assess whether the scatter of ratings for each individual, at each stimulus intensity, were biased away from 0 (hypothetical threshold). The binomial test is used when there are two possible outcomes, and assesses whether the distribution of the observed data deviates from a distribution that would theoretically be expected to occur by chance. This analysis was conducted for the SPARS and repeated on the NRS because the NRS is a widely accepted pain rating scale, and on the SRS as a first exploration of this question on a scale of non-painful sensation.

We dichotomised the rating data at each stimulus intensity as follows: if a participant reported pain (i.e. SPARS rating > 0, NRS rating > 0, or SRS rating = 0) a trial was considered ‘positive’, and if a participant reported a non-painful percept (i.e. SPARS rating < 0, NRS rating = 0, or SRS rating < 0) a trial was considered ‘negative’. In Experiment 1, participants were not allowed to select a SPARS rating of 0, so all ratings were easily categorised. In Experiment 2, participants could select a SPARS rating of 0. For the purposes of this analysis, we deemed SPARS ratings of 0 to be uninformative, and so we excluded ratings of 0 from the analysis (we report the number of 0 ratings per participant below). In addition, ratings on the NRS were dichotomized for a second analysis informed by the previous report that individuals use the first 15 points of a 0 to 100 NRS to record non-painful stimuli ^23^. On the basis of this, we coded NRS ratings ≤ 15 as ‘negative’, and ratings >15 as ‘positive’. We repeated the SRS analysis using a similar 15-point window (SRS ratings ≤ −15 as ‘negative’, and ratings > −15 as ‘positive’).

We modelled the data using the binomial test assuming a 50% probability of ‘success’ (positive rating arbitrarily chosen as success) (please see Supplement 3: Question for a detailed explanation). Because ratings on the SPARS can range from −50 to +50, we analysed the SPARS data using a two-tailed p-value to reflect that the distribution may shift to the left or right of the theoretical distribution. However, because the NRS and SRS have a floor rating of 0 (‘no pain’) and a ceiling rating of 0 (‘pain’) respectively, the change in rating can only occur in one direction away from 0, so we analysed the NRS and SRS data using one-tailed p-values. For all tests, significance was assessed at the a = 0.05 level. No family-wide corrections were made for multiple comparisons because this was an exploratory analysis. When the observed data were consistent with the expected distribution (p > 0.05), the result can be interpreted as indicating that the distribution of ‘painful’ and ‘non-painful’ judgements (at that intensity) does not differ from that predicted by chance – or, in other words, that intensity was not clearly outside the zone of uncertainty.

#### Analysis for question 2: Did the trial-by-trial change in intensity influence ratings?

Discussion of our findings raised the question of whether ratings had been influenced by the magnitude of the change in stimulus intensity between the preceding trial and the trial being rated. Therefore, we performed a visual analysis of the relationship between the difference in intensity between two successive stimuli and the rating of the second stimulus, using colour coding. First, we plotted the ratings at each stimulus intensity, using a two-colour coding system to distinguish ratings that had been given after the most extreme changes in stimulus intensity from ratings that had been given after the smallest changes in stimulus intensity. The absolute difference in stimulus intensity between a stimulus intensity and the preceding stimulus was used. If the rating of a certain stimulus intensity differed according to how different that stimulus was from the preceding one, we expected a visual pattern with a consistent distribution of ratings depending on whether there was a maximal intensity change or minimum intensity change. We hypothesised that ratings after the largest intensity changes would consistently be more extreme than ratings after the smallest intensity changes. Second, we used a gradient of colours to code each data point, at each stimulus intensity, according to the magnitude of the change in stimulus intensity between that trial and the preceding trial (i.e. colour indicates the difference in stimulus intensity between a given stimulus intensity and the preceding stimulus). If the rating of a certain stimulus intensity differed systematically according to how different that stimulus intensity was from the preceding one, we expected to see the colour gradient reproduced in the distribution of the data points in the plots. This analysis was conducted for the SPARS, NRS, and SRS data.

## Results

For Experiment 1, nineteen adults participated (11 female; 18-31 years old). Two participants showed marked skin reddening, leading to omission of high-intensity trials for those participants and missing response data – for one participant, at 4J only, and for another participant, at 3.5J and 3.75J. For Experiment 2, seven adults participated (5 female; 21-30 years old). One participant received stimuli within the range of 1.75J-3.75J, three participants received 2.25J-4.25J, and three participants received 2.50J-4.50J. Due to a software error, all SRS rating data for one participant were lost. The remaining data for that participant were included in the analysis. In Experiment 2, 21 out of 752 (3%) SPARS trials, across all participants, were rated 0 and excluded from the binomial test analysis.

### Results for question 1: How wide is the pain threshold?

The preliminary assessment of rating uncertainty using bootstrap 95% CI for individual-level and group-level data are shown in Figs S1 (individual-level SPARS), S2 (group-level SPARS), and S3 (individual- and group-level NRS and SRS). The ‘zone of uncertainty’ on the SPARS varied widely across participants, with its width spanning between 1 (ID05, ID08) and 8 (ID03, ID13, ID14) of the 13 different stimulus intensities in Experiment 1 (Fig. S1A) and 1 (ID05) to 8 (ID03) of 9 stimulus intensities in Experiment 2 (Fig. S1B). Moreover, within each individual, there was not necessarily a continuous block of intervals spanning 0. The magnitude of the inter- and intra-individual variation was obscured when looking at group-level data (Fig. S2), which shows a continuous zone spanning 8 of 13 intensities in Experiment 1 (Fig. S2A) and 5 of 9 intensities in Experiment 2 (Fig. S2B). Similarly, the NRS plots for each participant (Fig. S3A) show variable and fragmented ranges of stimulus intensities where the 95% CI included 0, varying between 0 (ID07) and 7 (ID01, ID03, ID06) of the 9 intensities, depending on the individual. Also, like the SPARS, the group-level NRS analysis (Fig. S3B) obscured the inter- and intra-individual variability, giving a false impression of consistency. Finally, the SRS data repeat the SPARS and NRS findings of high inter-individual variability (Fig. S3C), which is hidden when looking at the group-level data in isolation (Fig. S3D).

The results of the binomial test results are shown in Figs 2 (individual-level SPARS data) and 3 (individual-level NRS and SRS data). This more formal analysis confirms our preliminary findings that used 95% CI by showing a broad zone of uncertainty that varies substantially between individuals. In Experiment 1 (Fig. 2A), this zone spanned very little (ID15) to almost all (ID01) of the SPARS’s stimulus range and, although the distribution of intensity ratings tended not to differ from the expected distribution in the central range of stimulus intensities, there was evidence of uncertainty at non-adjacent intensities for most participants (indicated by columns of light-shaded dots between columns of dark-shaded dots). Indeed, for participants ID04 and ID05, there seemed to be no consistent stimulus-response relationship. In Experiment 2 (Fig. 2B), the SPARS’s zone of uncertainty showed similar inter-individual variability. The binomial test results for the NRS and SRS are shown in Fig. 3. Analysis of the NRS data according to a strict interpretation of any rating > 0 as indicating a painful percept (Fig. 3A) indicates that the observed distribution of ratings was consistent with the expected distribution for 0 (ID07) to 5 (ID01, ID06) of the 9 stimulus intensities, whereas analysis of the NRS data according to how existing literature suggests participants actually use it – considering the first 15 points on a 0-100 NRS to indicate a non-painful percept (Fig. 3B) – indicates that the difference in rating distributions for 5 (ID07) to 9 (ID01, ID03) of 9 intensities did not differ from the expected distribution. Unlike for the SPARS, where there frequently were non-contiguous blocks of rating that were not biased away from 0, the results for the NRS followed a consistent pattern of low intensity stimuli not deviating from the expected distribution, but a bias towards ratings being consistently greater than 0 at higher stimulus intensities. The SRS data are reported in Fig. 3C (using a < 0 threshold) and 3D (using a < −15 threshold) and show that, unlike the SPARS and NRS data, the majority of SRS ratings were biased away from the threshold, even when using the expanded definition of threshold, and the bias tended to increase as stimulus intensity decreased.

**Figure 2:**
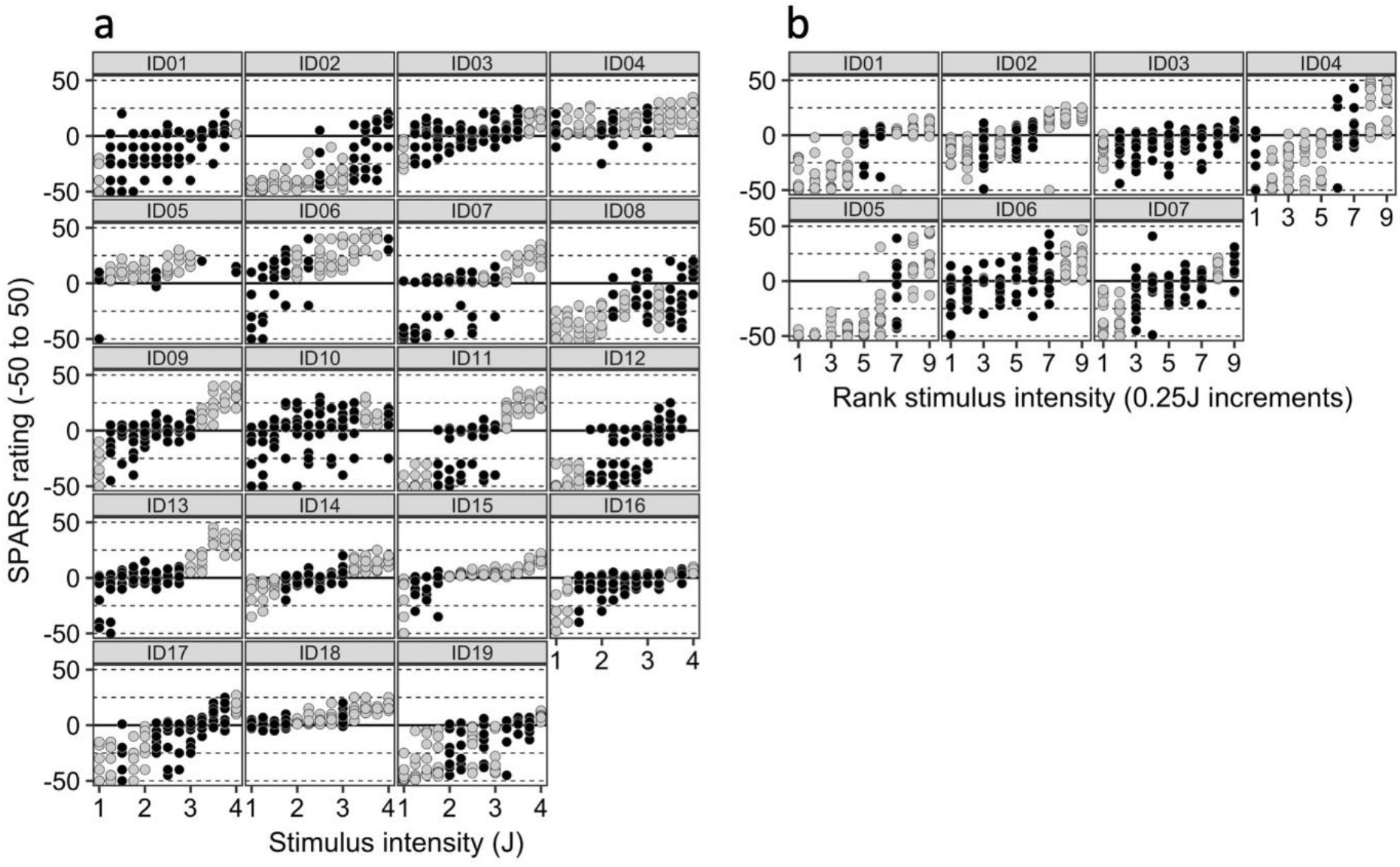
SPARS data: Results of the binomial test of positive/negative ratings distribution at each intensity, where probability of success = 0.5 (where ‘success’ is arbitrarily defined as a positive trial; see Methods for details and rationale); alpha = 0.05. The bootstrap procedure used 10 000 resamples. The solid line indicates the presumed pain threshold. Dark circles indicate that the distribution of ratings did not deviate significantly from the expected distribution (i.e. null hypothesis upheld). P-values are two-tailed. A: data from Experiment 1. B: data from Experiment 2.

**Figure 3:**
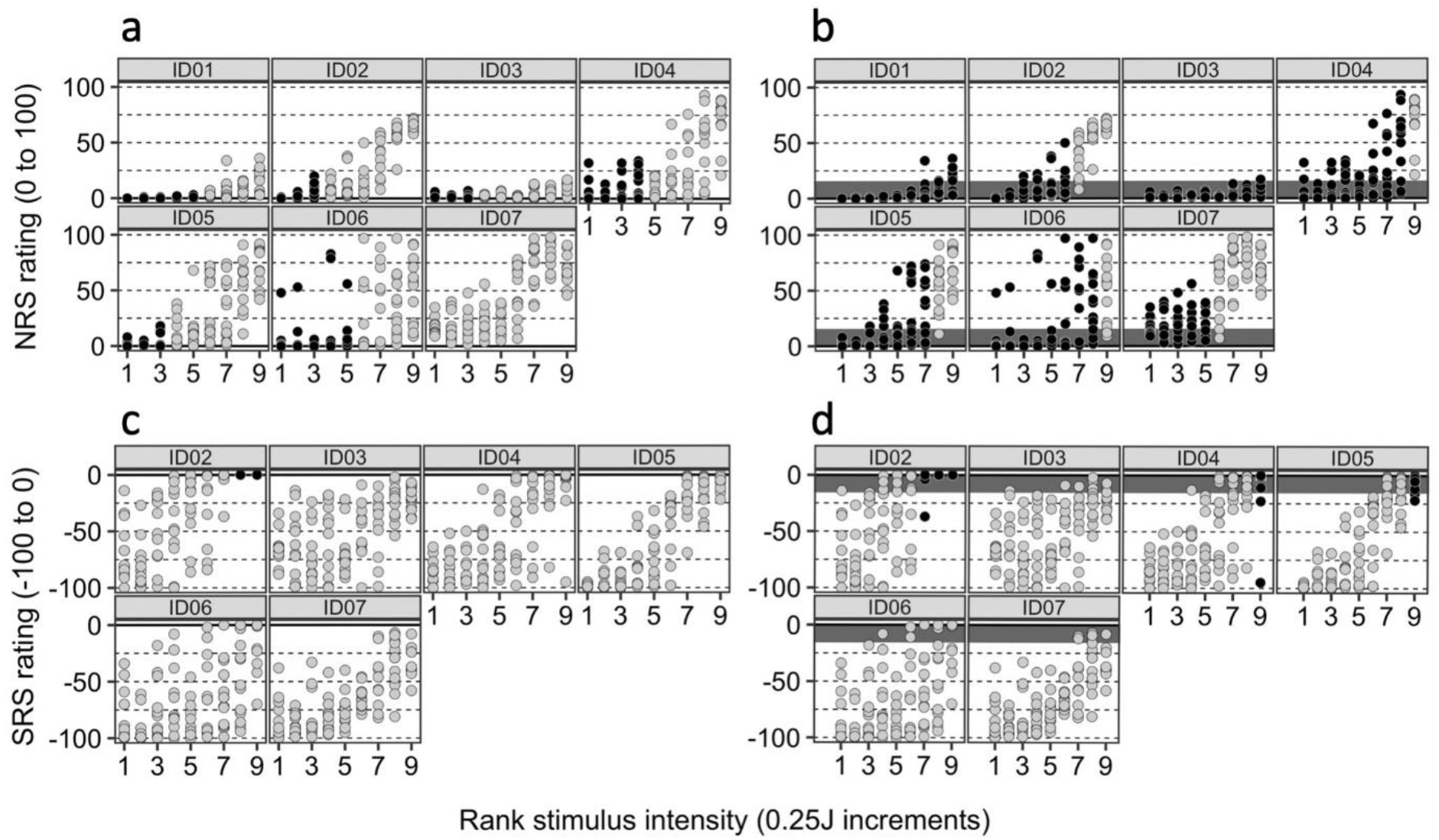
NRS and SRS data: Results of the binomial test of positive/negative ratings distribution at each intensity, where probability of success = 0.5 (where ‘success’ is arbitrarily defined as a positive trial); alpha = 0.05. The bootstrap procedure used 10 000 resamples. The solid line indicates the presumed pain threshold. Dark circles indicate that the distribution of ratings did not deviate significantly from the expected distribution (i.e. null hypothesis upheld). P-values are one-tailed. A: NRS data with rating > 0 considered ‘positive’. B: NRS data with rating ≥ 15 considered ‘positive’; dark-shaded region indicates range of NRS 0-15. C: SRS data with rating < 0 considered ‘negative’. D: SRS data with rating > 15 considered ‘positive’; dark-shaded region indicates SRS range of −15-0.

### Results for question 2: Did the jump in intensity influence ratings?

Figure 4 (SPARS) and Fig. 5 (NRS and SRS) depict the results of the graphical analysis of the relationship between the difference in intensity between two successive stimuli and the rating of the second stimulus. We had hypothesised that ratings given after the largest intensity changes would be more extreme (more negative or positive) than ratings after the smallest intensity changes. No clear pattern of rating can be observed for any of the participants for any of the scales. Similarly, no clear pattern of ratings could be discerned when plotting all ratings at a given intensity conditioned on the magnitude of the intensity change from the preceding stimulus (Fig. S4 (SPARS) and S5 (NRS and SRS).

**Figure 4:**
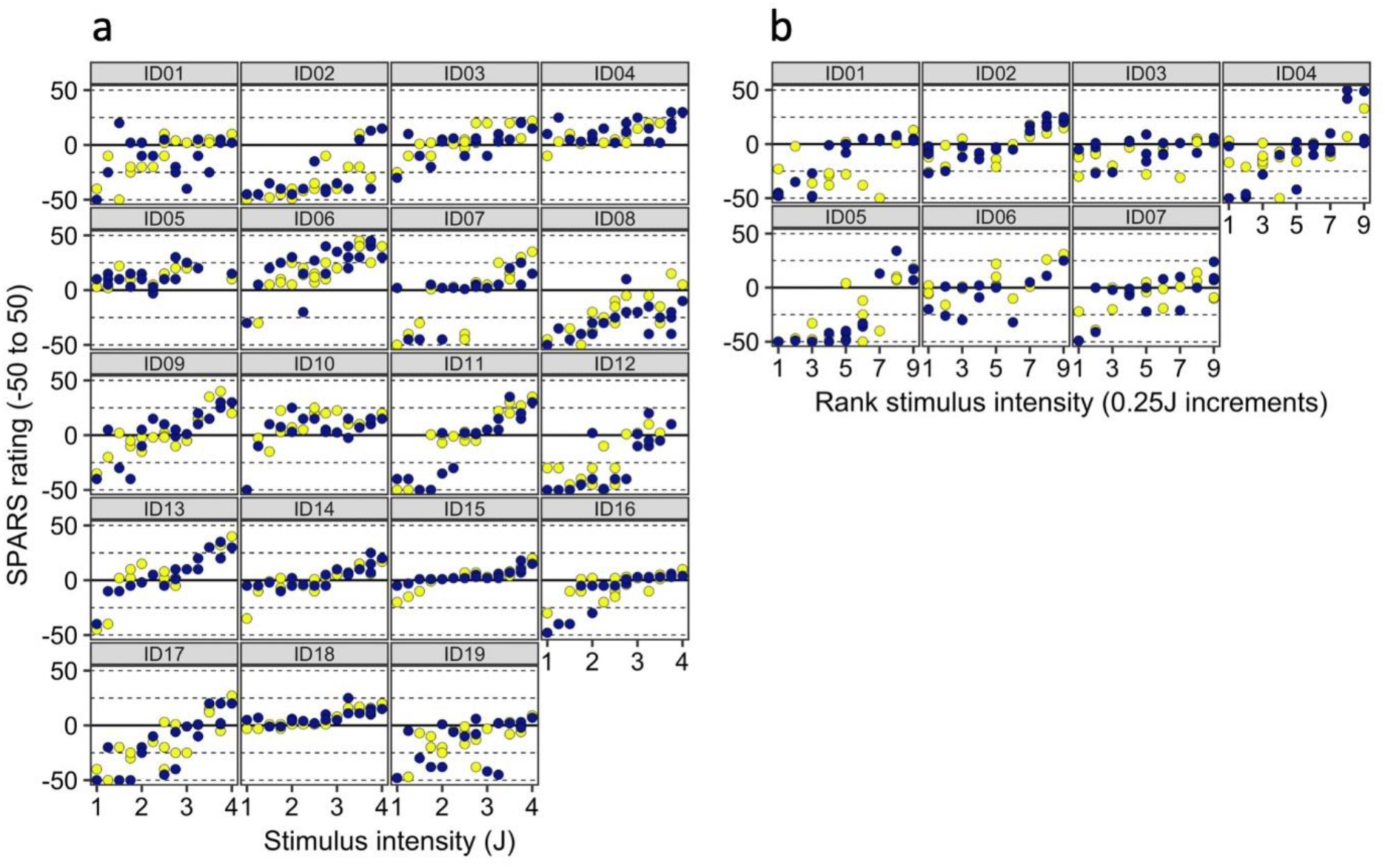
Results for question 2: Did the jump in intensity influence ratings? Individual scatterplots of SPARS ratings, including only the trials of each intensity that followed the maximum and minimum changes in stimulus intensity. Purple: trial followed the maximum change in intensity; yellow: trial followed the minimum change in intensity. Note that multiple dots of the same colour reflect that multiple trials followed the same maximal/minimal change in intensity. The absolute value of the change in intensity was used. A: data from Experiment 1. B: data from Experiment 2.

**Figure 5:**
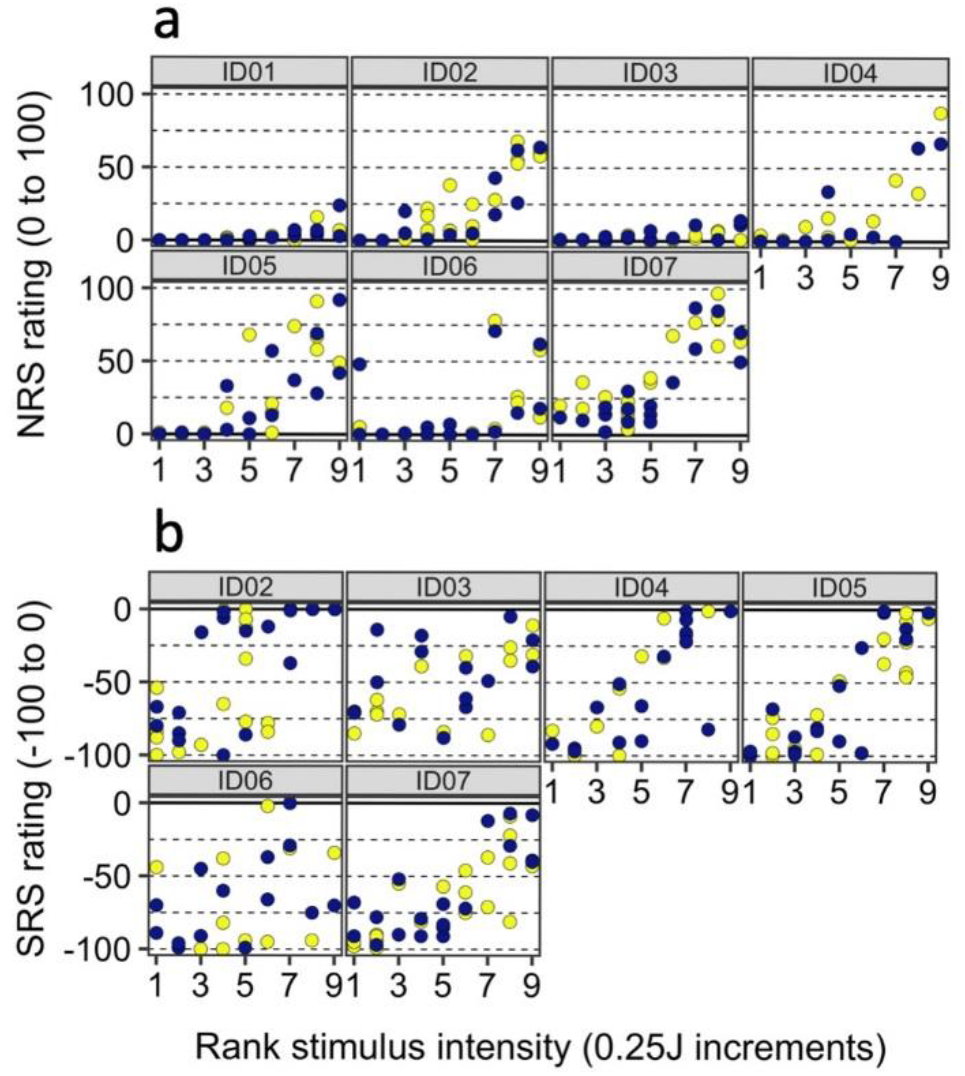
Results for question 2: Did the jump in intensity influence ratings? Individual scatterplots of (A) NRS and (B) SRS ratings, including only the trials of each intensity that followed the maximum and minimum changes in stimulus intensity. Purple: trial followed the maximum change in intensity; yellow: trial followed the minimum change in intensity. Note that multiple dots of the same colour reflect that multiple trials followed the same maximal/minimal change in intensity. The absolute value of the change in intensity was used.

## Discussion

This exploratory analysis followed our previous observation that the SPARS, NRS and SRS showed flattening of the stimulus-response curve as stimulus intensity approached what each scale predetermines to be pain threshold. In other words, participants seemed to have provided contradictory ratings of pain and no pain on different trials of the same stimulus intensities, with no significant bias. Here, we aimed to determine and compare the width of this ‘zone of uncertainty’ or pain threshold for the Sensation and Pain Rating Scale (SPARS), using data from the Numerical Rating Scale (NRS) to verify the findings on the SPARS. We also explored the distribution of ratings on the Sensation Rating Scale (SRS). We used the binomial test to formally identify intensities at which the distribution of ratings was biased away from 0 (hypothetical pain threshold), and then plotted these results for each individual so as to visually examine variability within and between individuals (Fig. 2). This revealed considerable intra-individual variability in ratings of a given stimulus intensity, and this variability in ratings differed between individuals. Across individuals, there was (inter-individual) variability in the width of the zone of uncertainty on the SPARS. The NRS also showed a wide range of stimulus intensities that were not clearly painful, whereas the SRS showed a non-existent to very small range of intensities that were possibly painful. When visualised using the 95% confidence intervals of the Tukey trimean (Fig. S2), even the group-level data showed a wide zone of uncertainty for the SPARS, including 8 (62%) of 13 intensities or 5 (56%) of 9 intensities. For the NRS, the group-level data showed that 6 (67%) of 9 intensities were not clearly painful; for the SRS, the group-level data showed that 2 (22%) of the 9 intensities were possibly painful.

It is surprising that most participants in the current study displayed uncertainty about painfulness at several different stimulus intensities, considering that the term ‘pain threshold’ suggests a clear boundary between non-painful and painful percept. One early description of a pain threshold identification procedure speaks of crossing the pain threshold as a “sharply defined experience” and reported less than 12% intra-individual variation in thresholds estimated by daily measurements from three people over the course of about a year^24^ – a sharp contrast to the high intra-individual variability seen in the current results (Fig. 2). Procedural differences may account for this: their testing procedure used an ascending series of 3-second phasic stimulations until the participant reported pain. That ‘method of limits’ is similar to the widely-used ramped stimulus approach to measuring the heat pain threshold with a tonic stimulus that increases in intensity until the participant reports pain^25^. In both ascending procedures, the explicit goal is to establish a point of transition, and that is the task the participant is given for every trial – it is a stimulus-dependent method^26^. In contrast, our experiments used an array of randomly-ordered phasic stimuli and participants were tasked with reporting the percept elicited by each one in isolation – a response-dependent method. Our approach shares features with the multiple staircase approach to calibrating stimulus intensity to a perceptual threshold^26^, which is also a response-dependent method; a method that may be better suited to calibrating stimuli that will be repeatedly delivered during an experimental procedure.

The assumption that a clear boundary or threshold exists between ‘non-painful’ and ‘painful’ percepts may have arisen from an extrapolation of the concept of a stimulus detection threshold. Certainly, sensory detection thresholds do seem to be a more reliable phenomenon than pain thresholds^25,27^, although the stimulus-response curve has previously been shown to be flatter close to the sensory detection threshold than at intensities that are consistently detected^28^. That is, as the stimulus intensity approaches the absolute threshold, there is a breakdown of the Weber-Fechner law^29–31^, such that the just noticeable difference is influenced more by the levels of the absolute threshold than by the initial stimulus strength. Perhaps the high variability in judgements about pain relative to the moderate variability in judgements about sensation is due to differences in how sensation and pain are generated. While both sensation and pain require higher centres, the relative contributions of afferent signalling and higher processing to the generation of sensation and pain may differ, resulting in a closer relationship of stimulus to response in sensation than in pain. Interestingly, while most participants in the current data showed uncertainty at several intensities, a small number did seem to have a clear boundary (see Fig. S1A ID06, Fig. S1B ID05). These observations point to the novelty of an analytical approach that allows meaningful quantification of the zone of uncertainty - perhaps the relative influence of afferent input to higher centre processing on pain differs between individuals; perhaps interventions modulate the width of the zone of uncertainty and this has clinical utility; perhaps biological (e.g. genetic), psychological (e.g. catastrophising, psychological flexibility), or contextual (e.g. social environment) factors relate to width of the zone; perhaps the width of the zone has predictive utility for outcome, response to treatment or risk of poor recovery. All these proposals are clearly speculative, but they seem worthy of further investigation.

Regardless of the explanation for the intra-individual variation in the current study, it presents a problem for experiments that aim to calibrate stimuli to the pain threshold; in fact, based on our data, we suggest that the ‘pain threshold’ may be more parsimoniously considered to be a ‘zone of uncertainty’ that varies in width and clarity by individual (and, likely, by modality and testing approach), and that may reflect a gradual shift in the relative contributions of nociceptive and non-nociceptive contributions to the percept.

The amount of intra-individual variability is also surprising in light of the fact that Nd:YAP laser provides a carefully calibrated stimulus ^11^. Although it is well established that nociception and pain do not share an isomorphic relationship ^32^ either within or outside the laboratory, this amount of intra-individual rating variation still seems remarkable within the highly controlled context of the laboratory. While some intra-individual variability is to be expected in data on pain, the current data on the Sensation Rating Scale as well as the SPARS may be the first demonstration of such a variable relationship between non-painful nociception and percept. ^33^intra-individual variability was also high on the SRS, yet the zone of uncertainty was considerably more narrow on the SRS than on the NRS or SPARS (Fig. 3 cf. Fig. 2). This may reflect a cueing effect: most of the SRS covers the non-painful range of intensities, and participants may be reluctant to commit to using the extreme upper anchor of ‘pain’. It is possible that presenting a ‘non-painful’ scale range provides a strong enough cue that the event will be non-painful to raise the likelihood that the actual percept will be of a non-painful event. This process could be framed in Bayesian terms (which we discuss in greater detail below) and/or in terms of the influence of conscious expectations^34,35^.

The high intra-individual variability in ratings on both the SPARS and the NRS has four main implications. The first implication is for studies that calibrate stimulus intensities to each individual participant. With this high rating variance, it is likely that a stimulation intensity chosen on the basis of a few ratings at the start of an experimental procedure will not reflect the participant’s percept upon every delivery of that same stimulation intensity. We suggest that asking participants to report the percept on every stimulation trial would help to verify the match between the rating given during calibration and the percept elicited by each stimulation trial – although this would be the case only if the scale used for ratings is able to capture a shift of the percept across the pain threshold. A second implication of these data relates to adequate powering of experimental studies: such wide variance (low precision) in ratings stands to inflate the sample sizes required to demonstrate differences between conditions or groups, with obvious ramifications for study burden, funding requirements and the progress of robust science. Indeed, the high intra- and inter-individual rating variance means that small samples sizes and few replicates per individual increases the risk of random error, potentially leading to spurious findings that misrepresent the direction and magnitude of the effect of an intervention^36,37^. Multilevel analysis approaches that allow for within-subject comparisons nested within group trends can help to control for inter-individual variability, but intra-individual variability will remain problematic. A third implication is for the profiling of clinical patients using subjective ratings of experimental stimuli. Recently, calls have been made to consider matching different treatment algorithms to groups of patients defined by their responses to psychophysical phenotyping procedures^38–40^. The current data suggest that caution may be indicated in the case of phenotyping procedures that use experimental stimuli: many repetitions may be required in order to ascertain the true range of an individual’s response to any one stimulus intensity. The final, and perhaps the most intriguing, implication is that the variability in how an individual perceives a highly controlled stimulus, even in a highly controlled setting, could reflect an optimal flexibility in the system. A Bayesian inferential – rather than a stimulus-response – account for pain suggests that a perceptual system that is malleable to fluctuations in external and internal information is useful. Such a system would allow for optimisation of the percept to the moment-by-moment context – an optimisation that must inevitably occur on the basis of incomplete information ^41^. Bayesian theory proposes that incoming input is integrated with existing priors to inform the perceptual experience ^35, 42^. We used a simple graphical approach to examine whether the ratings given in the current data were influenced by the difference between the preceding stimulus and the current stimulus (as a crude proxy for mismatch between the prior and the input), and found no evidence for such an influence in our data (Fig. 3). However, Bayesian theory proposes that the consistency of information determines how heavily it is weighted in the perceptual inference process ^43^, a concept captured by the principles that govern influence of neural networks on each other (see ^44^ for pain-specific discussion). It is possible that, in the current data, there was enough variability in the neural activation elicited even by laser stimuli of identical intensities to limit the influence of the change in stimulus intensity on the percept. Importantly, known variations in the spatial distribution of warmth receptor fields would likely contribute to this variability, because the laser stimulus was slightly shifted between trials results ^33^. Alternatively, perhaps other influences are sufficient to introduce noise despite stability in the nociceptive input - possible candidates include moment-by-moment changes in cognition or emotion, expectation, or even cross-modal modulation by minute visual or olfactory influences^13,34,35,45–51^. These ideas remain speculative because the current data were obtained during experiments designed according to a ‘stimulus-response’ model rather than a Bayesian inference model for pain. Nevertheless, the idea that the variability of an individual’s ratings could reflect the flexibility of his/her perceptual processing – and therefore, a beneficial feature of the system – is reminiscent of recent discussions of the potential benefits of psychological flexibility in relation to persistent pain ^52^. That healthy controls seem to have more variable ‘pain thresholds’ than people who have persistent pain^27^ would seem consistent with this idea. This raises the intriguing possibility that the width and clarity of the zone of uncertainty, as distinct from the average stimulus-response profile, represents a useful biomarker of risk of persistent pain, actual or predicted response to treatment.

The inter-individual variability in scale use suggested by the plots of the current data is also worthy of comment. Here, too, the distribution of warmth receptor fields differs between individuals, which likely contributes to the inter-individual variability seen in the results ^33^. The mismatch between the group-level plots and the individual plots in Figs S1, S2 and S3 shows that, in this data set, grouping of data from participants whose responses vary widely actually obscures the inter-individual variability in scale use, giving the artificial impression that the zone of uncertainty is consistent. Inter-individual variability is not a new discovery (e.g. see ^29^), yet it is frequently neglected in the analysis of experimental data. Although analysing data at a group level has a role in some studies, many modern analytical techniques allow for nesting of individuals within groups, and the current data suggest that such an approach would be warranted when analysing experimental self-rating data on painful and non-painful percepts. At the very least, a visual examination of inter-individual variability should be considered so as to determine the need for a multilevel analysis. Indeed, patients with pain typically seek clinical treatment for and relief of *their own* pain, not for the pain of a group. Considering that experimental studies on pain lay the groundwork for clinical studies that inform clinical treatments, experimental researchers would do well to design analyses to shed light on individual changes in percept as well as changes at the group level.

The current analysis used data obtained in the experimental setting and may not reflect the clinical reality. We can therefore only tentatively speculate on the implications of our findings for clinical research. That not only the new SPARS, but also the widely accepted NRS, shows such large inter-individual variability in the experimental context suggests that clinical trials, too, ought to consider investigating change at the individual level as well as the ubiquitous ‘group effect’. Another addition that might lend value in clinical research would be the previously recommended inclusion of an outcome that assesses whether the change achieved is meaningful to each individual^53^. This would represent an advance from merely comparing the change achieved to known estimates of the magnitude of change that is meaningful to ‘the average individual’.

The current analysis has limitations that should be taken into account. First, this was an exploratory analysis of existing data. Second, the heating elicited by even a precisely calibrated laser stimulation may vary according to skin pigmentation, perfusion and other localised factors^14^. Therefore, this study needs to be replicated, and such replication would lend additional value if it used a different (although still carefully controlled) stimulus modality. Third, we intentionally refrained from conducting a full-scale multilevel modelling analysis that pooled data across individuals and intensities, because this analysis was exploratory. Such an analysis would be ‘pushing our data too hard’, but would be appropriate in a replication study designed *a priori* for multilevel modelling. Fourth, while intra-individual variability in ratings was formally tested with the binomial test, inter-individual variability was not formally tested, but is visible in the plots. Finally, we did not assess for an effect of stimulus order in the current analysis. This is because the initial analysis of these data^9, 22^ had previously verified that stimulus order had no effect on ratings in these data.

Data from the SPARS, conventional NRS and SRS show that participants are inconsistent when categorising stimuli of known intensities as painful or non-painful over a reasonably wide range of stimulus intensities, suggesting that the transition between painful and non-painful may occur in a ‘zone of uncertainty’ rather than at a clear boundary. The width of this zone of uncertainty differs between individuals and depending on the scale being used.

## Conclusions

The current findings have implications for experiment design, the size of sample that is required for adequate statistical powering of experiments, and the use of experimental stimuli for clinical phenotyping. Further, the intra-individual variability in responses may reflect a flexibility that could be useful for predicting risk of persistent pain or response to treatment.

