## Supplementary Figure 1 for "Rethinking pain threshold as a zone of uncertainty"

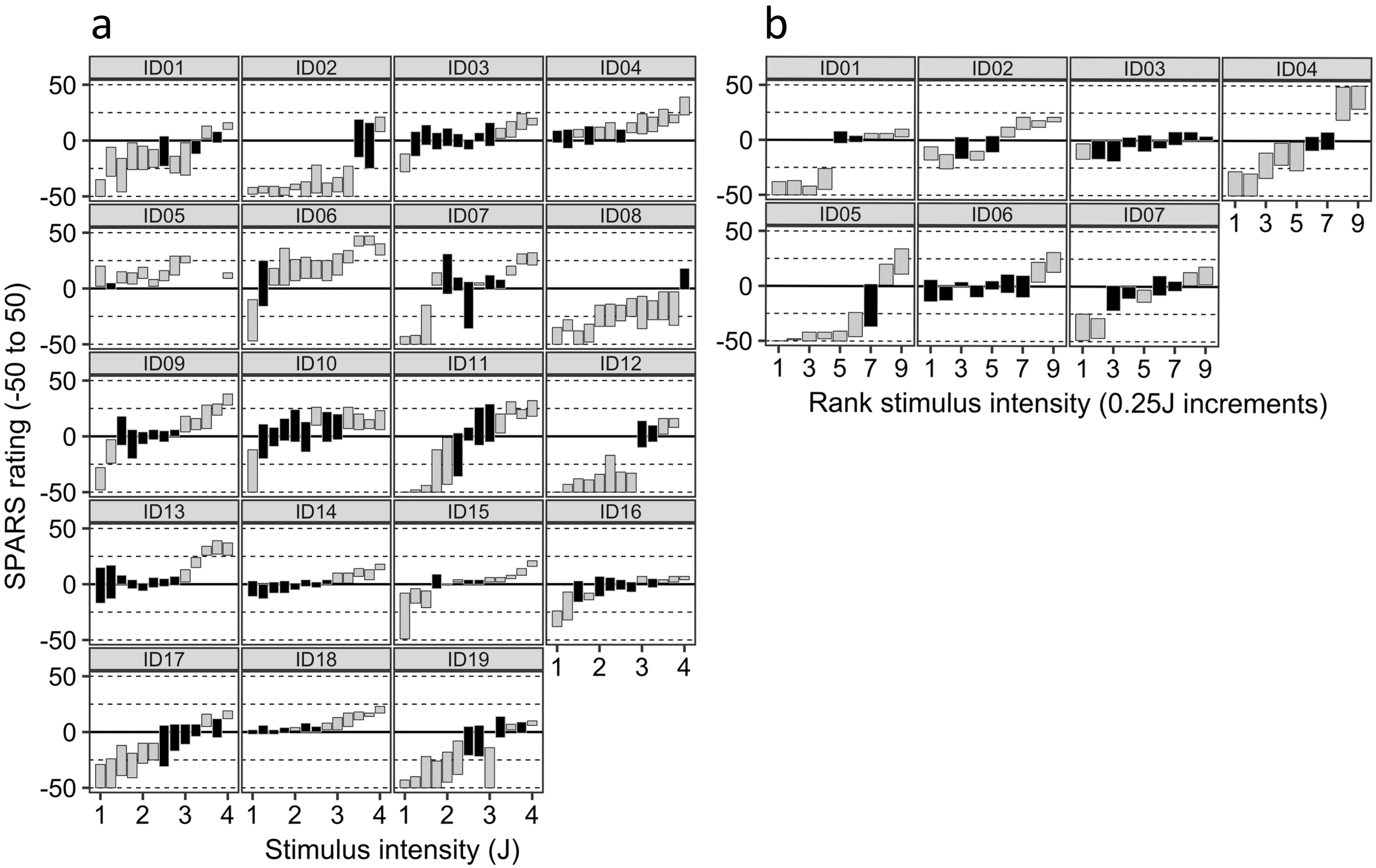


**Supplementary Figure 1:** Individual-level SPARS data from (a) Experiment 1 and (b) Experiment 2. Crossbar plots of bootstrapped 95% confidence interval of Tukey trimeans at each intensity. The bootstrap procedure used 10 000 resamples. The solid line indicates the presumed pain threshold. Grey shading of crossbars indicates that the 95% CI includes 0.
