## Supplementary Figure 3 for "Rethinking pain threshold as a zone of uncertainty"

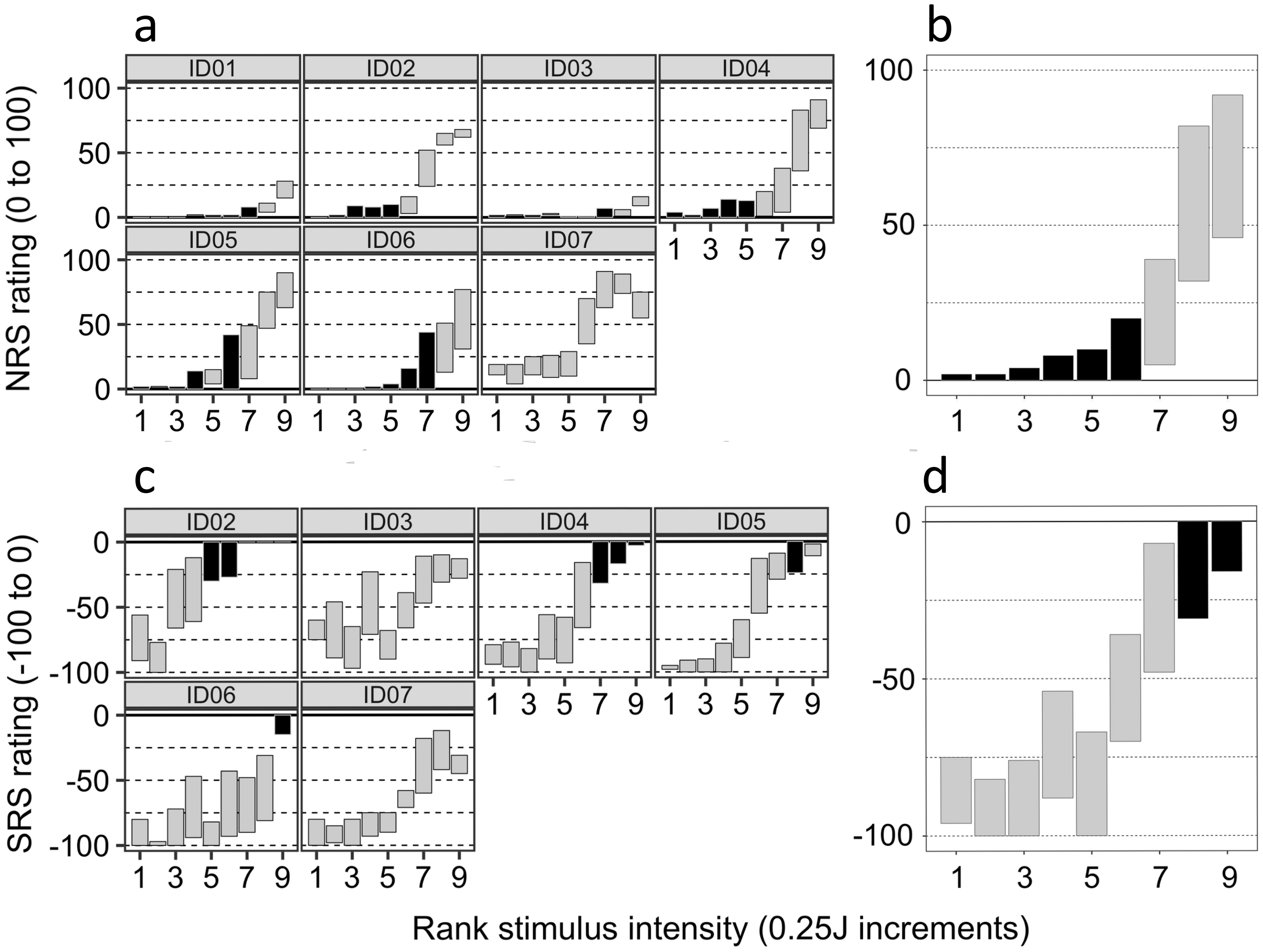


**Supplementary Figure 3:** NRS and SRS data from Experiment 2: crossbar plots of bootstrapped 95% confidence interval of Tukey trimeans at each intensity. (a) individual plots of NRS data, (b) group-level plot of NRS data, (c) individual plots of SRS data, (d) group-level plot of SRS data. The bootstrap procedure used 10 000 resamples. The solid line indicates the presumed pain threshold. Grey shading of crossbars indicates that the 95% CI includes 0.
