## Supplementary Figure 4 for "Rethinking pain threshold as a zone of uncertainty"

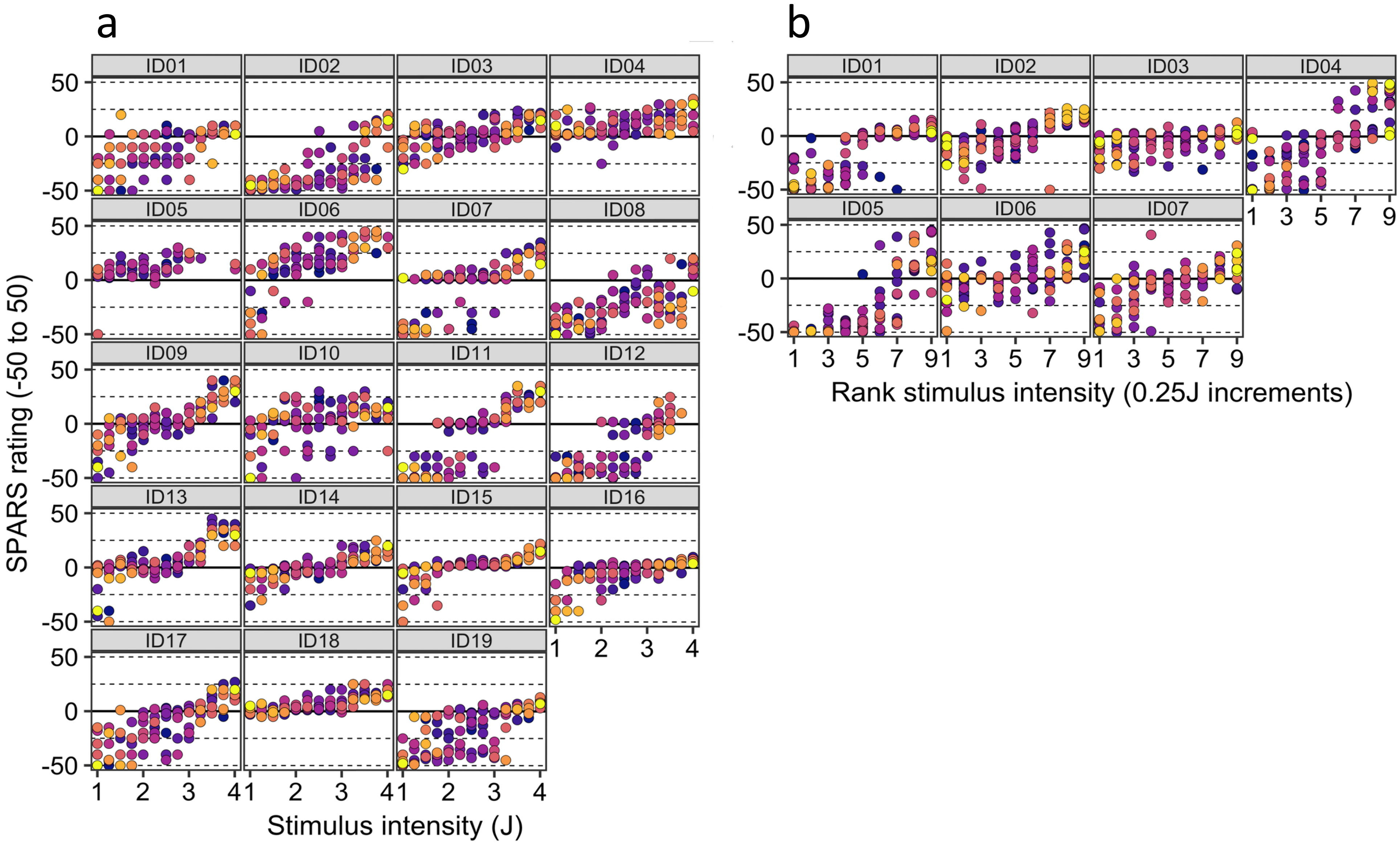


**Supplementary Figure 4:** Results for question 2: Did the jump in intensity influence ratings? Individual scatterplots of SPARS ratings, including all trials. Dots are coloured with a gradient according to the magnitude of the change in intensity that preceded that trial, with darker dot colour indicating a smaller change in intensity. Note that multiple dots of the same colour reflect that multiple trials followed the same maximal/minimal change in intensity. The absolute value of the change in intensity was used. a: data from Experiment 1. b: data from Experiment 2.
