## Supplementary File 1 for "Rethinking pain threshold as a zone of uncertainty"

SPARS A: Width of the pain threshold

*12 Jan 2019*

**Session information** **12**

---

#### Question

How wide is the pain threshold for participants taking part in the SPARS A trial?

To answer the question, we calculated the *Tukey Trimean* and bootstrapped 95% confidence interval (CI) for each individual, at each stimulus intensity. Next, we plotted these statistics to show the stimulus range over which each individual's CIs included zero on the SPARS (pain threshold).

To get an idea of the width of the stimulus range that included zero on the SPARS at the group level, we calculated the *Tukey trimean* for each individual, at each stimulus intensity, and then calculated the mean and bootstrapped 95% CI for the group at each stimulus intensity. These data were then plotted to show the stimulus range over which the group's CIs included zero (pain threshold).

The selection of the *tukey trimean* as the measure of central tendancy at the individual level was based on the analysis of central tendancy reported in the original description of the SPARS (Supplement_3.pdf). The *Tukey trimean* is defined as the weighted average of the distribution's median and its two quartiles, and is a robust measure of central tendancy that unlike a median, takes the spread of the data into account.

$$T_{mean} = \frac{1}{2}(Q_2 + \frac{Q_1 + Q_3}{2})$$

Where:

- $Q_1 = 25^{\text{th}}$ percentile

- $Q_2 = 50^{\text{th}}$ percentile (median)

- $Q_3 = 75^{\text{th}}$ percentile

```
# Define the tri_mean function
tri_mean <- function(x) {
    # Calculate quantiles
    q1 <- quantile(x, probs = 0.25, na.rm = TRUE)[[1]]
    q2 <- median(x, na.rm = TRUE)
    q3 <- quantile(x, probs = 0.75, na.rm = TRUE)[[1]]
    # Calculate trimean
    tri_mean <- (q2 + ((q1 + q3) / 2)) / 2
    # Round to a whole number
    tri_mean <- round(tri_mean)
    return(tri_mean)
}
```

**Note:** No inspection of block and stimulus order effects were undertaken because analysis of these factors in the original description of the SPARS revealed no order effects (Supplement_4.pdf).

The experimental protocol called for participants to be exposed to 13 stimuli, evenly spaced at 0.25J intervals over the range 1.00J to 4.00J. Each stimulus intensity was applied 8 times, giving a total of 104 exposures (trials). To prevent learning effects, the 104 trials were randomised across 4 experimental blocks (26 trials per block).

---

### Import and inspect data

```
# Import
data <- read_rds('data-cleaned/SPARS_A.rds')

# Inspect
glimpse(data)
```

```
## Observations: 1,927
## Variables: 6
## $ PID          <chr> "ID01", "ID01", "ID01", "ID01", "ID01", "ID01", "...
## $ block        <chr> "A", "A", "A", "A", "A", "A", "A", "A", "A", "A",...
## $ block_order  <dbl> 4, 4, 4, 4, 4, 4, 4, 4, 4, 4, 4, 4, 4, 4, 4, 4, 4...
## $ trial_number <dbl> 79, 80, 81, 82, 83, 84, 85, 86, 87, 88, 89, 90, 9...
## $ intensity    <dbl> 3.00, 2.25, 4.00, 3.25, 2.75, 2.25, 2.75, 4.00, 2...
## $ rating       <dbl> -40, -25, 10, 2, -10, -25, -20, 10, -25, -50, -25...
```

```
data %>%
    select(intensity, rating) %>%
    skim()
```

```
## Skim summary statistics
##  n obs: 1927
##  n variables: 2
##
## -- Variable type:numeric -----------------------------------------------------------
##   variable missing complete    n  mean    sd  p0    p25 p50   p75 p100
##  intensity       0     1927 1927  2.47  0.93   1   1.75 2.5  3.25    4
##     rating       0     1927 1927 -4.45 22.31 -50 -20    2   10      45
##      hist
```

```
##
##
```

---

### Data at the level of the individual

#### Bootstrapping procedure

```
# Nest data in preparation for bootstrapping at each stimulus intensity
data_boot <- data %>%
    group_by(PID, intensity) %>%
    nest()

# Define bootstrap function
boot_tri_mean <- function(d,i){
    tri_mean(d[i])
}

# Perform bootstrap
set.seed(123456789)
data_boot %<>%
        mutate(boot = map(.x = data,
                          ~ boot(data = .x$rating,
                                 statistic = boot_tri_mean,
                                 R = 10000, # For small sample size
                                 stype = 'i')))

# Remove NULL bootstrap row 56 (ID05, only one value = 20)
data_boot <- data_boot[-56, ]

# Extract CI from boot object
data_boot %<>%
    mutate(boot_ci = map(.x = boot,
                         ~ boot.ci(.x,
                                   type = 'basic')))

# Extract the data, giving original trimean and bootstrapped CI
data_boot %<>%
    mutate(tri_mean = map_dbl(.x = boot_ci,
                              ~ .x$t0),
           lower_ci = map_dbl(.x = boot_ci,
                              ~ .x$basic[[4]]),
           upper_ci = map_dbl(.x = boot_ci,
                              ~ .x$basic[[5]]))

# Delete unwanted columns
data_boot %<>%
    select(-data, -boot, -boot_ci)

# Clip CI intervals (SPARS ranges from -50 to 50)
data_boot %<>%
    mutate(upper_ci = ifelse(upper_ci > 50,
                             yes = 50,
```

```
                             no = upper_ci),
         lower_ci = ifelse(lower_ci < -50,
                           yes = -50,
                           no = lower_ci))

# Add fill column for plot
data_boot %<>%
    mutate(fill = ifelse(upper_ci >= 0 & lower_ci <= 0,
                         yes = 'inclusive',
                         no = 'exclusive'),
           fill = factor(fill,
                         levels = c('inclusive', 'exclusive'),
                         ordered = TRUE))
```

#### Plots

##### Scatter plots

```
# Plot scatter plot of ratings for each individual at every intensity
ggplot(data = data) +
    aes(x = intensity,
        y = rating,
        fill = intensity,
        colour = intensity) +
    geom_hline(yintercept = 0,
               size = 1) +
    geom_hline(yintercept = 25,
               linetype = 2) +
    geom_hline(yintercept = -25,
               linetype = 2) +
    geom_hline(yintercept = 50,
               linetype = 2) +
    geom_hline(yintercept = -50,
               linetype = 2) +
    geom_point(shape = 21,
               size = 4,
               stroke = 0.3) +
    scale_fill_gradient(low = '#CCCCCC', high = '#000000') +
    scale_colour_gradient(low = '#000000', high = '#CCCCCC') +
    scale_y_continuous(limits = c(-50, 50),
                       breaks = c(-50, 0, 50)) +
    scale_x_continuous(breaks = seq(from = 1,
                                    to = 4,
                                    by = 0.25),
                       labels = sprintf('%0.2f', round(seq(from = 1,
                                                           to = 4,
                                                           by = 0.25), 2))) +
    facet_wrap(~ PID, ncol = 4) +
    labs(title = "Individuals: Scatter plots of SPARS ratings at each stimulus intensity",
         subtitle = '- Dashed line: pain threshold\n- Colour gradient: stimulus intensity',
         x = 'Stimulus intensity (J)',
         y = 'SPARS rating (-50, 50)') +
    theme(legend.position = 'none',
          panel.grid = element_blank(),
          panel.spacing = unit(0.1, 'lines'),
```

```
strip.text = element_text(margin = margin(t = 0.1,
                                          b = 0.1,
                                          r = 1,
                                          l = 1,
                                          'lines')),
axis.text.x = element_text(angle = -90))
```

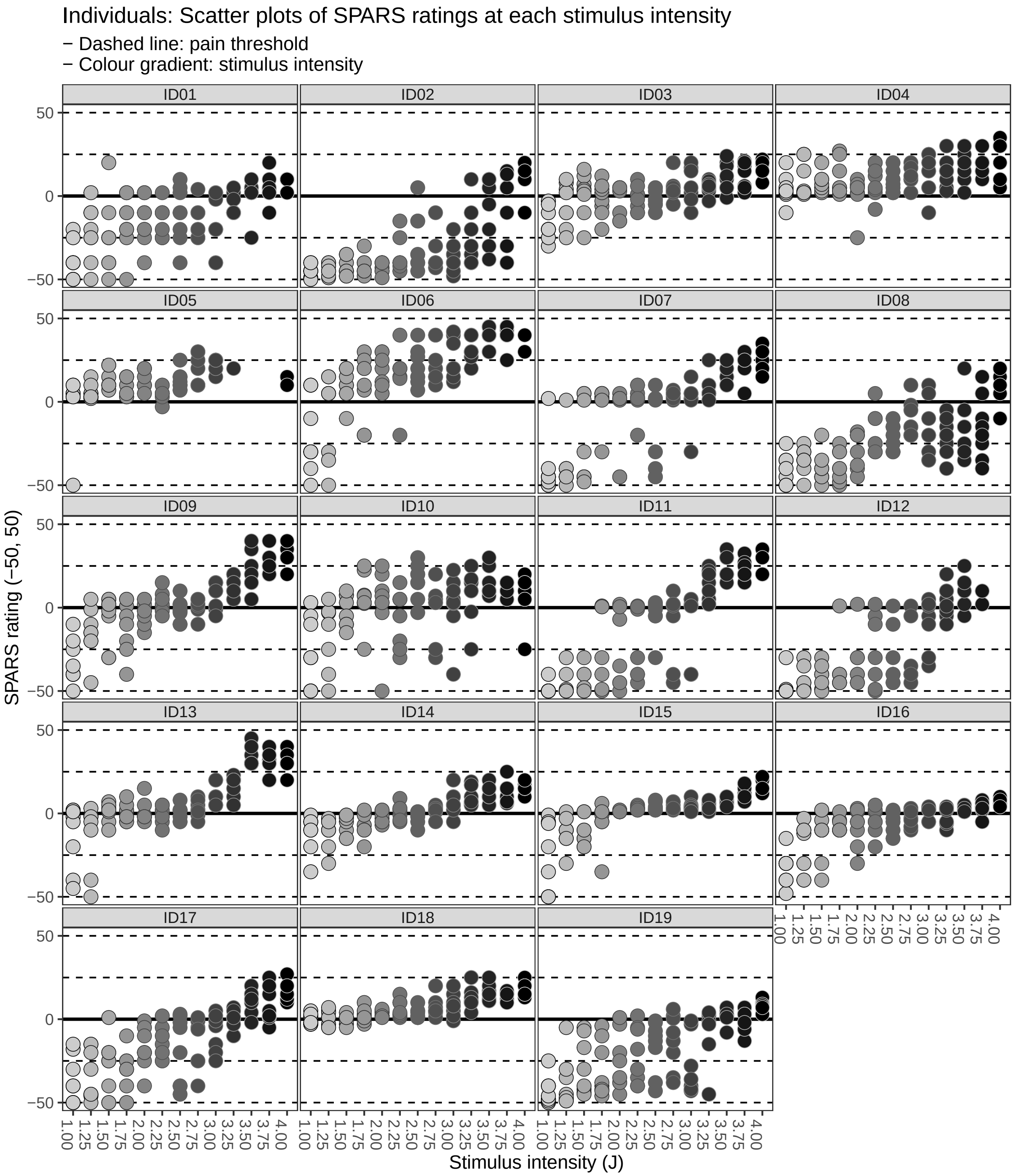

##### Trimean confidence interval plots

```
# Plot individual CIs at every intensity
ggplot(data = data_boot) +
    aes(x = intensity,
        fill = fill,
        colour = fill) +
    geom_hline(yintercept = 0,
               size = 1) +
    geom_hline(yintercept = -25,
               linetype = 2) +
    geom_hline(yintercept = 25,
               linetype = 2) +
    geom_hline(yintercept = 50,
               linetype = 2) +
    geom_hline(yintercept = -50,
               linetype = 2) +
    geom_crossbar(aes(y = tri_mean,
                      ymin = lower_ci,
                      ymax = upper_ci),
                  fatten = 0,
                  size = 0.3) +
    scale_fill_manual(values = c('#000000', '#CCCCCC')) +
    scale_colour_manual(values = c('#CCCCCC', '#000000')) +
    scale_y_continuous(limits = c(-50, 50),
                       breaks = c(-50, 0, 50)) +
    scale_x_continuous(breaks = seq(from = 1,
                                    to = 4,
                                    by = 0.25),
                       labels = sprintf('%0.2f', round(seq(from = 1,
                                                           to = 4,
                                                           by = 0.25), 2))) +
    facet_wrap(~ PID, ncol = 4) +
    labs(title = "Individuals: Crossbar plots of 95% CI of Tukey trimeans for SPARS ratings\nat each st[illegible]
         subtitle = '- Basic bootstrap 95% CI with 10,000 resamples\n- Dashed line: pain threshold | - [illegible]
         x = 'Stimulus intensity (J)',
         y = 'SPARS rating (-50, 50)') +
    theme(legend.position = 'none',
          panel.grid = element_blank(),
          panel.spacing = unit(0.1, 'lines'),
          strip.text = element_text(margin = margin(t = 0.1,
                                                    b = 0.1,
                                                    r = 1,
                                                    l = 1,
                                                    'lines')),
          axis.text.x = element_text(angle = -90))
```

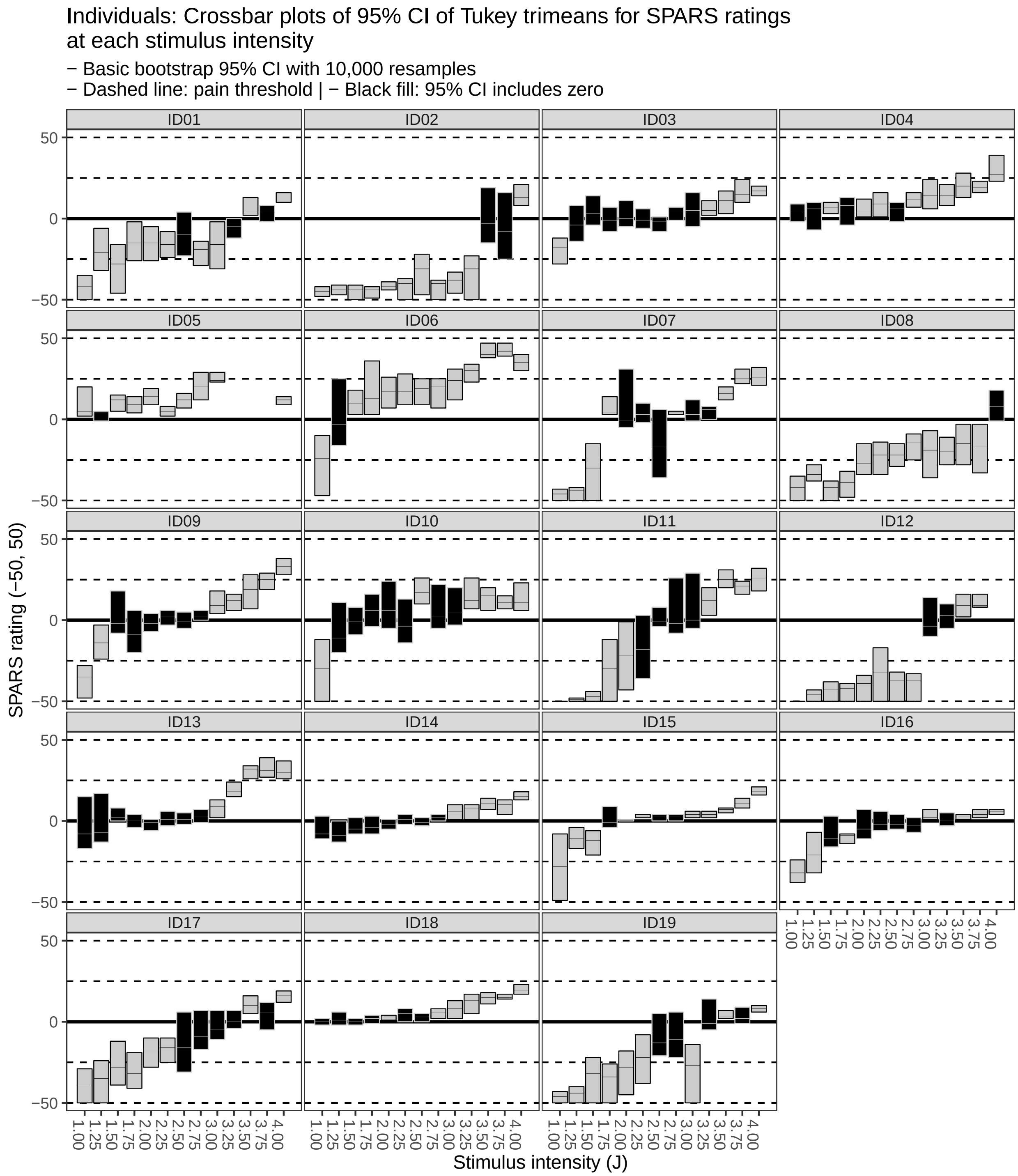

---

### Data at the level of the group

#### Bootstrapping procedure

```
# Calculate individual trimeans at each stimulus intensity
data_group <- data %>%
```

```
    group_by(PID, intensity) %>%
    summarise(tri_mean = tri_mean(rating)) %>%
    ungroup()

# Nest data in preparation for bootstrapping at each stimulus intensity
data_boot_group <- data_group %>%
    group_by(intensity) %>%
    nest()

# Perform bootstrap
set.seed(987654321)
data_boot_group %<>% mutate(boot = map(.x = data,
                                       ~ boot(data = .x$tri_mean,
                                              statistic = boot_tri_mean,
                                              R = 10000, # For small sample size
                                              stype = 'i')))

# Extract CI from boot object
data_boot_group %<>% mutate(boot_ci = map(.x = boot,
                                          ~ boot.ci(.x,
                                                    type = 'basic')))

# Extract the data, giving original median and bootstrapped CI
data_boot_group %<>% mutate(tri_mean = map(.x = boot_ci,
                                           ~ .x$t0),
                            lower_ci = map(.x = boot_ci,
                                           ~ .x$basic[[4]]),
                            upper_ci = map(.x = boot_ci,
                                           ~ .x$basic[[5]]))

# Delete unwanted columns
data_boot_group %<>% select(-data, -boot, -boot_ci) %>%
    unnest()

# Clip CI intervals (SPARS ranges from -50 to 50)
data_boot_group %<>%
    mutate(upper_ci = ifelse(upper_ci > 50,
                             yes = 50,
                             no = upper_ci),
           lower_ci = ifelse(lower_ci < -50,
                             yes = -50,
                             no = lower_ci))

# Add fill column for plot
data_boot_group %<>%
    mutate(fill = ifelse(upper_ci >= 0 & lower_ci <= 0,
                         yes = 'inclusive',
                         no = 'exclusive'),
           fill = factor(fill,
                         levels = c('inclusive', 'exclusive'),
                         ordered = TRUE))
```

#### Plots

##### Scatter plots

```
# Plot scatter plot of ratings for the group at every intensity
ggplot(data = data_group) +
    aes(x = intensity,
        y = tri_mean,
        fill = intensity,
        colour = intensity) +
    geom_hline(yintercept = 0,
               size = 1) +
    geom_hline(yintercept = -25,
               linetype = 2) +
    geom_hline(yintercept = 25,
               linetype = 2) +
    geom_hline(yintercept = 50,
               linetype = 2) +
    geom_hline(yintercept = -50,
               linetype = 2) +
    geom_point(shape = 21,
               size = 4,
               stroke = 0.3) +
    scale_fill_gradient(low = '#CCCCCC', high = '#000000') +
    scale_colour_gradient(low = '#000000', high = '#CCCCCC') +
    scale_y_continuous(limits = c(-50, 50),
                       breaks = c(-50, 0, 50)) +
    scale_x_continuous(breaks = seq(from = 1,
                                    to = 4,
                                    by = 0.25),
                       labels = sprintf('%0.2f', round(seq(from = 1,
                                                           to = 4,
                                                           by = 0.25), 2))) +
    labs(title = "Group: Scatter plots of SPARS Tukey trimean ratings\nat each stimulus intensity",
         subtitle = '- Dashed line: pain threshold\n- Colour gradient: stimulus intensity',
         x = 'Stimulus intensity (J)',
         y = 'SPARS rating (-50, 50)') +
    theme(legend.position = 'none',
          panel.grid = element_blank())
```

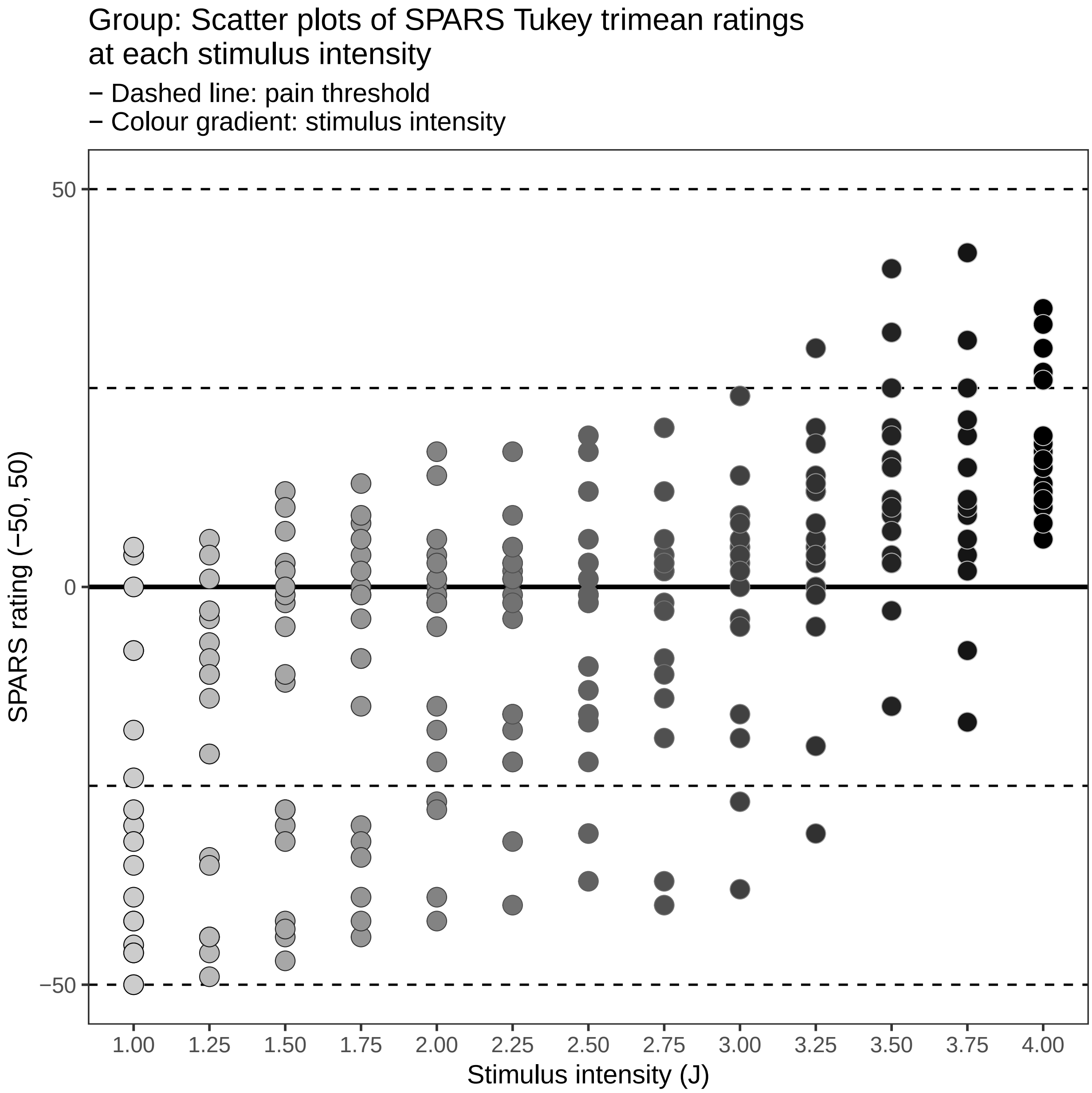

**Trimean confidence interval plots**

```
# Plot group CIs at every intensity
ggplot(data = data_boot_group) +
    aes(x = intensity) +
    geom_hline(yintercept = 0,
               size = 1) +
    geom_hline(yintercept = -25,
               linetype = 2) +
    geom_hline(yintercept = 25,
               linetype = 2) +
    geom_hline(yintercept = 50,
               linetype = 2) +
    geom_hline(yintercept = -50,
```

```
           linetype = 2) +
geom_crossbar(aes(y = tri_mean,
                  ymin = lower_ci,
                  ymax = upper_ci,
                  fill = fill,
                  colour = fill),
              fatten = 0,
              size = 0.3) +
scale_fill_manual(values = c('#000000', '#CCCCCC')) +
scale_colour_manual(values = c('#CCCCCC', '#000000')) +
scale_y_continuous(limits = c(-50, 50),
                   breaks = c(-50, 0, 50)) +
scale_x_continuous(breaks = seq(from = 1,
                                to = 4,
                                by = 0.25),
                   labels = sprintf('%0.2f', round(seq(from = 1,
                                                       to = 4,
                                                       by = 0.25), 2))) +
labs(title = "Group: Crossbar plots of 95% CI of Tukey trimeans for SPARS ratings\nat each stimulus
     subtitle = '- Basic bootstrap 95% CI with 10,000 resamples\n- Dashed line: pain threshold | - (
     x = 'Stimulus intensity (J)',
     y = 'SPARS rating (-50, 50)') +
theme(legend.position = 'none',
      panel.grid = element_blank())
```

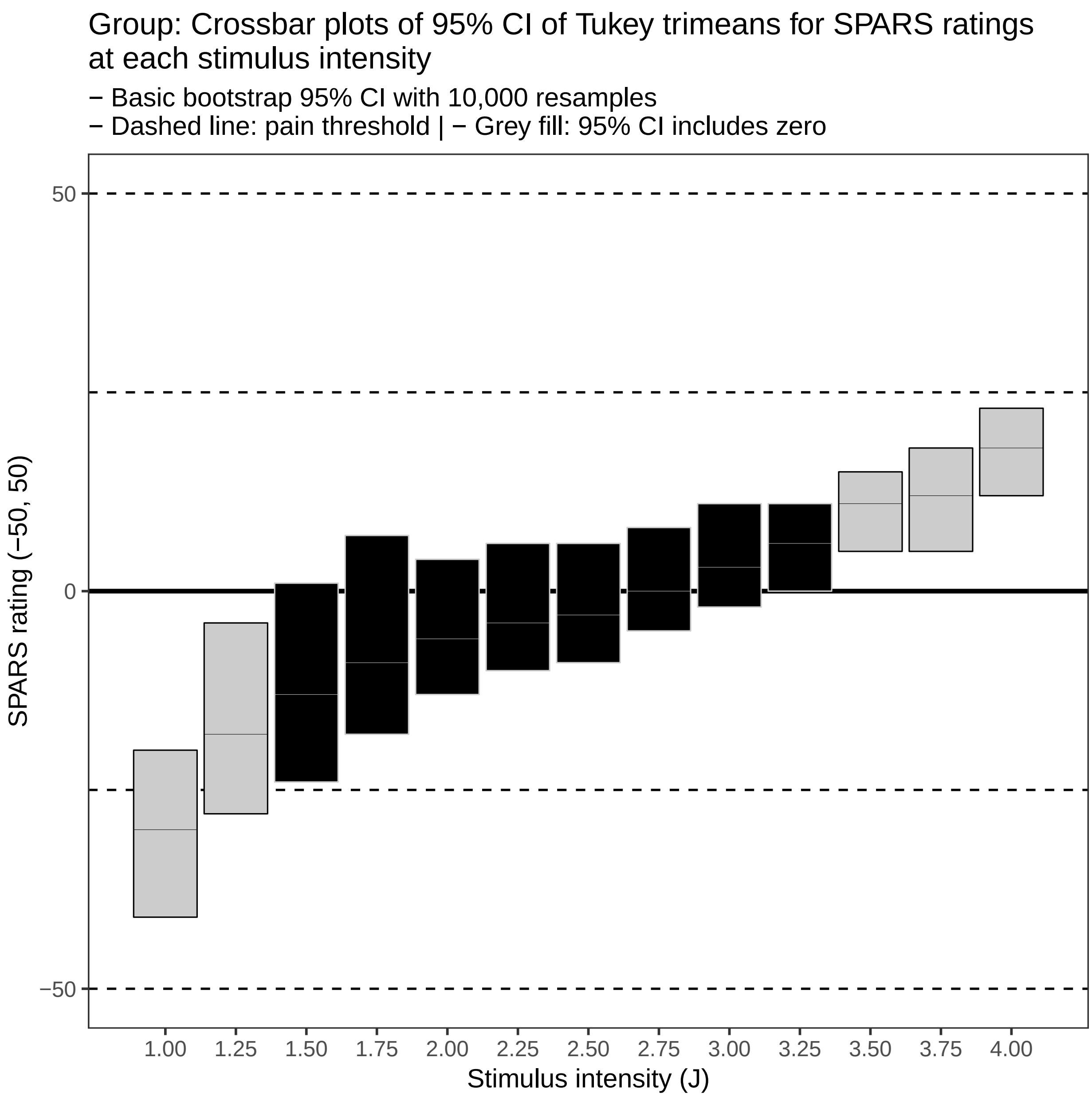

---

### Session information

```
sessionInfo()
```

```
## R version 3.5.1 (2018-07-02)
## Platform: x86_64-pc-linux-gnu (64-bit)
## Running under: Debian GNU/Linux 9 (stretch)
##
## Matrix products: default
## BLAS: /usr/lib/openblas-base/libblas.so.3
## LAPACK: /usr/lib/libopenblasp-r0.2.19.so
##
```

```
## locale:
##  [1] LC_CTYPE=en_US.UTF-8       LC_NUMERIC=C
##  [3] LC_TIME=en_US.UTF-8        LC_COLLATE=en_US.UTF-8
##  [5] LC_MONETARY=en_US.UTF-8    LC_MESSAGES=C
##  [7] LC_PAPER=en_US.UTF-8       LC_NAME=C
##  [9] LC_ADDRESS=C               LC_TELEPHONE=C
## [11] LC_MEASUREMENT=en_US.UTF-8 LC_IDENTIFICATION=C
##
## attached base packages:
## [1] stats     graphics  grDevices utils     datasets  methods   base
##
## other attached packages:
##  [1] bindrcpp_0.2.2  skimr_1.0.3     boot_1.3-20     magrittr_1.5
##  [5] forcats_0.3.0   stringr_1.3.1   dplyr_0.7.8     purrr_0.2.5
##  [9] readr_1.3.0     tidyr_0.8.2     tibble_1.4.2    ggplot2_3.1.0
## [13] tidyverse_1.2.1
##
## loaded via a namespace (and not attached):
##  [1] tidyselect_0.2.5 xfun_0.4         haven_2.0.0      lattice_0.20-35
##  [5] colorspace_1.3-2 generics_0.0.2   htmltools_0.3.6  yaml_2.2.0
##  [9] rlang_0.3.0.1    pillar_1.3.1     glue_1.3.0       withr_2.1.2
## [13] modelr_0.1.2     readxl_1.2.0     bindr_0.1.1      plyr_1.8.4
## [17] munsell_0.5.0    gtable_0.2.0     cellranger_1.1.0 rvest_0.3.2
## [21] evaluate_0.12    knitr_1.21       broom_0.5.1      Rcpp_1.0.0
## [25] scales_1.0.0     backports_1.1.3  jsonlite_1.6     hms_0.4.2
## [29] digest_0.6.18    stringi_1.2.4    grid_3.5.1       cli_1.0.1
## [33] tools_3.5.1      lazyeval_0.2.1   crayon_1.3.4     pkgconfig_2.0.2
## [37] xml2_1.2.0       lubridate_1.7.4  assertthat_0.2.0 rmarkdown_1.11
## [41] httr_1.4.0       rstudioapi_0.8   R6_2.3.0         nlme_3.1-137
## [45] compiler_3.5.1
```
