## Supplementary File 2 for "Rethinking pain threshold as a zone of uncertainty"

SPARS B (including NRS and SRS): Width of the pain threshold

*12 Jan 2019*

**Session information** **30**

---

#### Question

How wide is the pain threshold for participants taking part in the SPARS B trial?

To answer the question, we calculated the *Tukey Trimean* and bootstrapped 95% confidence interval (CI) for each individual, at each stimulus intensity. Next, we plotted these statistics to show the stimulus range over which each individual's CIs included zero on the SPARS (pain threshold).

To get an idea of the width of the stimulus range that included zero on the SPARS at the group level, we calculated the *Tukey trimean* for each individual, at each stimulus intensity, and then calculated the mean and bootstrapped 95% CI for the group at each stimulus intensity. These data were then plotted to show the stimulus range over which the group's CIs included zero (pain threshold).

The selection of the *tukey trimean* as the measure of central tendancy at the individual level was based on the analysis of central tendancy reported in the original description of the SPARS (Supplement_3.pdf). The *Tukey trimean* is defined as the weighted average of the distribution's median and its two quartiles, and is a robust measure of central tendancy that unlike a median, takes the spread of the data into account.

$$T_{mean} = \frac{1}{2}(Q_2 + \frac{Q_1 + Q_3}{2})$$

Where:

- $Q_1 = 25^{\text{th}}$ percentile
- $Q_2 = 50^{\text{th}}$ percentile (median)
- $Q_3 = 75^{\text{th}}$ percentile

```
# Define the tri_mean function
tri_mean <- function(x) {
    # Calculate quantiles
    q1 <- quantile(x, probs = 0.25, na.rm = TRUE)[[1]]
    q2 <- median(x, na.rm = TRUE)
    q3 <- quantile(x, probs = 0.75, na.rm = TRUE)[[1]]
    # Calculate trimean
    tri_mean <- (q2 + ((q1 + q3) / 2)) / 2
    # Round to a whole number
    tri_mean <- round(tri_mean)
    return(tri_mean)
}
```

**Note:** No inspection of block and stimulus order effects were undertaken because analysis of these factors in the original description of the SPARS revealed no order effects (Supplement_4.pdf).

**Note:** The three scales measure were used in the SPARS B trial (Trial B). These were:

- pain NRS: 0 (no pain) to 100 (worst pain you can imagine)
- SRS: 0 (no sensation) to 100 (pain)
- SPARS: -50 (no sensation), 0 (pain threshold), +50 (worst pain you can imagine)

The stimulus range was centred on the pre-determined pain threshold of each participant (compared to the fixed range of intensities used in Trial A), all analyses use the rank order of the nine stimulus intensities each participant was exposed to rather than the absolute intensities of the stimuli used.

The experimental design involved exposing each participant to four successive experimental blocks of 27 trials (laser stimulations) each for each of the three measurement scales. The sequence of stimulus intensities used within each block was pre-determined, and differed between blocks. The order of in which the measurement scales were assessed was randomized, but for convenience of reporting, the plots are always shown in the order: pain NRS, SRS, and SPARS.

---

### Import and inspect data

```
# Import
data <- read_rds('data-cleaned/SPARS_B.rds')
```

```
# Rank stimulus intensity
```

```
data %<>%
    group_by(PID, scale) %>%
    arrange(intensity) %>%
    mutate(intensity_rank = dense_rank(intensity)) %>%
    select(-intensity) %>%
    rename(intensity = intensity_rank) %>%
    ungroup()

# Inspect
glimpse(data)
```

```
## Observations: 6,771
## Variables: 6
## $ PID          <chr> "ID06", "ID06", "ID06", "ID06", "ID06", "ID06", "...
## $ block_number <int> 1, 1, 1, 1, 1, 1, 1, 1, 1, 2, 2, 2, 2, 2, 2, 2, 2...
## $ trial_number <dbl> 4, 4, 4, 6, 6, 6, 27, 27, 27, 9, 9, 9, 13, 13, 13...
## $ scale        <chr> "SPARS", "NRS", "SRS", "SPARS", "NRS", "SRS", "SP...
## $ rating       <dbl> -49, NA, NA, 2, NA, NA, -6, NA, NA, 3, NA, NA, -2...
## $ intensity    <int> 1, 1, 1, 1, 1, 1, 1, 1, 1, 1, 1, 1, 1, 1, 1, 1, 1...
```

```
data %>%
    select(intensity, rating) %>%
    skim()
```

```
## Skim summary statistics
##  n obs: 6771
##  n variables: 2
##
## -- Variable type:integer --------------------------------------------------------------
##   variable missing complete    n mean   sd p0 p25 p50 p75 p100     hist
##  intensity       0     6771 6771    5 2.58  1   3   5   7    9
##
## -- Variable type:numeric --------------------------------------------------------------
##  variable missing complete    n   mean    sd   p0 p25 p50 p75 p100
##    rating    4622     2149 6771 -12.53 41.35 -100 -38   0   6   98
##      hist
##
```

---

### Data at the level of the individual

#### Bootstrapping procedure for SPARS, NRS, and SRS

```
############################################################
#                                                          #
#                          SPARS                           #
#                                                          #
############################################################
# Extract SPARS data
data_spars <- data %>%
    filter(scale == 'SPARS') %>%
    filter(!is.na(rating))

# Nest data in preparation for bootstrapping at each stimulus intensity
spars_boot <- data_spars %>%
```

```r
    group_by(PID, intensity) %>%
    nest()

# Define bootstrap function
boot_tri_mean <- function(d,i){
    tri_mean(d[i])
}

# Perform bootstrap
set.seed(123456789)
spars_boot %<>%
        mutate(boot = map(.x = data,
                          ~ boot(data = .x$rating,
                                 statistic = boot_tri_mean,
                                 R = 10000, # For small sample size
                                 stype = 'i')))

# Extract CI from boot object
spars_boot %<>%
    mutate(boot_ci = map(.x = boot,
                         ~ boot.ci(.x,
                                   type = 'basic')))

# Extract the data, giving original trimean and bootstrapped CI
spars_boot %<>%
    mutate(tri_mean = map_dbl(.x = boot_ci,
                              ~ .x$t0),
           lower_ci = map_dbl(.x = boot_ci,
                              ~ .x$basic[[4]]),
           upper_ci = map_dbl(.x = boot_ci,
                              ~ .x$basic[[5]]))

# Delete unwanted columns
spars_boot %<>%
    select(-data, -boot, -boot_ci)

# Clip CI intervals (SPARS ranges from -50 to 50)
spars_boot %<>%
    mutate(upper_ci = ifelse(upper_ci > 50,
                             yes = 50,
                             no = upper_ci),
           lower_ci = ifelse(lower_ci < -50,
                             yes = -50,
                             no = lower_ci))

# Add fill column for plot
spars_boot %<>%
    mutate(fill = ifelse(upper_ci >= 0 & lower_ci <= 0,
                         yes = 'inclusive',
                         no = 'exclusive'),
           fill = factor(fill,
                         levels = c('inclusive', 'exclusive'),
                         ordered = TRUE))
```

```r
############################################################
#                                                          #
#                          NRS                             #
#                                                          #
############################################################
# Extract NRS data
data_nrs <- data %>%
    filter(scale == 'NRS') %>%
    filter(!is.na(rating))

# Nest data in preparation for bootstrapping at each stimulus intensity
nrs_boot <- data_nrs %>%
    group_by(PID, intensity) %>%
    nest()

# Define bootstrap function
boot_tri_mean <- function(d,i){
    tri_mean(d[i])
}

# Perform bootstrap
set.seed(123456789)
nrs_boot %<>%
        mutate(boot = map(.x = data,
                          ~ boot(data = .x$rating,
                                 statistic = boot_tri_mean,
                                 R = 10000, # For small sample size
                                 stype = 'i')))

# Extract CI from boot object
nrs_boot %<>%
    mutate(boot_ci = map(.x = boot,
                         ~ boot.ci(.x,
                                   type = 'basic')))

# Extract the data, giving original trimean and bootstrapped CI
nrs_boot %<>%
    mutate(tri_mean = map_dbl(.x = boot_ci,
                              ~ .x$t0),
           lower_ci = map_dbl(.x = boot_ci,
                              ~ .x$basic[[4]]),
           upper_ci = map_dbl(.x = boot_ci,
                              ~ .x$basic[[5]]))

# Delete unwanted columns
nrs_boot %<>%
    select(-data, -boot, -boot_ci)

# Clip CI intervals (NRS ranges from 0 to 100)
nrs_boot %<>%
    mutate(upper_ci = ifelse(upper_ci > 100,
                             yes = 100,
                             no = upper_ci),
           lower_ci = ifelse(lower_ci < 0,
```

```
                              yes = 0,
                              no = lower_ci))

# Add fill column for plot
nrs_boot %<>%
    mutate(fill = ifelse(lower_ci == 0,
                         yes = 'inclusive',
                         no = 'exclusive'),
           fill = factor(fill,
                         levels = c('inclusive', 'exclusive'),
                         ordered = TRUE))

############################################################
#                                                          #
#                           SRS                            #
#                                                          #
############################################################
# Extract SRS data
data_srs <- data %>%
    filter(scale == 'SRS') %>%
    filter(!is.na(rating)) %>%
    # Remove ID01 (didn't complete SRS)
    filter(PID != 'ID01')

# Nest data in preparation for bootstrapping at each stimulus intensity
srs_boot <- data_srs %>%
    group_by(PID, intensity) %>%
    nest()

# Define bootstrap function
boot_tri_mean <- function(d,i){
    tri_mean(d[i])
}

# Perform bootstrap
set.seed(123456789)
srs_boot %<>%
        mutate(boot = map(.x = data,
                          ~ boot(data = .x$rating,
                                 statistic = boot_tri_mean,
                                 R = 10000, # For small sample size
                                 stype = 'i')))

# Extract CI from boot object
srs_boot %<>%
    mutate(boot_ci = map(.x = boot,
                         ~ boot.ci(.x,
                                   type = 'basic')))

# Extract the data, giving original trimean and bootstrapped CI
srs_boot %<>%
    mutate(tri_mean = map_dbl(.x = boot_ci,
                              ~ .x$t0),
           lower_ci = map_dbl(.x = boot_ci,
```

```r
                               ~ .x$basic[[4]]),
             upper_ci = map_dbl(.x = boot_ci,
                               ~ .x$basic[[5]]))

# Delete unwanted columns
srs_boot %<>%
    select(-data, -boot, -boot_ci)

# Clip CI intervals (SRS ranges from -100 to 0)
srs_boot %<>%
    mutate(upper_ci = ifelse(upper_ci > 0,
                             yes = 0,
                             no = upper_ci),
           lower_ci = ifelse(lower_ci < -100,
                             yes = -100,
                             no = lower_ci))

# Add fill column for plot
srs_boot %<>%
    mutate(fill = ifelse(upper_ci == 0,
                         yes = 'inclusive',
                         no = 'exclusive'),
           fill = factor(fill,
                         levels = c('inclusive', 'exclusive'),
                         ordered = TRUE))
```

#### Plots

##### Scatter plots

###### SPARS

```r
# Plot scatter plot of ratings for each individual at every intensity
ggplot(data = data_spars) +
    aes(x = intensity,
        y = rating,
        fill = intensity,
        colour = intensity) +
    geom_hline(yintercept = 0,
               size = 1) +
    geom_hline(yintercept = 25,
               linetype = 2) +
    geom_hline(yintercept = -25,
               linetype = 2) +
    geom_hline(yintercept = 50,
               linetype = 2) +
    geom_hline(yintercept = -50,
               linetype = 2) +
    geom_point(shape = 21,
               size = 4,
               stroke = 0.3) +
    scale_fill_gradient(low = '#CCCCCC', high = '#000000') +
    scale_colour_gradient(low = '#000000', high = '#CCCCCC') +
    scale_y_continuous(limits = c(-50, 50),
                       breaks = c(-50, 0, 50)) +
```

```r
    scale_x_continuous(breaks = seq(from = 1,
                                    to = 9,
                                    by = 1)) +
    facet_wrap(~ PID, ncol = 4) +
    labs(title = "SPARS individuals: Scatter plots of ratings at each stimulus intensity",
         subtitle = '- Dashed line: pain threshold\n- Colour gradient: stimulus intensity',
         x = 'Rank stimulus intensity (0.25J increments)',
         y = 'SPARS rating (-50, 50)') +
    theme(legend.position = 'none',
          panel.grid = element_blank(),
          panel.spacing = unit(0.1, 'lines'),
          strip.text = element_text(margin = margin(t = 0.1,
                                                    b = 0.1,
                                                    r = 1,
                                                    l = 1,
                                                    'lines')))
```

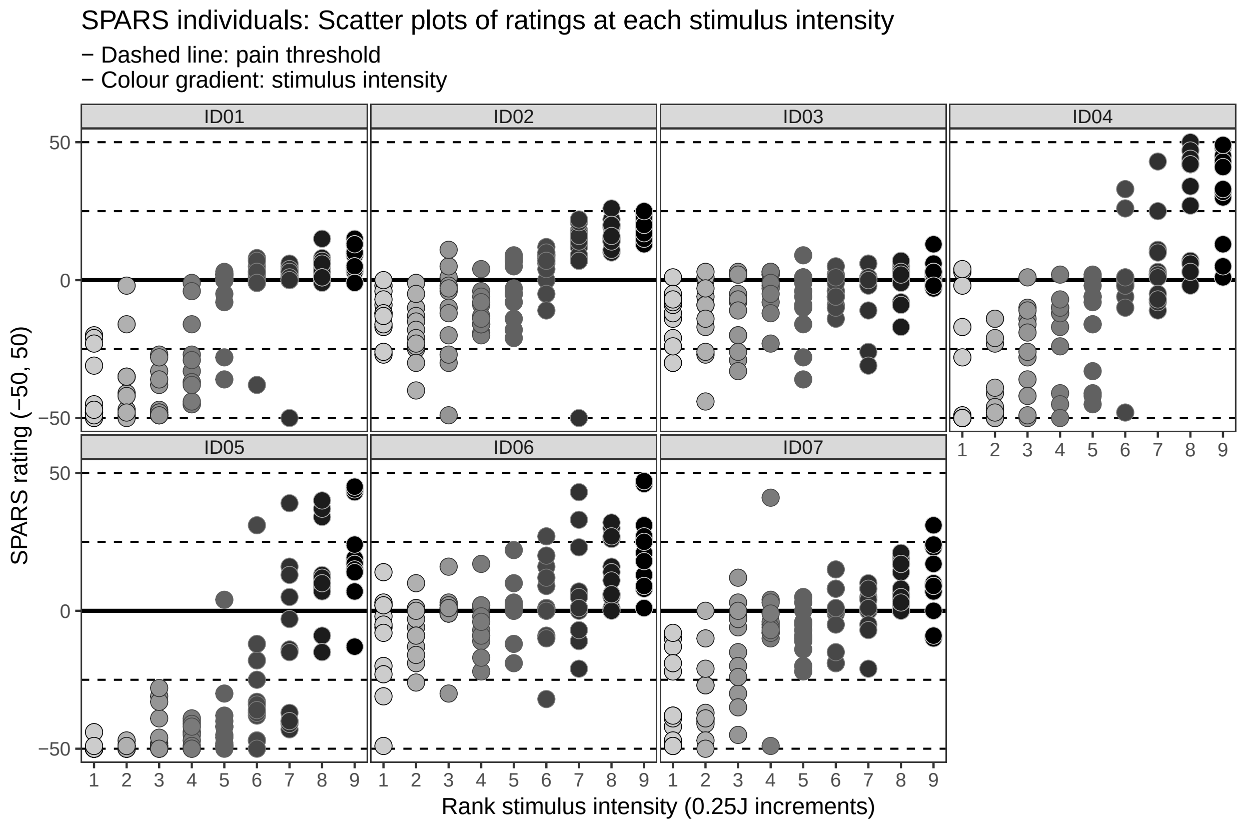

#### NRS

```r
# Plot scatter plot of ratings for each individual at every intensity
ggplot(data = data_nrs) +
    aes(x = intensity,
        y = rating,
        fill = intensity,
        colour = intensity) +
    geom_hline(yintercept = 0,
               size = 1) +
```

```r
geom_hline(yintercept = 25,
           linetype = 2) +
geom_hline(yintercept = 75,
           linetype = 2) +
geom_hline(yintercept = 100,
           linetype = 2) +
geom_point(shape = 21,
           size = 4,
           stroke = 0.3) +
scale_fill_gradient(low = '#CCCCCC', high = '#000000') +
scale_colour_gradient(low = '#000000', high = '#CCCCCC') +
scale_y_continuous(limits = c(0, 100),
                   breaks = c(0, 50, 100)) +
scale_x_continuous(breaks = seq(from = 1,
                                to = 9,
                                by = 1)) +
facet_wrap(~ PID, ncol = 4) +
labs(title = "NRS individuals: Scatter plots of ratings at each stimulus intensity",
     subtitle = '- Dashed line: pain threshold\n- Colour gradient: stimulus intensity',
     x = 'Rank stimulus intensity (0.25J increments)',
     y = 'NRS rating (0, 100)') +
theme(legend.position = 'none',
      panel.grid = element_blank(),
      panel.spacing = unit(0.1, 'lines'),
      strip.text = element_text(margin = margin(t = 0.1,
                                                b = 0.1,
                                                r = 1,
                                                l = 1,
                                                'lines')))
```

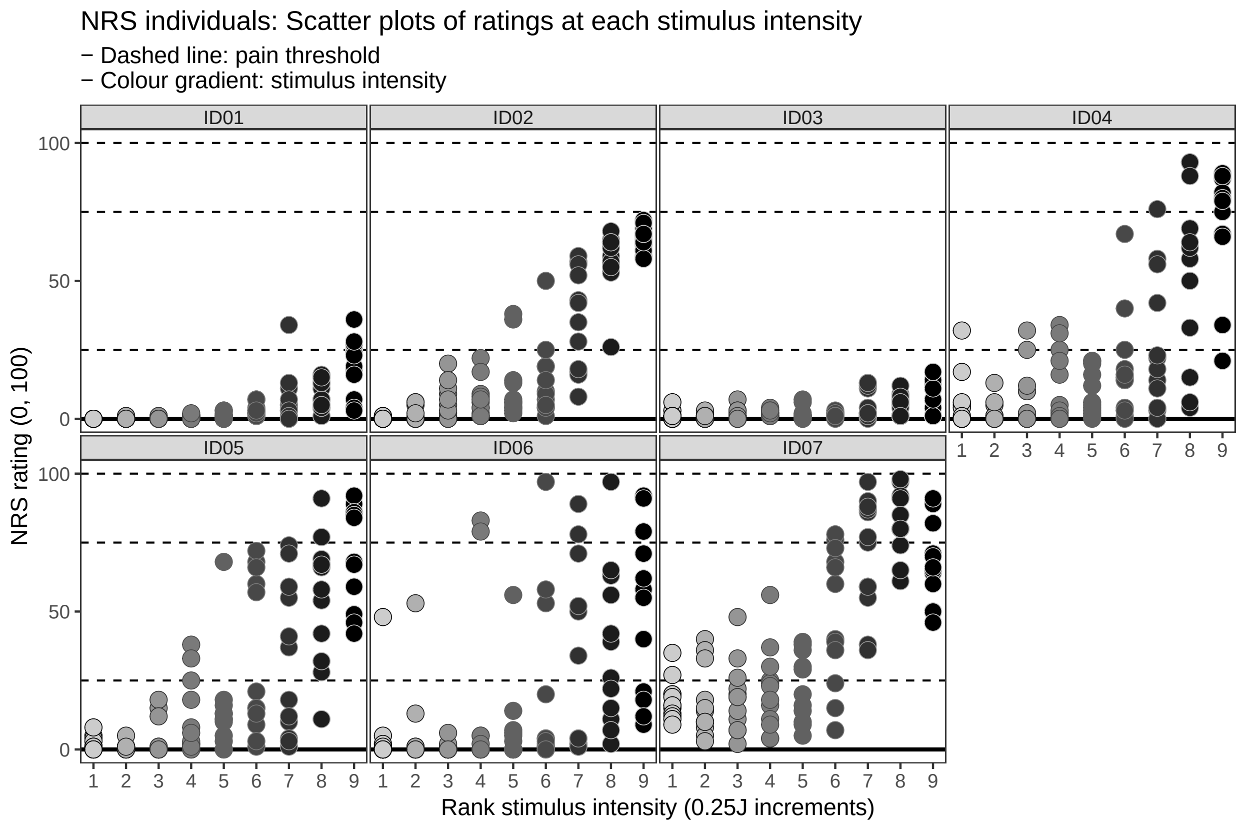

##### SRS

```
# Plot scatter plot of ratings for each individual at every intensity
ggplot(data = data_srs) +
    aes(x = intensity,
        y = rating,
        fill = intensity,
        colour = intensity) +
    geom_hline(yintercept = 0,
               size = 1) +
    geom_hline(yintercept = -25,
               linetype = 2) +
    geom_hline(yintercept = -75,
               linetype = 2) +
    geom_hline(yintercept = -100,
               linetype = 2) +
    geom_point(shape = 21,
               size = 4,
               stroke = 0.3) +
    scale_fill_gradient(low = '#CCCCCC', high = '#000000') +
    scale_colour_gradient(low = '#000000', high = '#CCCCCC') +
    scale_y_continuous(limits = c(-100, 0),
                       breaks = c(-100, -50, 0)) +
    scale_x_continuous(breaks = seq(from = 1,
                                    to = 9,
                                    by = 1)) +
    facet_wrap(~ PID, ncol = 4) +
```

```
    labs(title = "SRS individuals: Scatter plots of ratings at each stimulus intensity",
         subtitle = '- Dashed line: pain threshold\n- Colour gradient: stimulus intensity',
         x = 'Rank stimulus intensity (0.25J increments)',
         y = 'SRS rating (-100, 0)') +
    theme(legend.position = 'none',
          panel.grid = element_blank(),
          panel.spacing = unit(0.1, 'lines'),
          strip.text = element_text(margin = margin(t = 0.1,
                                                    b = 0.1,
                                                    r = 1,
                                                    l = 1,
                                                    'lines')))
```

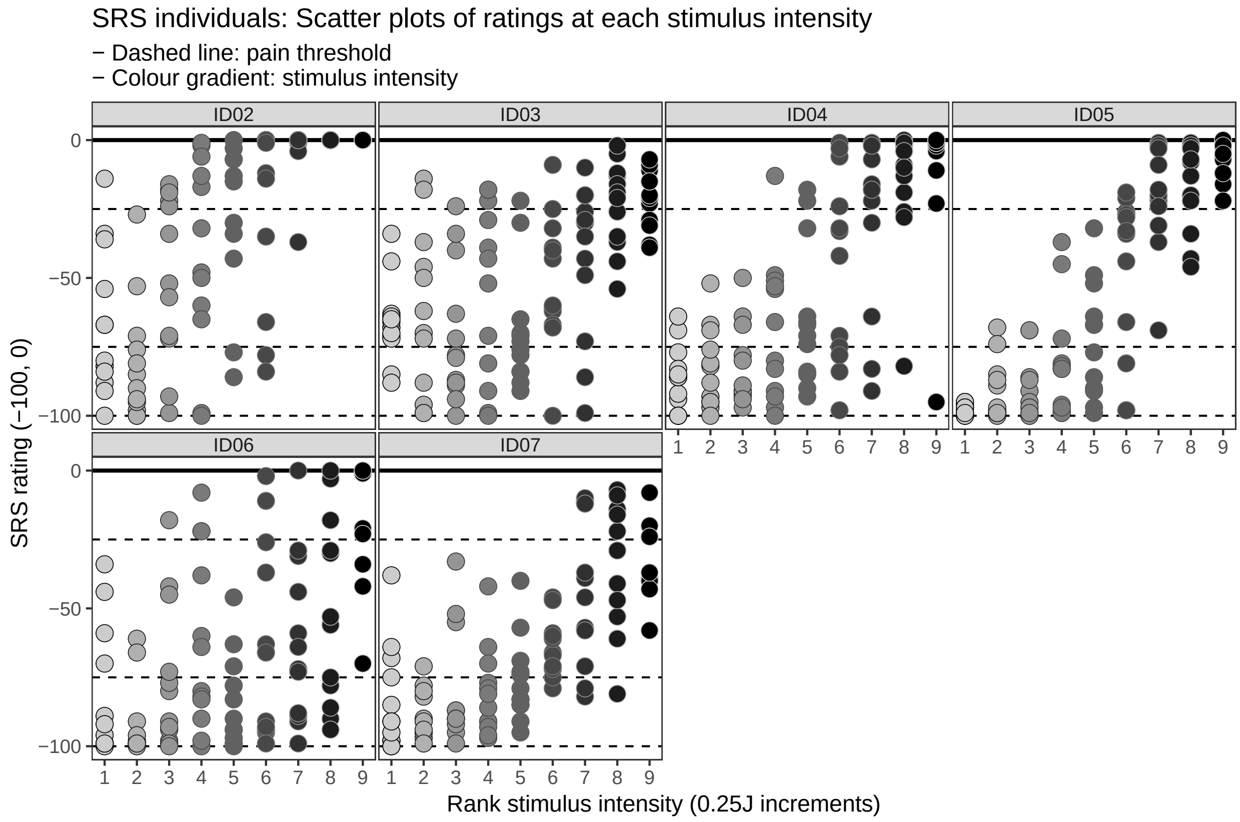

##### Trimean confidence interval plots

##### SPARS

```
# Plot individual CIs at every intensity
ggplot(data = spars_boot) +
    aes(x = intensity,
        fill = fill,
        colour = fill) +
    geom_hline(yintercept = 0,
               size = 1) +
    geom_hline(yintercept = 25,
               linetype = 2) +
    geom_hline(yintercept = -25,
```

```
               linetype = 2) +
geom_hline(yintercept = 50,
           linetype = 2) +
geom_hline(yintercept = -50,
           linetype = 2) +
geom_crossbar(aes(y = tri_mean,
                  ymin = lower_ci,
                  ymax = upper_ci),
              fatten = 0,
              size = 0.3) +
scale_fill_manual(values = c('#000000', '#CCCCCC')) +
scale_colour_manual(values = c('#CCCCCC', '#000000')) +
scale_y_continuous(limits = c(-50, 50),
                   breaks = c(-50, 0, 50)) +
scale_x_continuous(breaks = seq(from = 1,
                                to = 9,
                                by = 1)) +
facet_wrap(~ PID, ncol = 4) +
labs(title = "SPARS individuals: Crossbar plots of 95% CI of Tukey trimeans\nfor SPARS ratings at ea
     subtitle = '- Basic bootstrap 95% CI with 10,000 resamples\n- Dashed line: pain threshold | - I
     x = 'Rank stimulus intensity (0.25J increments)',
     y = 'SPARS rating (-50, 50)') +
theme(legend.position = 'none',
      panel.grid = element_blank(),
      panel.spacing = unit(0.1, 'lines'),
      strip.text = element_text(margin = margin(t = 0.1,
                                                b = 0.1,
                                                r = 1,
                                                l = 1,
                                                'lines')))
```

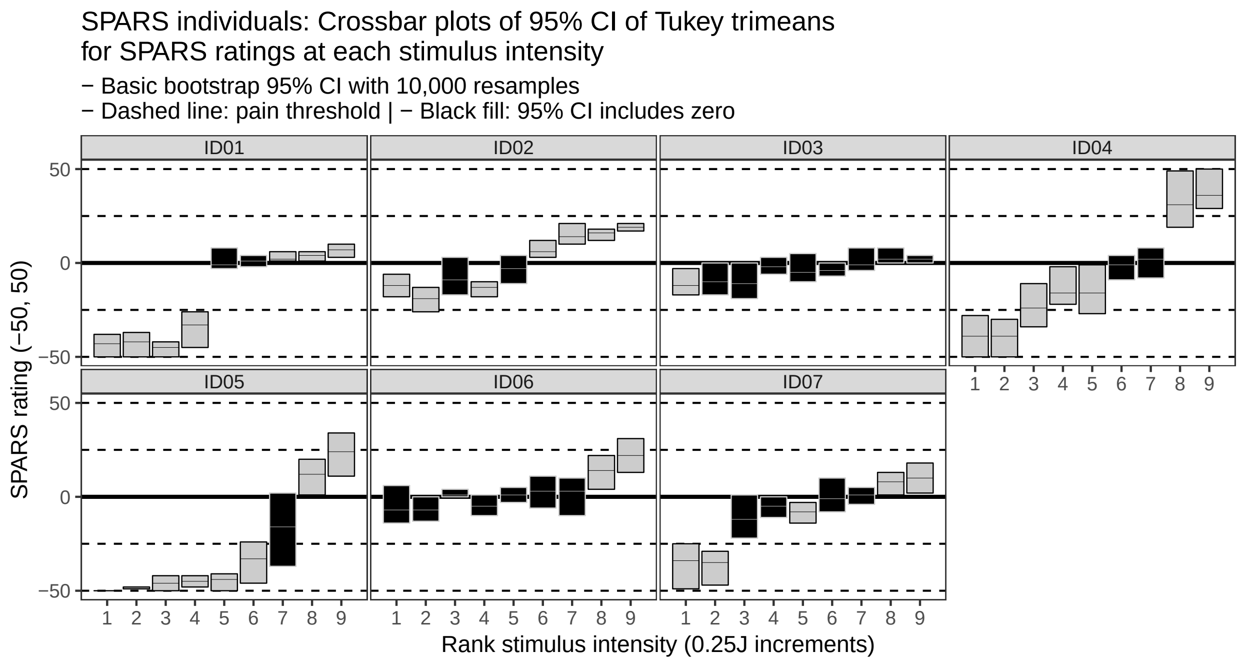

**NRS**

```
# Plot individual CIs at every intensity
ggplot(data = nrs_boot) +
    aes(x = intensity,
        fill = fill,
        colour = fill) +
    geom_hline(yintercept = 0,
               size = 1) +
    geom_hline(yintercept = 25,
               linetype = 2) +
    geom_hline(yintercept = 75,
               linetype = 2) +
    geom_hline(yintercept = 100,
               linetype = 2) +
    geom_crossbar(aes(y = tri_mean,
                      ymin = lower_ci,
                      ymax = upper_ci),
                  fatten = 0,
                  size = 0.3) +
    scale_fill_manual(values = c('#000000', '#CCCCCC')) +
    scale_colour_manual(values = c('#CCCCCC', '#000000')) +
    scale_y_continuous(limits = c(0, 100),
                       breaks = c(0, 50, 100)) +
    scale_x_continuous(breaks = seq(from = 1,
                                    to = 9,
                                    by = 1)) +
    facet_wrap(~ PID, ncol = 3) +
    labs(title = "NRS individuals: Crossbar plots of 95% CI of Tukey trimeans\nfor NRS ratings at each s
         subtitle = '- Basic bootstrap 95% CI with 10,000 resamples\n- Dashed line: pain threshold | - I
         x = 'Rank stimulus intensity (0.25J increments)',
         y = 'NRS rating (0, 100)') +
    theme(legend.position = 'none',
          panel.grid = element_blank(),
          panel.spacing = unit(0.1, 'lines'),
          strip.text = element_text(margin = margin(t = 0.1,
                                                    b = 0.1,
                                                    r = 1,
                                                    l = 1,
                                                    'lines')))
```

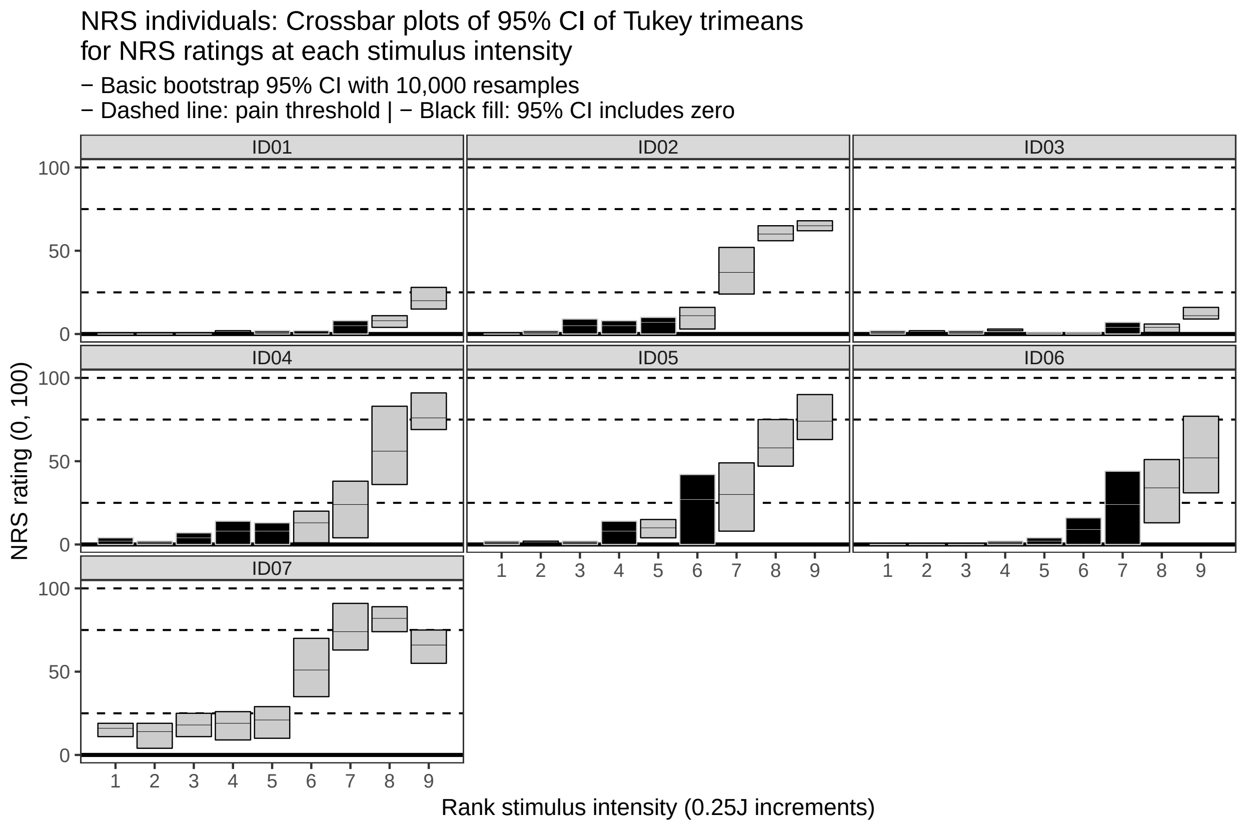

##### SRS

```
# Plot individual CIs at every intensity
ggplot(data = srs_boot) +
    aes(x = intensity,
        fill = fill,
        colour = fill) +
    geom_hline(yintercept = 0,
               size = 1) +
    geom_hline(yintercept = -25,
               linetype = 2) +
    geom_hline(yintercept = -50,
               linetype = 2) +
    geom_hline(yintercept = -75,
               linetype = 2) +
    geom_hline(yintercept = -100,
               linetype = 2) +
    geom_crossbar(aes(y = tri_mean,
                      ymin = lower_ci,
                      ymax = upper_ci),
                  fatten = 0,
                  size = 0.3) +
    scale_fill_manual(values = c('#000000', '#CCCCCC')) +
    scale_colour_manual(values = c('#CCCCCC', '#000000')) +
    scale_y_continuous(limits = c(-100, 0),
                       breaks = c(-100, -50, 0)) +
    scale_x_continuous(breaks = seq(from = 1,
```

```
                                  to = 9,
                                  by = 1)) +
  facet_wrap(~ PID, ncol = 3) +
  labs(title = "SRS individuals: Crossbar plots of 95% CI of Tukey trimeans\nfor SRS ratings at each s
       subtitle = '- Basic bootstrap 95% CI with 10,000 resamples\n- Dashed line: pain threshold | - (
       x = 'Rank stimulus intensity (0.25J increments)',
       y = 'SRS rating (-100, 0)') +
  theme(legend.position = 'none',
        panel.grid = element_blank(),
        panel.spacing = unit(0.1, 'lines'),
        strip.text = element_text(margin = margin(t = 0.1,
                                                  b = 0.1,
                                                  r = 1,
                                                  l = 1,
                                                  'lines')))
```

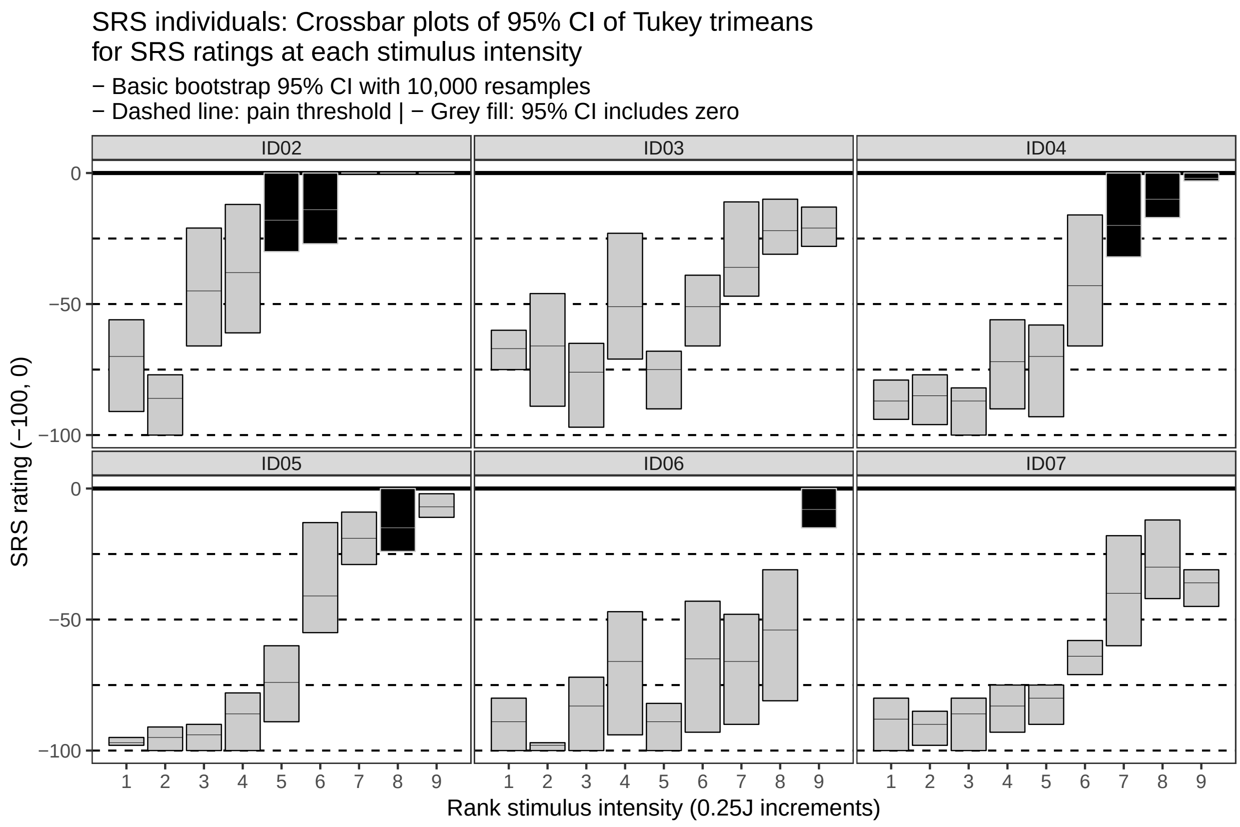

---

### Data at the level of the group

#### Bootstrapping procedure

```
############################################################
#                                                          #
#                          SPARS                           #
#                                                          #
```

```r
############################################################
# Calculate individual trimeans at each stimulus intensity
group_spars <- data_spars %>%
    group_by(PID, intensity) %>%
    summarise(tri_mean = tri_mean(rating)) %>%
    ungroup()

# Nest data in preparation for bootstrapping at each stimulus intensity
spars_boot_group <- group_spars %>%
    group_by(intensity) %>%
    nest()

# Perform bootstrap
set.seed(987654321)
spars_boot_group %<>% mutate(boot = map(.x = data,
                                        ~ boot(data = .x$tri_mean,
                                               statistic = boot_tri_mean,
                                               R = 10000, # For small sample size
                                               stype = 'i')))

# Extract CI from boot object
spars_boot_group %<>% mutate(boot_ci = map(.x = boot,
                                           ~ boot.ci(.x,
                                                     type = 'basic')))

# Extract the data, giving original median and bootstrapped CI
spars_boot_group %<>% mutate(tri_mean = map(.x = boot_ci,
                                            ~ .x$t0),
                             lower_ci = map(.x = boot_ci,
                                            ~ .x$basic[[4]]),
                             upper_ci = map(.x = boot_ci,
                                            ~ .x$basic[[5]]))

# Delete unwanted columns
spars_boot_group %<>% select(-data, -boot, -boot_ci) %>%
    unnest()

# Clip CI intervals (SPARS ranges from -50 to 50)
spars_boot_group %<>%
    mutate(upper_ci = ifelse(upper_ci > 50,
                             yes = 50,
                             no = upper_ci),
           lower_ci = ifelse(lower_ci < -50,
                             yes = -50,
                             no = lower_ci))

# Add fill column for plot
spars_boot_group %<>%
    mutate(fill = ifelse(upper_ci >= 0 & lower_ci <= 0,
                         yes = 'inclusive',
                         no = 'exclusive'),
           fill = factor(fill,
                         levels = c('inclusive', 'exclusive'),
                         ordered = TRUE))
```

```
############################################################
#                                                          #
#                           NRS                            #
#                                                          #
############################################################
# Calculate individual trimeans at each stimulus intensity
group_nrs <- data_nrs %>%
    group_by(PID, intensity) %>%
    summarise(tri_mean = tri_mean(rating)) %>%
    ungroup()

# Nest data in preparation for bootstrapping at each stimulus intensity
nrs_boot_group <- group_nrs %>%
    group_by(intensity) %>%
    nest()

# Perform bootstrap
set.seed(987654321)
nrs_boot_group %<>% mutate(boot = map(.x = data,
                                      ~ boot(data = .x$tri_mean,
                                             statistic = boot_tri_mean,
                                             R = 10000, # For small sample size
                                             stype = 'i')))

# Extract CI from boot object
nrs_boot_group %<>% mutate(boot_ci = map(.x = boot,
                                         ~ boot.ci(.x,
                                                   type = 'basic')))

# Extract the data, giving original median and bootstrapped CI
nrs_boot_group %<>% mutate(tri_mean = map(.x = boot_ci,
                                          ~ .x$t0),
                           lower_ci = map(.x = boot_ci,
                                          ~ .x$basic[[4]]),
                           upper_ci = map(.x = boot_ci,
                                          ~ .x$basic[[5]]))

# Delete unwanted columns
nrs_boot_group %<>% select(-data, -boot, -boot_ci) %>%
    unnest()

# Clip CI intervals (NRS ranges from 0 to 100)
nrs_boot_group %<>%
    mutate(upper_ci = ifelse(upper_ci > 100,
                             yes = 100,
                             no = upper_ci),
           lower_ci = ifelse(lower_ci < 0,
                             yes = 0,
                             no = lower_ci))

# Add fill column for plot
nrs_boot_group %<>%
    mutate(fill = ifelse(lower_ci == 0,
```

```r
                            yes = 'inclusive',
                            no = 'exclusive'),
              fill = factor(fill,
                            levels = c('inclusive', 'exclusive'),
                            ordered = TRUE))

############################################################
#                                                          #
#                           SRS                            #
#                                                          #
############################################################
# Calculate individual trimeans at each stimulus intensity
group_srs <- data_srs %>%
    group_by(PID, intensity) %>%
    summarise(tri_mean = tri_mean(rating)) %>%
    ungroup()

# Nest data in preparation for bootstrapping at each stimulus intensity
srs_boot_group <- group_srs %>%
    group_by(intensity) %>%
    nest()

# Perform bootstrap
set.seed(987654321)
srs_boot_group %<>% mutate(boot = map(.x = data,
                                      ~ boot(data = .x$tri_mean,
                                             statistic = boot_tri_mean,
                                             R = 10000, # For small sample size
                                             stype = 'i')))

# Extract CI from boot object
srs_boot_group %<>% mutate(boot_ci = map(.x = boot,
                                         ~ boot.ci(.x,
                                                   type = 'basic')))

# Extract the data, giving original median and bootstrapped CI
srs_boot_group %<>% mutate(tri_mean = map(.x = boot_ci,
                                          ~ .x$t0),
                           lower_ci = map(.x = boot_ci,
                                          ~ .x$basic[[4]]),
                           upper_ci = map(.x = boot_ci,
                                          ~ .x$basic[[5]]))

# Delete unwanted columns
srs_boot_group %<>% select(-data, -boot, -boot_ci) %>%
    unnest()

# Clip CI intervals (SRS ranges from -100 to 0)
srs_boot_group %<>%
    mutate(upper_ci = ifelse(upper_ci > 0,
                             yes = 0,
                             no = upper_ci),
           lower_ci = ifelse(lower_ci < -100,
                             yes = -100,
```

```r
                              no = lower_ci))

# Add fill column for plot
srs_boot_group %<>%
    mutate(fill = ifelse(upper_ci == 0,
                         yes = 'inclusive',
                         no = 'exclusive'),
           fill = factor(fill,
                         levels = c('inclusive', 'exclusive'),
                         ordered = TRUE))
```

#### Plots

##### Scatter plot

##### SPARS

```r
# Plot scatter plot of ratings for the group at every intensity
ggplot(data = group_spars) +
    aes(x = intensity,
        y = tri_mean,
        fill = intensity,
        colour = intensity) +
    geom_hline(yintercept = 0,
               size = 1) +
    geom_hline(yintercept = 25,
               linetype = 2) +
    geom_hline(yintercept = -25,
               linetype = 2) +
    geom_hline(yintercept = 50,
               linetype = 2) +
    geom_hline(yintercept = -50,
               linetype = 2) +
    geom_point(shape = 21,
               size = 4,
               stroke = 0.3) +
    scale_fill_gradient(low = '#CCCCCC', high = '#000000') +
    scale_colour_gradient(low = '#000000', high = '#CCCCCC') +
    scale_y_continuous(limits = c(-50, 50),
                       breaks = c(-50, 0, 50)) +
    scale_x_continuous(breaks = seq(from = 1,
                                    to = 9,
                                    by = 1)) +
    labs(title = "SPARS group: Scatter plots of Tukey trimean ratings at each stimulus intensity",
         subtitle = '- Dashed line: pain threshold\n- Colour gradient: stimulus intensity',
         x = 'Rank stimulus intensity (0.25J increments)',
         y = 'SPARS rating (-50, 50)') +
    theme(legend.position = 'none',
          panel.grid = element_blank())
```

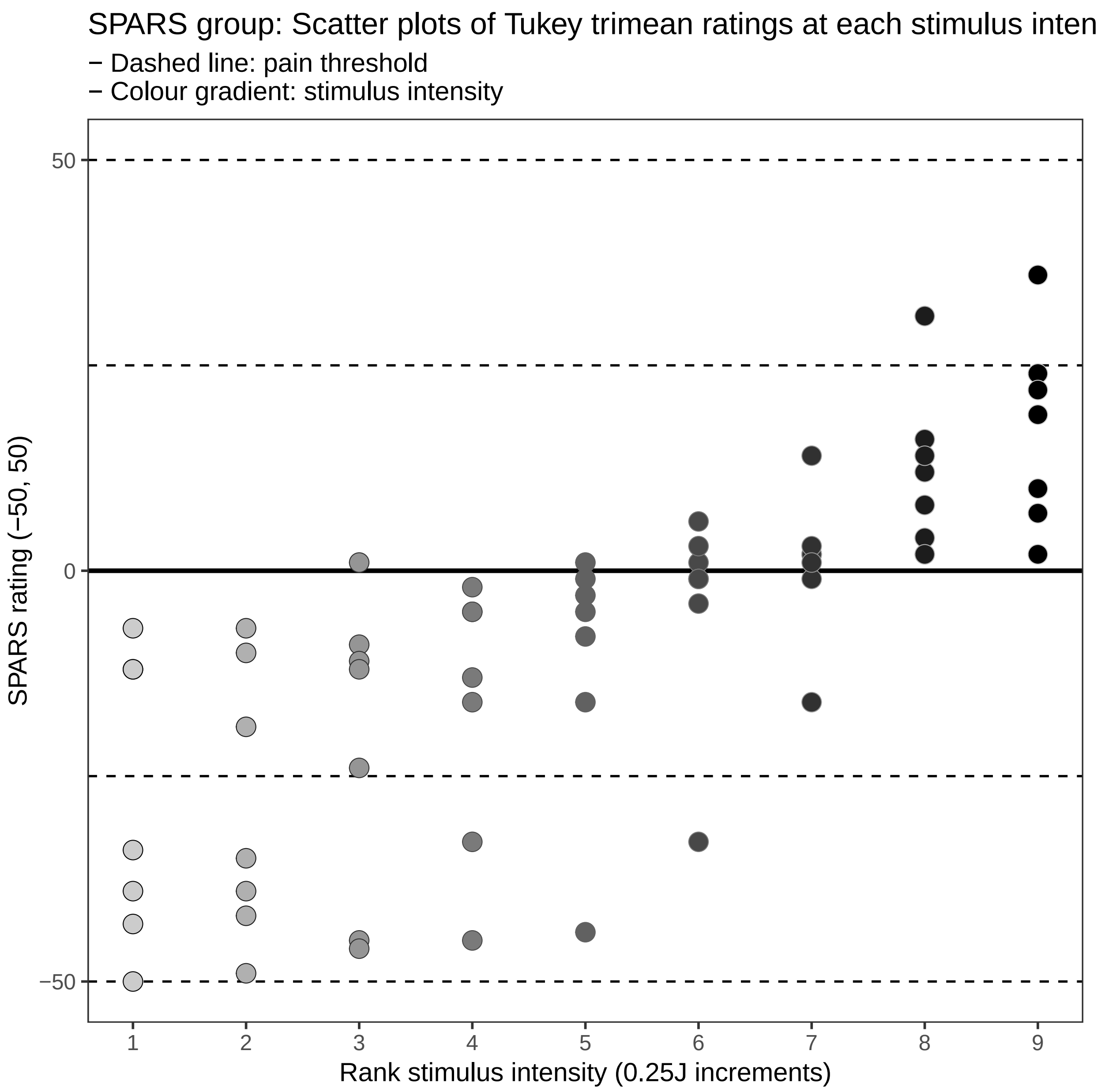

##### NRS

```
# Plot scatter plot of ratings for the group at every intensity
ggplot(data = group_nrs) +
    aes(x = intensity,
        y = tri_mean,
        fill = intensity,
        colour = intensity) +
    geom_hline(yintercept = 0,
               size = 1) +
    geom_hline(yintercept = 25,
               linetype = 2) +
    geom_hline(yintercept = 75,
               linetype = 2) +
```

```r
geom_hline(yintercept = 100,
           linetype = 2) +
geom_point(shape = 21,
           size = 4,
           stroke = 0.3) +
scale_fill_gradient(low = '#CCCCCC', high = '#000000') +
scale_colour_gradient(low = '#000000', high = '#CCCCCC') +
scale_y_continuous(limits = c(0, 100),
                   breaks = c(0, 50, 100)) +
scale_x_continuous(breaks = seq(from = 1,
                                to = 9,
                                by = 1)) +
labs(title = "NRS group: Scatter plots of Tukey trimean ratings at each stimulus intensity",
     subtitle = '- Dashed line: pain threshold\n- Colour gradient: stimulus intensity',
     x = 'Rank stimulus intensity (0.25J increments)',
     y = 'NRS rating (0, 100)') +
theme(legend.position = 'none',
      panel.grid = element_blank())
```

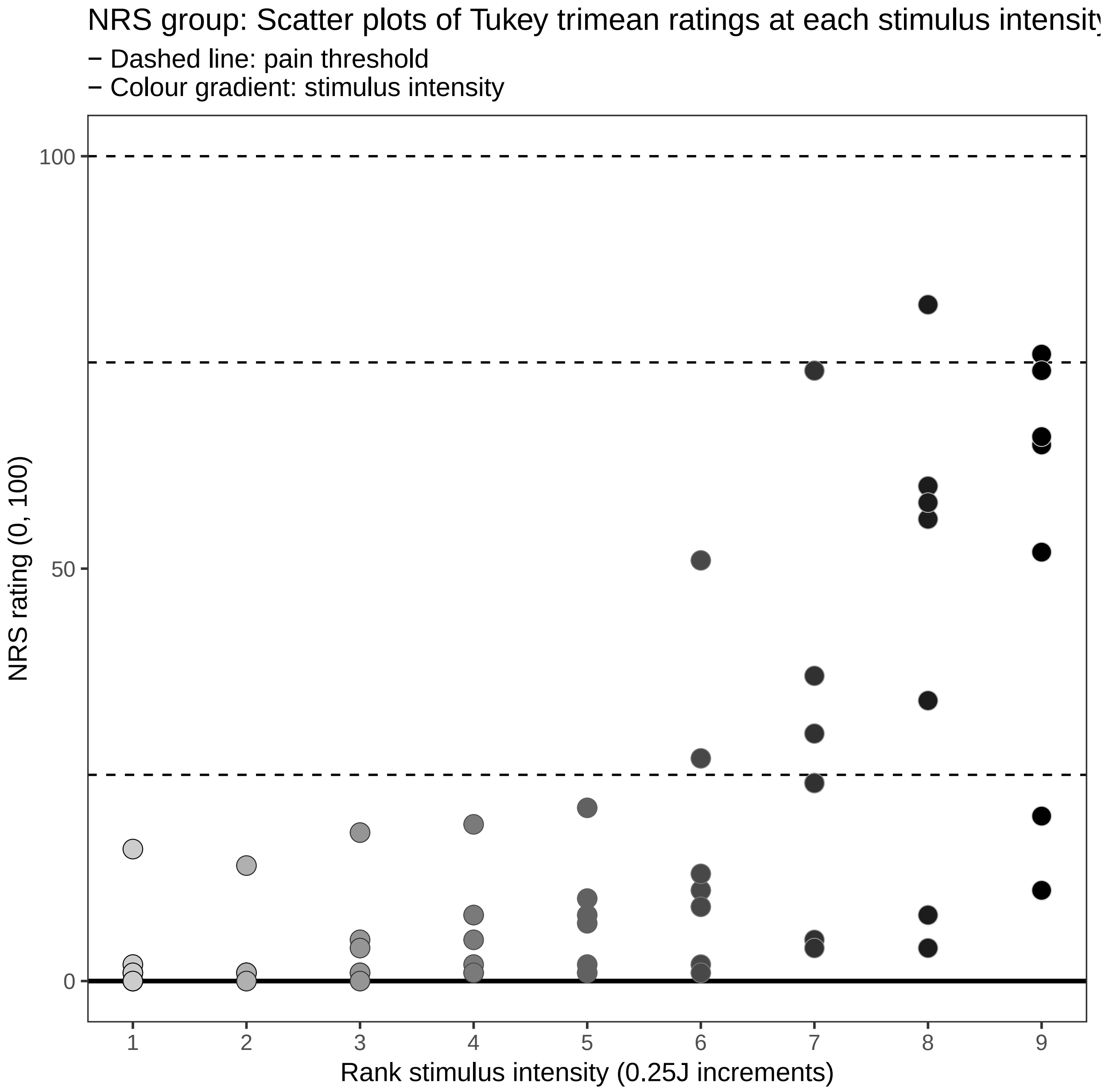

##### SRS

```
# Plot scatter plot of ratings for the group at every intensity
ggplot(data = group_srs) +
    aes(x = intensity,
        y = tri_mean,
        fill = intensity,
        colour = intensity) +
    geom_hline(yintercept = 0,
               size = 1) +
    geom_hline(yintercept = -25,
               linetype = 2) +
    geom_hline(yintercept = -75,
               linetype = 2) +
```

```r
geom_hline(yintercept = -100,
           linetype = 2) +
geom_point(shape = 21,
           size = 4,
           stroke = 0.3) +
scale_fill_gradient(low = '#CCCCCC', high = '#000000') +
scale_colour_gradient(low = '#000000', high = '#CCCCCC') +
scale_y_continuous(limits = c(-100, 0),
                   breaks = c(-100, -50, 0)) +
scale_x_continuous(breaks = seq(from = 1,
                                to = 9,
                                by = 1)) +
labs(title = "SRS group: Scatter plots of Tukey trimean ratings at each stimulus intensity",
     subtitle = '- Dashed line: pain threshold\n- Colour gradient: stimulus intensity',
     x = 'Rank stimulus intensity (0.25J increments)',
     y = 'SRS rating (-100, 0)') +
theme(legend.position = 'none',
      panel.grid = element_blank())
```

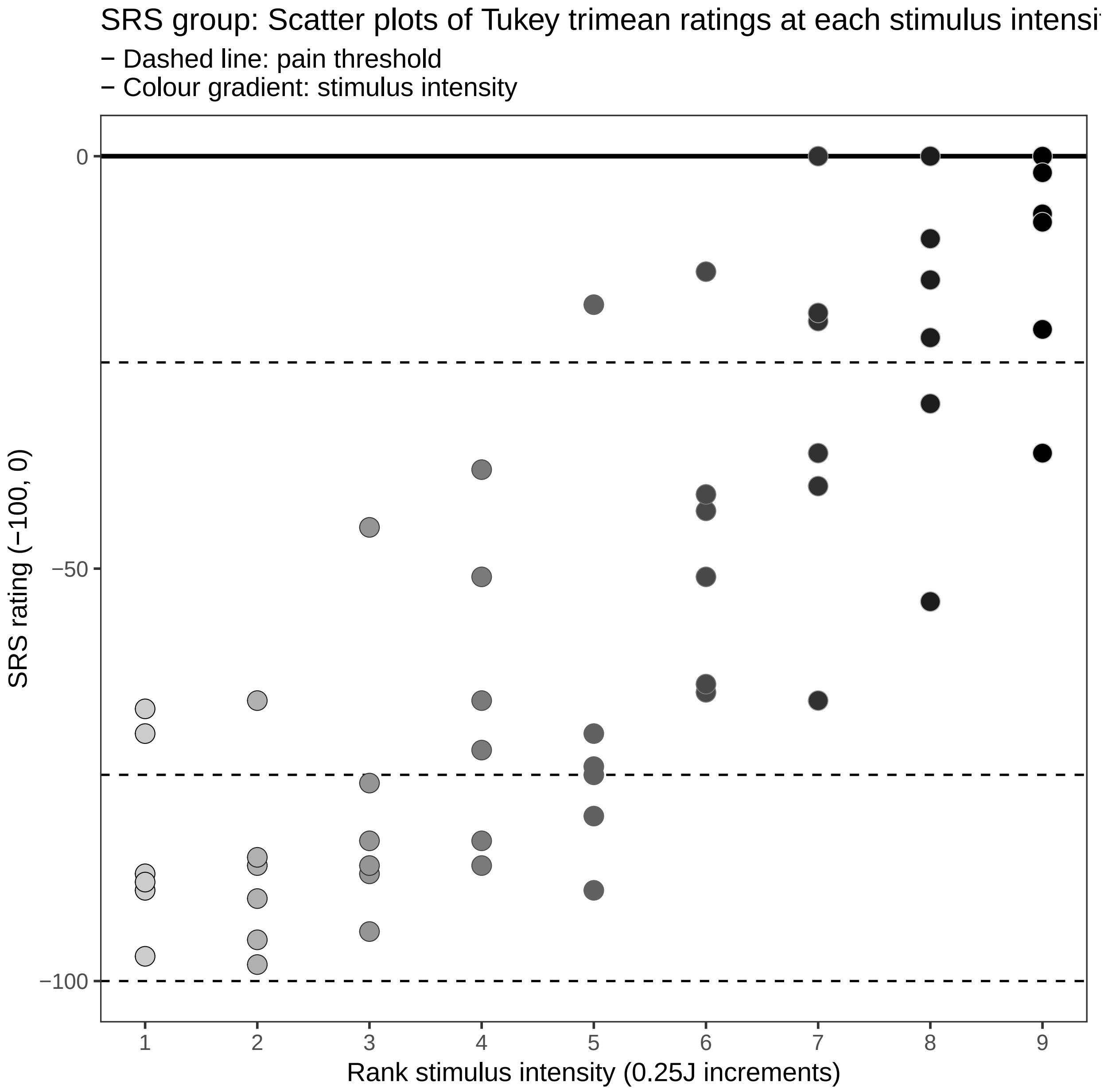

**Trimean confidence interval plots**

**SPARS**

```
# Plot group CIs at every intensity
ggplot(data = spars_boot_group) +
    aes(x = intensity) +
    geom_hline(yintercept = 0,
               size = 1) +
    geom_hline(yintercept = 25,
               linetype = 2) +
    geom_hline(yintercept = -25,
               linetype = 2) +
    geom_hline(yintercept = 50,
```

```
           linetype = 2) +
geom_hline(yintercept = -50,
           linetype = 2) +
geom_crossbar(aes(y = tri_mean,
                  ymin = lower_ci,
                  ymax = upper_ci,
                  fill = fill,
                  colour = fill),
              fatten = 0,
              size = 0.3) +
scale_fill_manual(values = c('#000000', '#CCCCCC')) +
scale_colour_manual(values = c('#CCCCCC', '#000000')) +
scale_y_continuous(limits = c(-50, 50),
                   breaks = c(-50, 0, 50)) +
scale_x_continuous(breaks = seq(from = 1,
                                to = 9,
                                by = 1)) +
labs(title = "SPARS Group: Crossbar plots of 95% CI of Tukey trimeans\nfor ratings at each stimulus
     subtitle = '- Basic bootstrap 95% CI with 10,000 resamples\n- Dashed line: pain threshold | - I
     x = 'Rank stimulus intensity (0.25J increments)',
     y = 'SPARS rating (-50, 50)') +
theme(legend.position = 'none',
      panel.grid = element_blank())
```

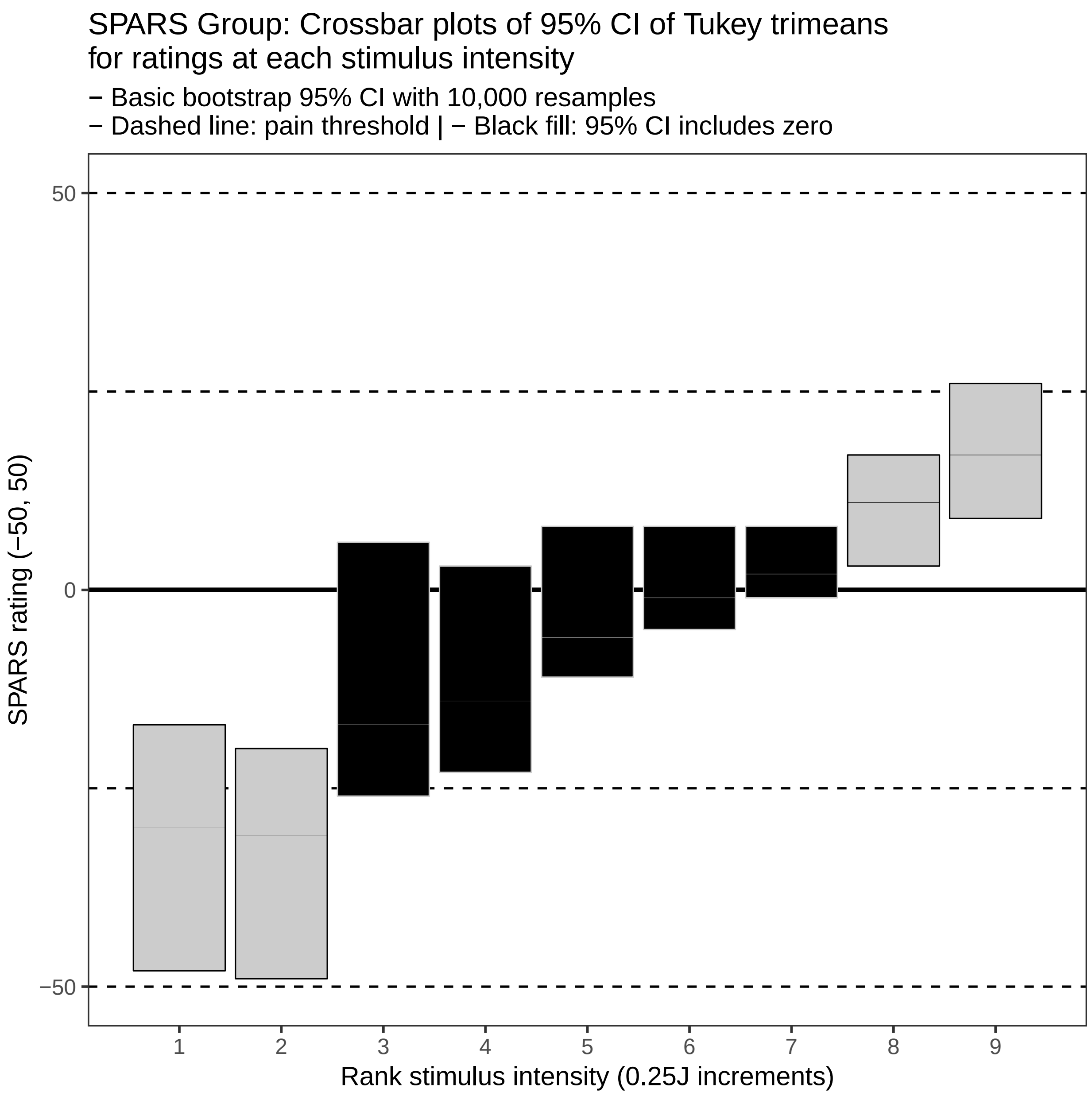

##### NRS

```
# Plot group CIs at every intensity
ggplot(data = nrs_boot_group) +
    aes(x = intensity) +
    geom_hline(yintercept = 0,
               size = 1) +
    geom_hline(yintercept = 25,
               linetype = 2) +
    geom_hline(yintercept = 50,
               linetype = 2) +
    geom_hline(yintercept = 75,
               linetype = 2) +
    geom_hline(yintercept = 100,
```

```
            linetype = 2) +
geom_crossbar(aes(y = tri_mean,
                  ymin = lower_ci,
                  ymax = upper_ci,
                  fill = fill,
                  colour = fill),
              fatten = 0,
              size = 0.3) +
scale_fill_manual(values = c('#000000', '#CCCCCC')) +
scale_colour_manual(values = c('#CCCCCC', '#000000')) +
scale_y_continuous(limits = c(0, 100),
                   breaks = c(0, 50, 100)) +
scale_x_continuous(breaks = seq(from = 1,
                                to = 9,
                                by = 1)) +
labs(title = "NRS Group: Crossbar plots of 95% CI of Tukey trimeans\nfor ratings at each stimulus i
     subtitle = '- Basic bootstrap 95% CI with 10,000 resamples\n- Dashed line: pain threshold | - 
     x = 'Rank stimulus intensity (0.25J increments)',
     y = 'NRS rating (0, 100)') +
theme(legend.position = 'none',
      panel.grid = element_blank())
```

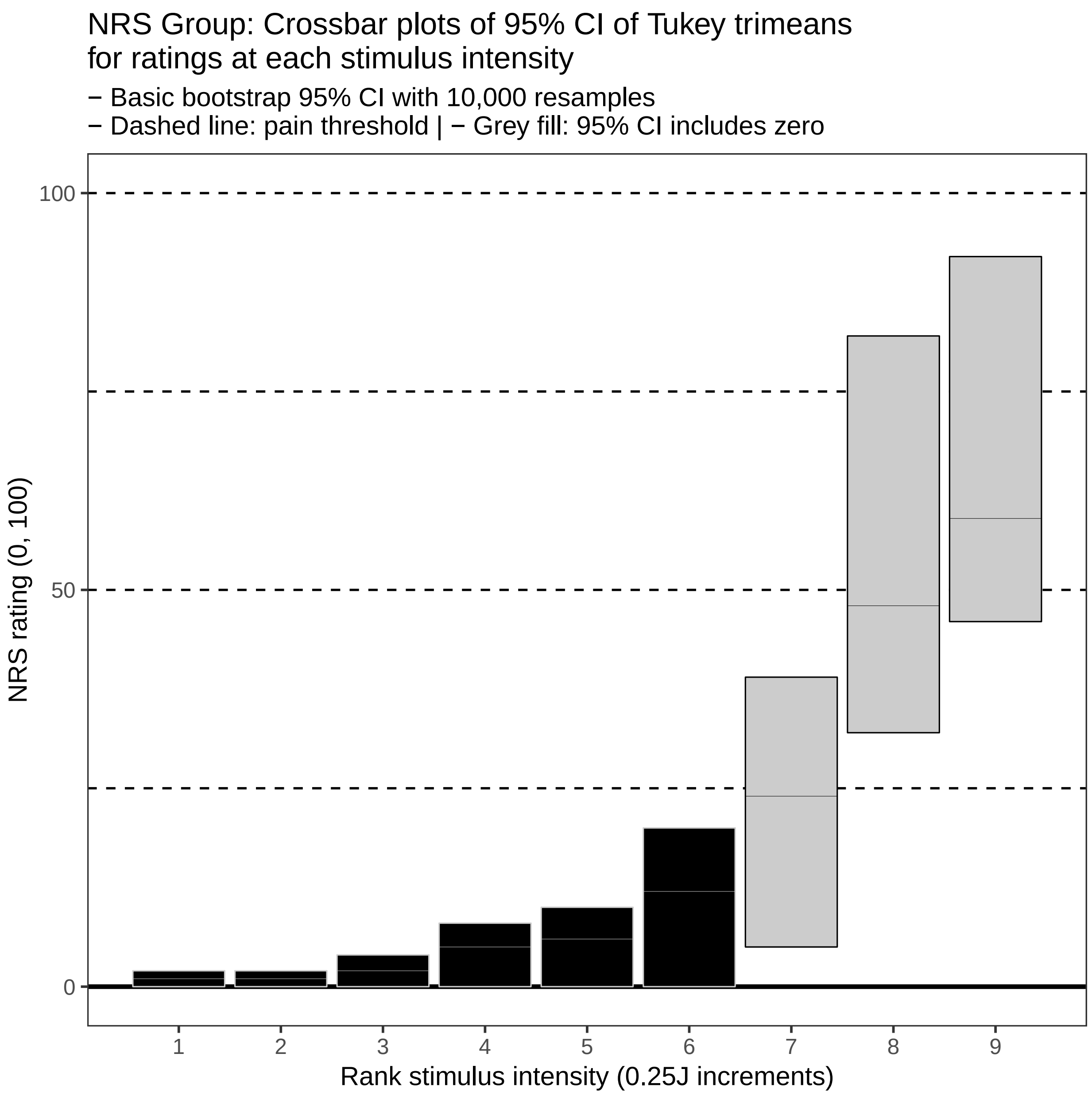

##### SRS

```
# Plot group CIs at every intensity
ggplot(data = srs_boot_group) +
    aes(x = intensity) +
    geom_hline(yintercept = 0,
               size = 1) +
    geom_hline(yintercept = -25,
               linetype = 2) +
    geom_hline(yintercept = -50,
               linetype = 2) +
    geom_hline(yintercept = -75,
               linetype = 2) +
    geom_hline(yintercept = -100,
```

```
           linetype = 2) +
geom_crossbar(aes(y = tri_mean,
                  ymin = lower_ci,
                  ymax = upper_ci,
                  fill = fill,
                  colour = fill),
              fatten = 0,
              size = 0.3) +
scale_fill_manual(values = c('#000000', '#CCCCCC')) +
scale_colour_manual(values = c('#CCCCCC', '#000000')) +
scale_y_continuous(limits = c(-100, 0),
                   breaks = c(-100, -50, 0)) +
scale_x_continuous(breaks = seq(from = 1,
                                to = 9,
                                by = 1)) +
labs(title = "SRS Group: Crossbar plots of 95% CI of Tukey trimeans\nfor ratings at each stimulus i
     subtitle = '- Basic bootstrap 95% CI with 10,000 resamples\n- Dashed line: pain threshold | - (
     x = 'Stimulus intensity (0.25J increments)',
     y = 'SRS rating (-100, 0)') +
theme(legend.position = 'none',
      panel.grid = element_blank())
```

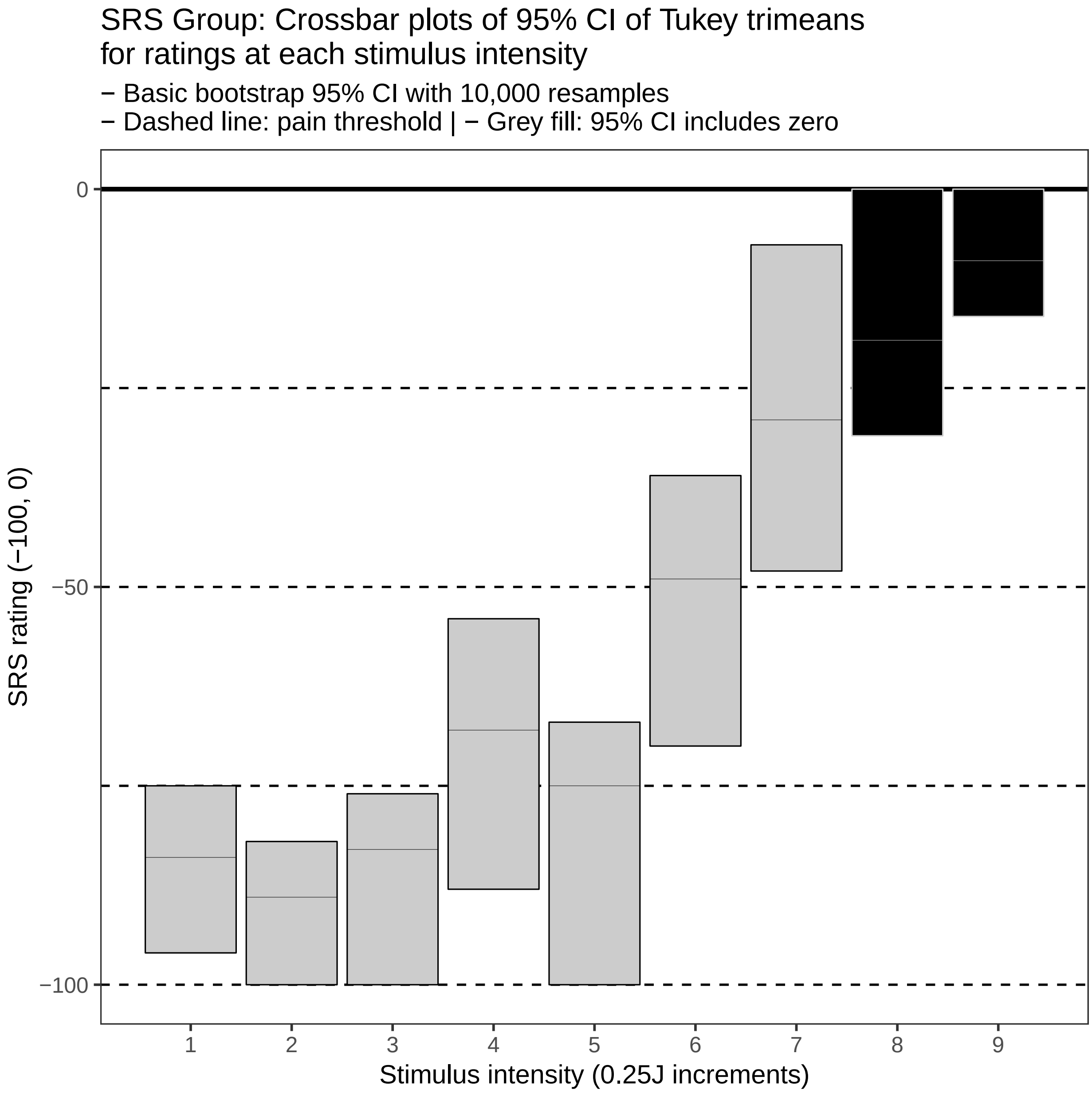

---

#### Session information

```r
sessionInfo()
```

```
## R version 3.5.1 (2018-07-02)
## Platform: x86_64-pc-linux-gnu (64-bit)
## Running under: Debian GNU/Linux 9 (stretch)
##
## Matrix products: default
## BLAS: /usr/lib/openblas-base/libblas.so.3
## LAPACK: /usr/lib/libopenblasp-r0.2.19.so
##
```

```
## locale:
##  [1] LC_CTYPE=en_US.UTF-8       LC_NUMERIC=C
##  [3] LC_TIME=en_US.UTF-8        LC_COLLATE=en_US.UTF-8
##  [5] LC_MONETARY=en_US.UTF-8    LC_MESSAGES=C
##  [7] LC_PAPER=en_US.UTF-8       LC_NAME=C
##  [9] LC_ADDRESS=C               LC_TELEPHONE=C
## [11] LC_MEASUREMENT=en_US.UTF-8 LC_IDENTIFICATION=C
##
## attached base packages:
## [1] stats     graphics  grDevices utils     datasets  methods   base
##
## other attached packages:
##  [1] bindrcpp_0.2.2  skimr_1.0.3     boot_1.3-20     magrittr_1.5
##  [5] forcats_0.3.0   stringr_1.3.1   dplyr_0.7.8     purrr_0.2.5
##  [9] readr_1.3.0     tidyr_0.8.2     tibble_1.4.2    ggplot2_3.1.0
## [13] tidyverse_1.2.1
##
## loaded via a namespace (and not attached):
##  [1] tidyselect_0.2.5 xfun_0.4         haven_2.0.0      lattice_0.20-35
##  [5] colorspace_1.3-2 generics_0.0.2   htmltools_0.3.6  yaml_2.2.0
##  [9] rlang_0.3.0.1    pillar_1.3.1     glue_1.3.0       withr_2.1.2
## [13] modelr_0.1.2     readxl_1.2.0     bindr_0.1.1      plyr_1.8.4
## [17] munsell_0.5.0    gtable_0.2.0     cellranger_1.1.0 rvest_0.3.2
## [21] evaluate_0.12    knitr_1.21       broom_0.5.1      Rcpp_1.0.0
## [25] scales_1.0.0     backports_1.1.3  jsonlite_1.6     hms_0.4.2
## [29] digest_0.6.18    stringi_1.2.4    grid_3.5.1       cli_1.0.1
## [33] tools_3.5.1      lazyeval_0.2.1   crayon_1.3.4     pkgconfig_2.0.2
## [37] xml2_1.2.0       lubridate_1.7.4  assertthat_0.2.0 rmarkdown_1.11
## [41] httr_1.4.0       rstudioapi_0.8   R6_2.3.0         nlme_3.1-137
## [45] compiler_3.5.1
```
