## Supplementary File 3 for "Rethinking pain threshold as a zone of uncertainty"

Binomial test analysis: Does the distribution of ratings differ from a theoretical 'NULL' distribution?

*12 Jan 2019*

**Session information** 25

---

#### Question

For each participant, and at each stimulus intensity, does the distribution of SPARS, NRS, or SRS ratings differ significantly from a theoretical 'NULL' distribution?

To answer the question, we used the binomial test. The binomial test is useful when there are two possible outcomes because it assesses whether the distribution of the observed data deviates from a distribution that would theoretically be expected to occur by chance. Using the test required that we dichotomize the continuous rating data we collected (generally not a good thing, but here it is appropriate).

The SPARS ranges from -50 ('no sensation') to +50 ('most intense pain you can imagine'), and therefore ratings can span 0 (pain threshold, 'the exact point at which you feel transitions to pain'). We therefore coded SPARS ratings $< 0$ as being **'negative'**, and ratings $> 0$ as being **'positive'**. In the first SPARS experiment (SPARS A), participants were not allowed to record a stimulus as 0 , but in the second SPARS experiment (SPARS B), they could record stimuli as 0 on the scale. We felt that the 0 ratings in the SPARS B experiment were uninformative, and so we excluded ratings of 0 from the analysis (we describe the number of zero ratings per participant below). The NRS ranges from 0 ('no pain') to 100 ('most intense pain you can imagine'), and therefore ratings immediately to the right of the 0-point of the scale mark the transition from non-painful to painful sensation. We therefore coded NRS ratings $= 0$ as being **'negative'**, and ratings $> 0$ as being **'positive'**. In addition, it has been reported that individuals use the first 15 points of a 0 to 100 NRS to record non-painful stimuli (Littman et al, 1985), and so we also analysed the NRS data with NRS ratings $\leq 15$ as being **'negative'**, and ratings $>15$ as being **'positive'**.

The SRS ranges from -100 ('no sensation') to 0 ('just painful/pain threshold'), and therefore ratings immediately to the left of the 0-point of the scale mark the transition from non-painful to painful sensation. We therefore coded SRS ratings $= 0$ as being **'positive'**, and ratings $< 0$ as being **'negative'**.

In all cases, we modelled the data using the binomial test with a 50% probability of 'success' (positive rating arbitrarily chosen as success). This is a conservative approach as one would expect that for the SPARS and the NRS, as stimulus intensity increases above pain threshold, the probability of recording a 'positive' rating increases. Similarly, in the case of the SPARS (which allows the rating of intensity of noxious and non-noxious stimuli), one would expect that the probability of recording a 'negative' rating would increase as stimulus intensity decreased. However, since we did not know the approximate intensity of a theshold stimulus, and there was high inter-individual variation in sensitivity, we were unable to gage at which stimulus intensities we should start shifting the probability of 'success' away from 50% (see Supplement 1 and Supplement 2).

Because ratings on the SPARS can range from -50 to +50, we analysed the data using a two-tailed p-value. That is, the distribution may shift to the left or right of the theoretical distribution. However, because the NRS has a floor rating of 0 ('no pain') and the SRS has a ceiling rating of 0 ('pain threshold'), the change in rating from 0 is unidirectional ($> 0$), so we performed the binomial test with a one-tailed p-value. For all test, significance was assessed at the $\alpha = 0.05$ level. And, because this was an exploratory analysis, we did not make any family-wide corrections for multiple comparisons.

---

### SPARS A

#### Import and inspect data

```
# Import
data_sparsA <- read_rds('data-cleaned/SPARS_A.rds')

# Inspect
glimpse(data_sparsA)
```

```
## Observations: 1,927
## Variables: 6
## $ PID          <chr> "ID01", "ID01", "ID01", "ID01", "ID01", "ID01", "...
## $ block        <chr> "A", "A", "A", "A", "A", "A", "A", "A", "A", "A",...
## $ block_order  <dbl> 4, 4, 4, 4, 4, 4, 4, 4, 4, 4, 4, 4, 4, 4, 4, 4, 4...
## $ trial_number <dbl> 79, 80, 81, 82, 83, 84, 85, 86, 87, 88, 89, 90, 9...
## $ intensity    <dbl> 3.00, 2.25, 4.00, 3.25, 2.75, 2.25, 2.75, 4.00, 2...
## $ rating       <dbl> -40, -25, 10, 2, -10, -25, -20, 10, -25, -50, -25...
```

```r
data_sparsA %>%
    select(intensity, rating) %>%
    skim()
```

```
## Skim summary statistics
##  n obs: 1927
##  n variables: 2
##
## -- Variable type:numeric -----------------------------------------------------------
##   variable missing complete    n  mean    sd  p0    p25 p50   p75 p100
##  intensity       0     1927 1927  2.47  0.93   1   1.75 2.5  3.25    4
##     rating       0     1927 1927 -4.45 22.31 -50 -20    2   10      45
##      hist
##
##
```

##### Binomial test

```r
# Select columns
data_sparsA %<>%
    select(PID, intensity, rating)

# Nest data by PID and stimulus intensity
sparsA_nest <- data_sparsA %>%
    group_by(PID, intensity) %>%
    nest()

# Generate data
sparsA_nest %<>%
    # Add probability of success column
    mutate(prob = 0.5) %>%
    # Extract rating data from dataframe
    mutate(data_vec = map(.x = data,
                      ~ .$rating)) %>%
    # Recode rating data as categories according to sign
    mutate(data_cat = map(.x = data_vec,
                      ~ ifelse(.x < 0,
                               yes = 'negative',
                               no = 'positive'))) %>%
    # Count the number of positive and negative ratings
    ## positive numbers arbitrarily listed first == 'success'
    mutate(success_count = map(.x = data_cat,
                       ~ c(length(.x[.x == 'positive']),
                           length(.x[.x == 'negative'])))) %>%
    # Conduct binomial test (two-sided)
    mutate(binomial_test = map2(.x = success_count,
                            .y = prob,
                            ~ binom.test(x = .x,
                                         p = .y,
                                         alternative = 'two.sided'))) %>%
    # Extract p-value from binomial_test
    mutate(binomial_p.value = map(.x = binomial_test,
                              ~ .x$p.value %>%
                                  round(., 3))) %>%
    # Categorise p-value using a p < 0.05 threshold
```

```
    ## Significant: distribution deviates significantly
    ## from the theoretical distribution
    ## No correction for multiple comparisons
    ## (too conservative for explorartory analysis)
    mutate(significant_p.value = map(.x = binomial_p.value,
                                     ~ ifelse(.x < 0.05,
                                              yes = 'yes',
                                              no = 'no')))
```

##### Plot

For each paticipant, we plotted raw SPARS ratings at each stimulus intensity and colour-coded the data according to whether the p-value returned by the binomial test was significant (distribution of data points deviates significantly from the theoretical expected distribution).

```
sparsA_nest %>%
    # Select data columns
    select(PID, intensity, significant_p.value) %>%
    # Unnest data
    unnest() %>%
    # Join with original data
    right_join(data_sparsA) %>%
    # Reclass intensity as an ordered factor
    mutate(intensity = factor(intensity,
                              ordered = TRUE)) %>%
    # Plot
    ggplot(data = .) +
    aes(x = intensity,
        y = rating,
        fill = significant_p.value,
        colour = significant_p.value) +
    geom_hline(yintercept = 0,
               size = 1) +
    geom_hline(yintercept = 25,
               linetype = 2) +
    geom_hline(yintercept = -25,
               linetype = 2) +
    geom_hline(yintercept = 50,
               linetype = 2) +
    geom_hline(yintercept = -50,
               linetype = 2) +
    geom_point(shape = 21,
               size = 4,
               stroke = 0.3) +
    labs(title = "SPARS A: Binomial test of positive/negative rating distribution",
         subtitle = "Probability of 'success' = 0.5* | alpha = 0.05 | two-tailed p-value\nFilled circle
         caption = "* 'success' arbitrarily chosen as a positive SPARS ratings",
         x = 'Stimulus intensity (J)',
         y = 'SPARS rating (-50, 50)') +
    scale_x_discrete(breaks = seq(from = 1,
                                  to = 4,
                                  by = 0.5),
                     labels = sprintf('%.2f', seq(from = 1,
                                                  to = 4,
                                                  by = 0.5))) +
```

```
scale_y_continuous(limits = c(-50, 50),
                   breaks = c(-50, 0, 50),
                   labels = c(-50, 0, 50)) +
scale_fill_manual(values = c('#000000', '#CCCCCC')) +
scale_colour_manual(values = c('#CCCCCC', '#000000')) +
facet_wrap(~ PID, ncol = 4) +
theme(legend.position = 'none',
      panel.grid = element_blank(),
      panel.spacing = unit(0.1, 'lines'),
      strip.text = element_text(margin = margin(t = 0.1,
                                                 b = 0.1,
                                                 r = 1,
                                                 l = 1,
                                                 'lines')),
      axis.text.x = element_text(angle = -90,
                                 vjust = 0.5))
```

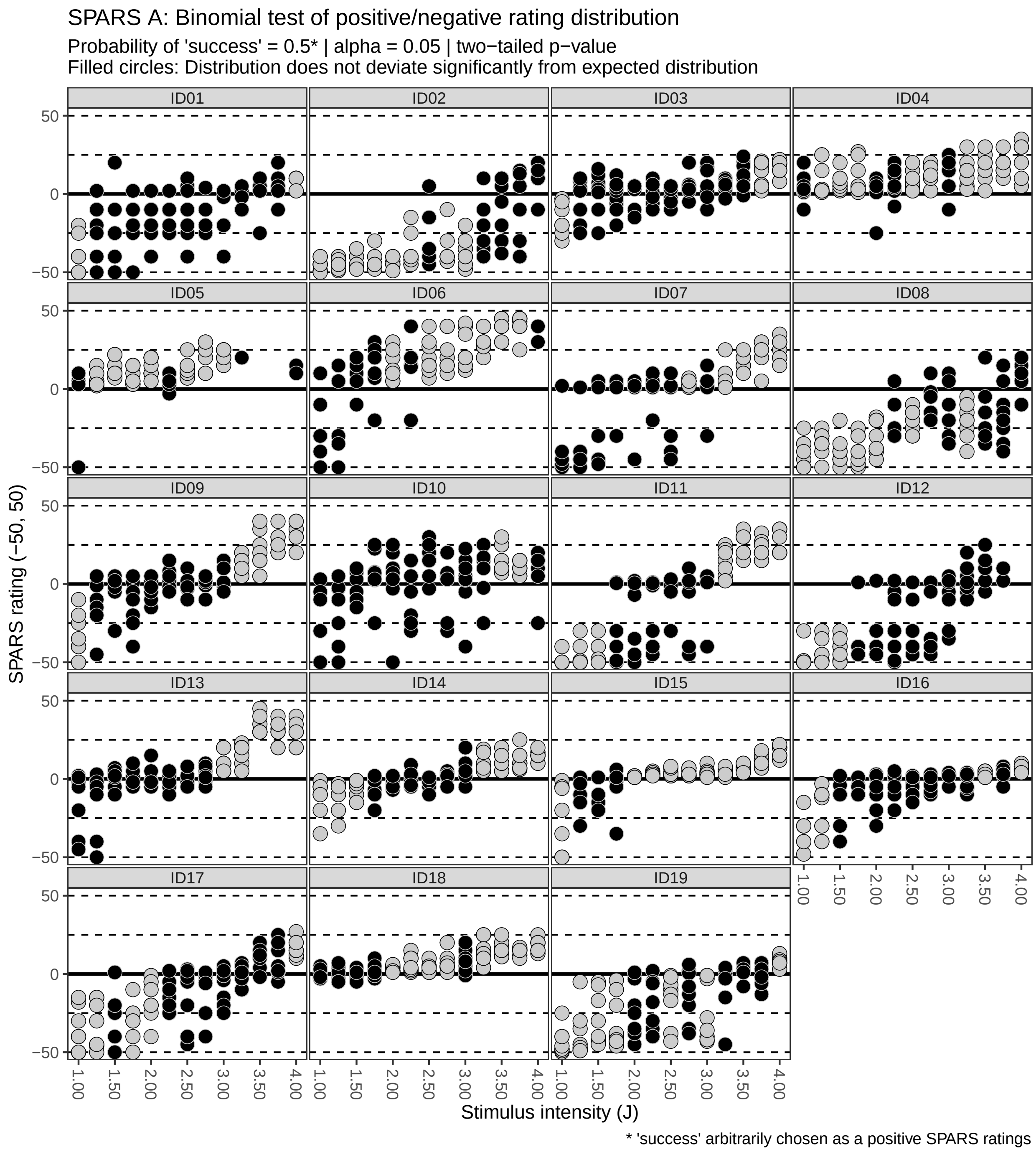

---

### SPARS B

#### Import and inspect data

```
# Import
data_sparsB <- read_rds('data-cleaned/SPARS_B.rds') %>%
    # Extract trials rated using the SPARS
```

```
    filter(scale == 'SPARS') %>%
    # Remove <NA>
    filter(!is.na(rating))

# Rank stimulus intensity
data_sparsB %<>%
    group_by(PID, scale) %>%
    arrange(intensity) %>%
    mutate(intensity_rank = dense_rank(intensity)) %>%
    select(-intensity) %>%
    rename(intensity = intensity_rank) %>%
    ungroup()

# Inspect
glimpse(data_sparsB)
```

```
## Observations: 752
## Variables: 6
## $ PID          <chr> "ID06", "ID06", "ID06", "ID06", "ID06", "ID06", "...
## $ block_number <int> 1, 1, 1, 2, 2, 2, 3, 3, 3, 4, 4, 4, 1, 1, 1, 2, 2...
## $ trial_number <dbl> 4, 6, 27, 9, 13, 20, 20, 24, 27, 4, 18, 22, 5, 16...
## $ scale        <chr> "SPARS", "SPARS", "SPARS", "SPARS", "SPARS", "SPA...
## $ rating       <dbl> -49, 2, -6, 3, -20, -2, -31, 2, -5, -8, -23, 14, ...
## $ intensity    <int> 1, 1, 1, 1, 1, 1, 1, 1, 1, 1, 1, 1, 2, 2, 2, 2, 2...
```

```
data_sparsB %>%
    select(intensity, rating) %>%
    skim()
```

```
## Skim summary statistics
##  n obs: 752
##  n variables: 2
##
## -- Variable type:integer -----------------------------------------------------------
##   variable missing complete   n mean   sd p0 p25 p50 p75 p100     hist
##  intensity       0      752 752    5 2.58  1   3   5   7    9
##
## -- Variable type:numeric -----------------------------------------------------------
##  variable missing complete   n  mean    sd  p0 p25 p50 p75 p100     hist
##    rating       0      752 752 -8.83 23.46 -50 -26  -4   5   50
```

```
# Number of 0 ratings
data_sparsB %>%
    # Retain ratings of 0
    filter(rating == 0) %>%
    # Select columns
    select(PID, intensity, rating) %>%
    # Group by individual and intensity
    group_by(PID, intensity) %>%
    # Summarise
    summarise(zero_count = n()) %>%
    ftable(.)
```

```
##                 zero_count 1 2 3 4
## PID  intensity
## ID01 1                     0 0 0 0
```

```
##      2                    0 0 0 0
##      3                    0 0 0 0
##      5                    0 0 0 1
##      6                    1 0 0 0
##      7                    0 0 1 0
##      8                    0 0 0 0
##      9                    0 0 0 0
## ID02 1                    1 0 0 0
##      2                    0 0 0 0
##      3                    0 0 0 0
##      5                    0 0 0 0
##      6                    1 0 0 0
##      7                    0 0 0 0
##      8                    0 0 0 0
##      9                    0 0 0 0
## ID03 1                    0 0 0 0
##      2                    1 0 0 0
##      3                    0 0 0 0
##      5                    0 0 0 0
##      6                    0 0 0 0
##      7                    0 1 0 0
##      8                    0 0 0 0
##      9                    0 0 0 0
## ID04 1                    0 0 0 0
##      2                    0 0 0 0
##      3                    0 0 0 0
##      5                    0 0 0 0
##      6                    0 1 0 0
##      7                    0 0 0 0
##      8                    0 0 0 0
##      9                    0 0 0 0
## ID06 1                    0 0 0 0
##      2                    1 0 0 0
##      3                    1 0 0 0
##      5                    0 1 0 0
##      6                    0 1 0 0
##      7                    1 0 0 0
##      8                    1 0 0 0
##      9                    0 0 0 0
## ID07 1                    0 0 0 0
##      2                    1 0 0 0
##      3                    1 0 0 0
##      5                    1 0 0 0
##      6                    0 1 0 0
##      7                    0 0 1 0
##      8                    0 1 0 0
##      9                    1 0 0 0
```

#### Binomial test

```
# Select data
data_sparsB %<>%
    # Remove ratings of 0
    filter(rating != 0) %>%
```

```r
    # Select columns
    select(PID, intensity, rating)

# Nest data by PID and stimulus intensity
sparsB_nest <- data_sparsB %>%
    group_by(PID, intensity) %>%
    nest()

# Generate data
sparsB_nest %<>%
    # Add probability of success column
    mutate(prob = 0.5) %>%
    # Extract rating data from dataframe
    mutate(data_vec = map(.x = data,
                          ~ .$rating)) %>%
    # Recode rating data as categories according to sign
    mutate(data_cat = map(.x = data_vec,
                          ~ ifelse(.x < 0,
                                   yes = 'negative',
                                   no = 'positive'))) %>%
    # Count the number of positive and negative ratings
    ## positive numbers arbitrarily listed first == 'success'
    mutate(success_count = map(.x = data_cat,
                               ~ c(length(.x[.x == 'positive']),
                                   length(.x[.x == 'negative'])))) %>%
    # Conduct binomial test (two-sided)
    mutate(binomial_test = map2(.x = success_count,
                                .y = prob,
                                ~ binom.test(x = .x,
                                             p = .y,
                                             alternative = 'two.sided'))) %>%
    # Extract p-value from binomial_test
    mutate(binomial_p.value = map(.x = binomial_test,
                                  ~ .x$p.value %>%
                                    round(., 3))) %>%
    # Categorise p-value using a p < 0.05 threshold
    ## Significant: distribution deviates significantly
    ## from the theoretical distribution
    ## No correction for multiple comparisons
    ## (too conservative for explorartory analysis)
    mutate(significant_p.value = map(.x = binomial_p.value,
                                     ~ ifelse(.x < 0.05,
                                              yes = 'yes',
                                              no = 'no')))
```

#### Plot

```r
sparsB_nest %>%
    # Select data columns
    select(PID, intensity, significant_p.value) %>%
    # Unnest data
    unnest() %>%
    # Join with original data
    right_join(data_sparsB) %>%
```

```r
# Reclass intensity as an ordered factor
mutate(intensity = factor(intensity,
                          ordered = TRUE)) %>%
# Plot
ggplot(data = .) +
aes(x = intensity,
    y = rating,
    fill = significant_p.value,
    colour = significant_p.value) +
geom_hline(yintercept = 0,
           size = 1) +
geom_hline(yintercept = 25,
           linetype = 2) +
geom_hline(yintercept = -25,
           linetype = 2) +
geom_hline(yintercept = 50,
           linetype = 2) +
geom_hline(yintercept = -50,
           linetype = 2) +
geom_point(shape = 21,
           size = 4,
           stroke = 0.3) +
labs(title = "SPARS B: Binomial test of positive/negative rating distribution",
     subtitle = "Probability of 'success' = 0.5* | alpha = 0.05 | two-tailed p-value\nFilled circle
     caption = "* 'success' arbitrarily chosen as a positive SPARS ratings",
     x = 'Rank stimulus intensity (0.25J increments)',
     y = 'SPARS rating (-50, 50)') +
scale_x_discrete(breaks = seq(from = 1,
                              to = 9,
                              by = 1),
                 labels = sprintf('%.0f', seq(from = 1,
                                              to = 9,
                                              by = 1))) +
scale_y_continuous(limits = c(-50, 50),
                   breaks = c(-50, 0, 50),
                   labels = c(-50, 0, 50)) +
scale_fill_manual(values = c('#000000', '#CCCCCC')) +
scale_colour_manual(values = c('#CCCCCC', '#000000')) +
facet_wrap(~ PID, ncol = 4) +
theme(legend.position = 'none',
      panel.grid = element_blank(),
      panel.spacing = unit(0.1, 'lines'),
      strip.text = element_text(margin = margin(t = 0.1,
                                                b = 0.1,
                                                r = 1,
                                                l = 1,
                                                'lines')))
```

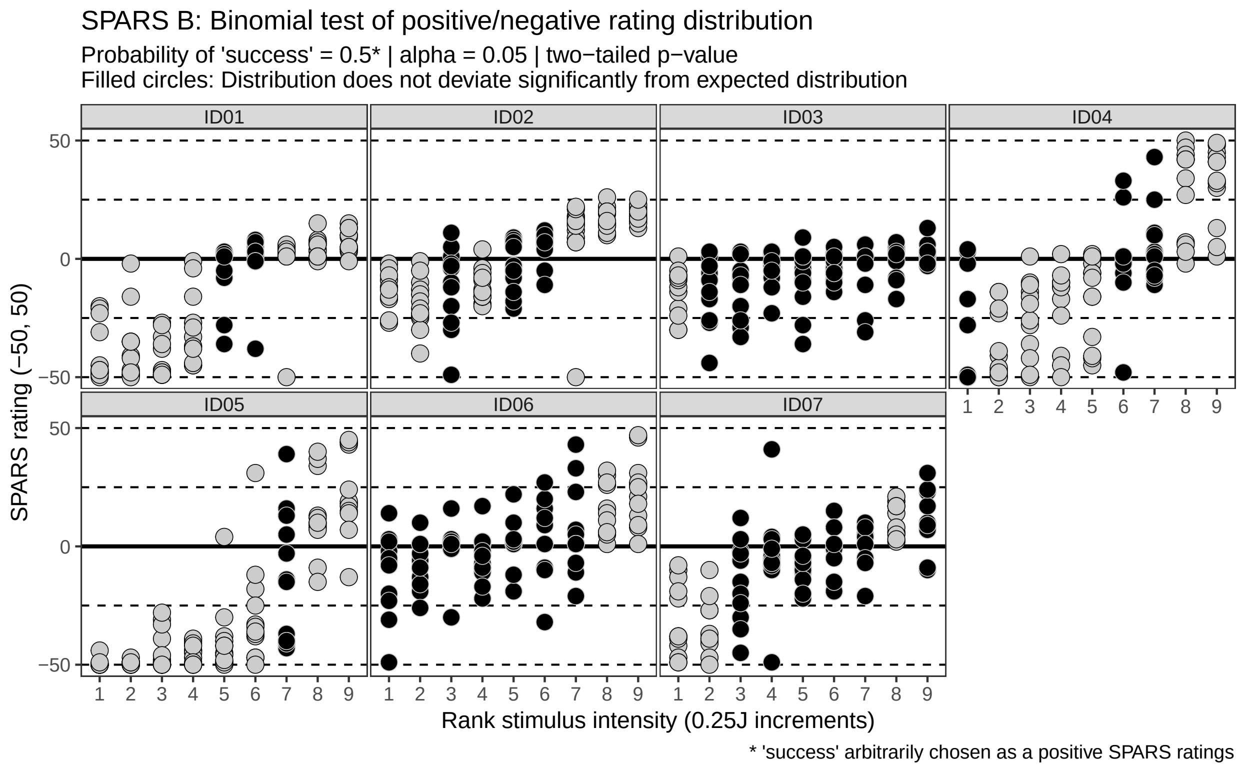

---

### NRS (zero: 0)

#### Import and inspect data

```
# Import
data_nrs <- read_rds('data-cleaned/SPARS_B.rds') %>%
    # Extract trials rated using the SPARS
    filter(scale == 'NRS') %>%
    # Remove <NA>
    filter(!is.na(rating))

# Rank stimulus intensity
data_nrs %<>%
    group_by(PID, scale) %>%
    arrange(intensity) %>%
    mutate(intensity_rank = dense_rank(intensity)) %>%
    select(-intensity) %>%
    rename(intensity = intensity_rank) %>%
    ungroup()

# Inspect
glimpse(data_nrs)
```

```
## Observations: 753
## Variables: 6
## $ PID          <chr> "ID06", "ID06", "ID06", "ID06", "ID06", "ID06", "...
## $ block_number <int> 9, 9, 9, 10, 10, 10, 11, 11, 11, 12, 12, 12, 9, 9...
```

```
## $ trial_number <dbl> 7, 9, 26, 4, 9, 27, 2, 4, 12, 4, 7, 10, 5, 6, 27,...
## $ scale        <chr> "NRS", "NRS", "NRS", "NRS", "NRS", "NRS", "NRS", ...
## $ rating       <dbl> 5, 2, 0, 0, 0, 0, 0, 1, 0, 0, 0, 48, 1, 0, 53, 0,...
## $ intensity    <int> 1, 1, 1, 1, 1, 1, 1, 1, 1, 1, 1, 1, 2, 2, 2, 2, 2...
```

```
data_nrs %>%
    select(intensity, rating) %>%
    skim()
```

```
## Skim summary statistics
##  n obs: 753
##  n variables: 2
##
## -- Variable type:integer -----------------------------------------------------------
##   variable missing complete   n mean   sd p0 p25 p50 p75 p100     hist
##  intensity       0      753 753    5 2.59  1   3   5   7    9
##
## -- Variable type:numeric -----------------------------------------------------------
##  variable missing complete   n  mean    sd p0 p25 p50 p75 p100     hist
##    rating       0      753 753 19.63 26.82  0   1   5  28   98
```

```
# Number of 0 ratings
data_nrs %>%
    # Retain ratings of 0
    filter(rating == 0) %>%
    # Select columns
    select(PID, intensity, rating) %>%
    # Group by individual and intensity
    group_by(PID, intensity) %>%
    # Summarise
    summarise(zero_count = n()) %>%
    ftable(.)
```

```
##                zero_count 1 2 3 4 5 6 7 8 10 12
## PID  intensity
## ID01 1                    0 0 0 0 0 0 0 0  0  1
##      2                    0 0 0 0 0 0 0 0  1  0
##      3                    0 0 0 0 0 0 0 0  1  0
##      4                    0 0 0 1 0 0 0 0  0  0
##      5                    0 0 0 1 0 0 0 0  0  0
##      6                    0 0 0 0 0 0 0 0  0  0
##      7                    0 1 0 0 0 0 0 0  0  0
## ID02 1                    0 0 0 0 0 0 0 0  1  0
##      2                    0 0 0 0 0 0 1 0  0  0
##      3                    0 0 1 0 0 0 0 0  0  0
##      4                    0 0 0 0 0 0 0 0  0  0
##      5                    0 0 0 0 0 0 0 0  0  0
##      6                    0 0 0 0 0 0 0 0  0  0
##      7                    0 0 0 0 0 0 0 0  0  0
## ID03 1                    0 0 0 0 1 0 0 0  0  0
##      2                    0 0 0 1 0 0 0 0  0  0
##      3                    0 0 1 0 0 0 0 0  0  0
##      4                    0 0 0 0 0 0 0 0  0  0
##      5                    0 1 0 0 0 0 0 0  0  0
##      6                    1 0 0 0 0 0 0 0  0  0
##      7                    1 0 0 0 0 0 0 0  0  0
```

```
## ID04 1                    0 0 0 0 0 1 0 0  0  0
##      2                    0 0 0 0 0 1 0 0  0  0
##      3                    0 0 0 1 0 0 0 0  0  0
##      4                    0 0 1 0 0 0 0 0  0  0
##      5                    1 0 0 0 0 0 0 0  0  0
##      6                    1 0 0 0 0 0 0 0  0  0
##      7                    1 0 0 0 0 0 0 0  0  0
## ID05 1                    0 0 0 1 0 0 0 0  0  0
##      2                    0 0 0 1 0 0 0 0  0  0
##      3                    0 0 0 0 0 0 1 0  0  0
##      4                    0 1 0 0 0 0 0 0  0  0
##      5                    0 1 0 0 0 0 0 0  0  0
##      6                    0 0 0 0 0 0 0 0  0  0
##      7                    0 0 0 0 0 0 0 0  0  0
## ID06 1                    0 0 0 0 0 0 0 1  0  0
##      2                    0 0 0 0 0 0 0 1  0  0
##      3                    0 0 0 0 0 0 0 1  0  0
##      4                    0 0 0 0 0 0 1 0  0  0
##      5                    0 0 0 0 0 1 0 0  0  0
##      6                    0 1 0 0 0 0 0 0  0  0
##      7                    0 0 0 0 0 0 0 0  0  0
```

#### Binomial test

```
# Select data
data_nrs %<>%
    # Select columns
    select(PID, intensity, rating)

# Nest data by PID and stimulus intensity
nrs_nest <- data_nrs %>%
    group_by(PID, intensity) %>%
    nest()

# Generate data
nrs0_nest <- nrs_nest %>%
    # Add probability of success column
    mutate(prob = 0.5) %>%
    # Extract rating data from dataframe
    mutate(data_vec = map(.x = data,
                          ~ .$rating)) %>%
    # Recode rating data as categories according to whether
    # the value is greater than 0 (minimum rating on NRS)
    mutate(data_cat = map(.x = data_vec,
                          ~ ifelse(.x == 0,
                                   yes = 'negative',
                                   no = 'positive'))) %>%
    # Count the number of positive and negative ratings
    ## positive numbers arbitrarily listed first == 'success'
    mutate(success_count = map(.x = data_cat,
                           ~ c(length(.x[.x == 'positive']),
                               length(.x[.x == 'negative'])))) %>%
    # Conduct binomial test (two-sided)
    mutate(binomial_test = map2(.x = success_count,
```

```
                                .y = prob,
                                ~ binom.test(x = .x,
                                             p = .y,
                                             alternative = 'greater'))) %>%
    # Extract p-value from binomial_test
    mutate(binomial_p.value = map(.x = binomial_test,
                                  ~ .x$p.value %>%
                                      round(., 3))) %>%
    # Categorise p-value using a p < 0.05 threshold
    ## Significant: distribution deviates significantly
    ## from the theoretical distribution
    ## No correction for multiple comparisons
    ## (too conservative for explorartory analysis)
    mutate(significant_p.value = map(.x = binomial_p.value,
                                      ~ ifelse(.x < 0.05,
                                               yes = 'yes',
                                               no = 'no')))
```

#### Plot

```
nrs0_nest %>%
    # Select data columns
    select(PID, intensity, significant_p.value) %>%
    # Unnest data
    unnest() %>%
    # Join with original data
    right_join(data_nrs) %>%
    # Reclass intensity as an ordered factor
    mutate(intensity = factor(intensity,
                              ordered = TRUE)) %>%
    # Plot
    ggplot(data = .) +
    aes(x = intensity,
        y = rating,
        fill = significant_p.value,
        colour = significant_p.value) +
    geom_hline(yintercept = 0,
               size = 1) +
    geom_hline(yintercept = 25,
               linetype = 2) +
    geom_hline(yintercept = 50,
               linetype = 2) +
    geom_hline(yintercept = 75,
               linetype = 2) +
    geom_hline(yintercept = 100,
               linetype = 2) +
    geom_point(shape = 21,
               size = 4,
               stroke = 0.3) +
    labs(title = "NRS (0): Binomial test of positive/negative rating distribution",
         subtitle = "Probability of 'success' = 0.5* | alpha = 0.05 | two-tailed p-value\nFilled circle
         caption = "* 'success' arbitrarily chosen as NRS rating > 0",
         x = 'Rank stimulus intensity (0.25J increments)',
         y = 'NRS rating (0, 100)') +
```

```
scale_x_discrete(breaks = seq(from = 1,
                              to = 9,
                              by = 1),
                 labels = sprintf('%.0f', seq(from = 1,
                                              to = 9,
                                              by = 1))) +
scale_y_continuous(limits = c(0, 100),
                   breaks = c(0, 50, 100),
                   labels = c(0, 50, 100)) +
scale_fill_manual(values = c('#000000', '#CCCCCC')) +
scale_colour_manual(values = c('#CCCCCC', '#000000')) +
facet_wrap(~ PID, ncol = 4) +
theme(legend.position = 'none',
      panel.grid = element_blank(),
      panel.spacing = unit(0.1, 'lines'),
      strip.text = element_text(margin = margin(t = 0.1,
                                                b = 0.1,
                                                r = 1,
                                                l = 1,
                                                'lines')))
```

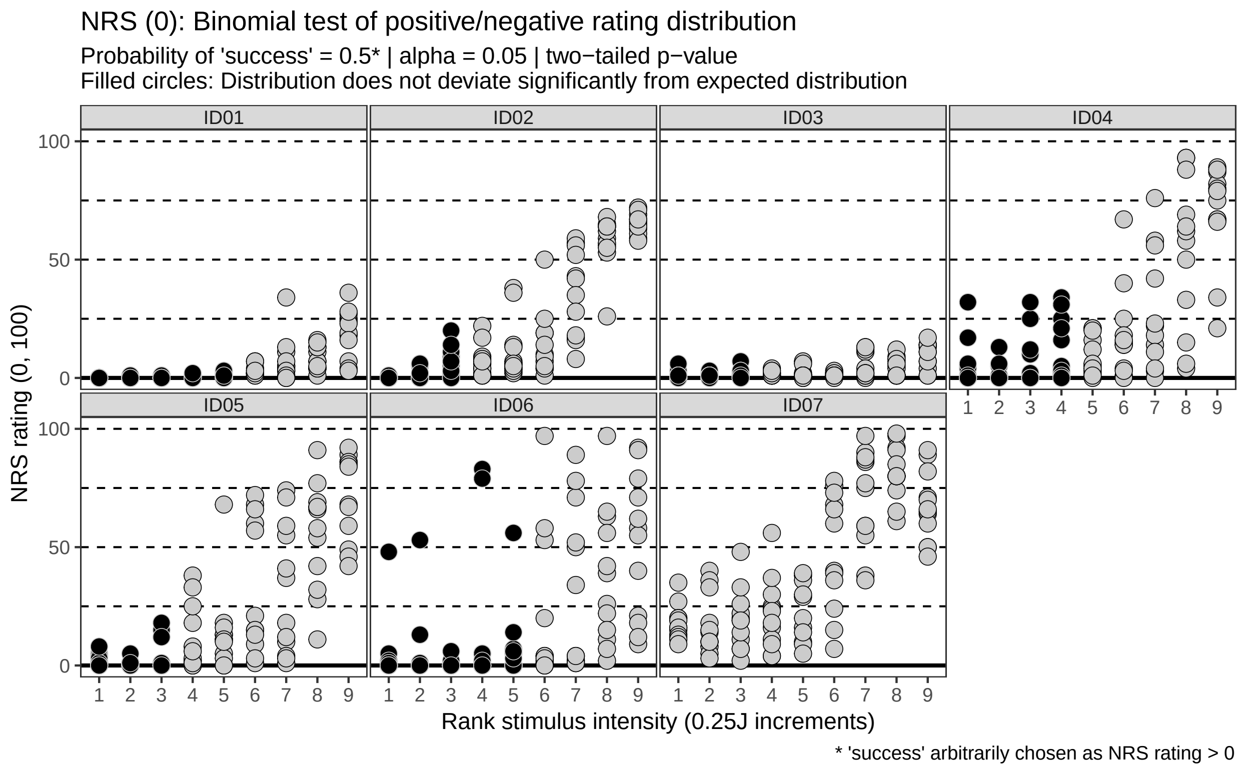

---

### NRS (zero: 0 to 15)

#### Import and inspect data

Data already imported, inspected, and nested *(data_nrs, nrs_nest)*.

#### Binomial test

```
# Generate data
nrs15_nest <- nrs_nest %>%
    # Add probability of success column
    mutate(prob = 0.5) %>%
    # Extract rating data from dataframe
    mutate(data_vec = map(.x = data,
                          ~ .$rating)) %>%
    # Recode rating data as categories according to whether
    # the value is greater than 0 (minimum rating on NRS)
    mutate(data_cat = map(.x = data_vec,
                          ~ ifelse(.x <= 15,
                                   yes = 'negative',
                                   no = 'positive'))) %>%
    # Count the number of positive and negative ratings
    ## positive numbers arbitrarily listed first == 'success'
    mutate(success_count = map(.x = data_cat,
                               ~ c(length(.x[.x == 'positive']),
                                   length(.x[.x == 'negative'])))) %>%
    # Conduct binomial test (two-sided)
    mutate(binomial_test = map2(.x = success_count,
                                .y = prob,
                                ~ binom.test(x = .x,
                                             p = .y,
                                             alternative = 'greater'))) %>%
    # Extract p-value from binomial_test
    mutate(binomial_p.value = map(.x = binomial_test,
                                  ~ .x$p.value %>%
                                      round(., 3))) %>%
    # Categorise p-value using a p < 0.05 threshold
    ## Significant: distribution deviates significantly
    ## from the theoretical distribution
    ## No correction for multiple comparisons
    ## (too conservative for explorartory analysis)
    mutate(significant_p.value = map(.x = binomial_p.value,
                                     ~ ifelse(.x < 0.05,
                                              yes = 'yes',
                                              no = 'no')))
```

#### Plot

```
nrs15_nest %>%
    # Select data columns
    select(PID, intensity, significant_p.value) %>%
    # Unnest data
    unnest() %>%
    # Join with original data
    right_join(data_nrs) %>%
    # Reclass intensity as an ordered factor
    mutate(intensity = factor(intensity,
                              ordered = TRUE)) %>%
    # Plot
    ggplot(data = .) +
    aes(x = intensity,
```

```
    y = rating,
    fill = significant_p.value,
    colour = significant_p.value) +
geom_rect(aes(ymin = 0, ymax = 15,
              xmin = -1, xmax = 10),
          fill = '#666666',
          colour = '#666666') +
geom_hline(yintercept = 0,
           size = 1) +
geom_hline(yintercept = 25,
           linetype = 2) +
geom_hline(yintercept = 50,
           linetype = 2) +
geom_hline(yintercept = 75,
           linetype = 2) +
geom_hline(yintercept = 100,
           linetype = 2) +
geom_point(shape = 21,
           size = 4,
           stroke = 0.3) +
labs(title = "NRS (0-15): Binomial test of positive/negative rating distribution",
     subtitle = "Probability of 'success' = 0.5* | alpha = 0.05 | two-tailed p-value\nFilled circle
     caption = "* 'success' arbitrarily chosen as NRS rating > 15",
     x = 'Rank stimulus intensity (0.25J increments)',
     y = 'NRS rating (0, 100)') +
scale_x_discrete(breaks = seq(from = 1,
                              to = 9,
                              by = 1),
                 labels = sprintf('%.0f', seq(from = 1,
                                              to = 9,
                                              by = 1))) +
scale_y_continuous(limits = c(0, 100),
                   breaks = c(0, 50, 100),
                   labels = c(0, 50, 100)) +
scale_fill_manual(values = c('#000000', '#CCCCCC')) +
scale_colour_manual(values = c('#CCCCCC', '#000000')) +
facet_wrap(~ PID, ncol = 4) +
theme(legend.position = 'none',
      panel.grid = element_blank(),
      panel.spacing = unit(0.1, 'lines'),
      strip.text = element_text(margin = margin(t = 0.1,
                                                b = 0.1,
                                                r = 1,
                                                l = 1,
                                                'lines')))
```

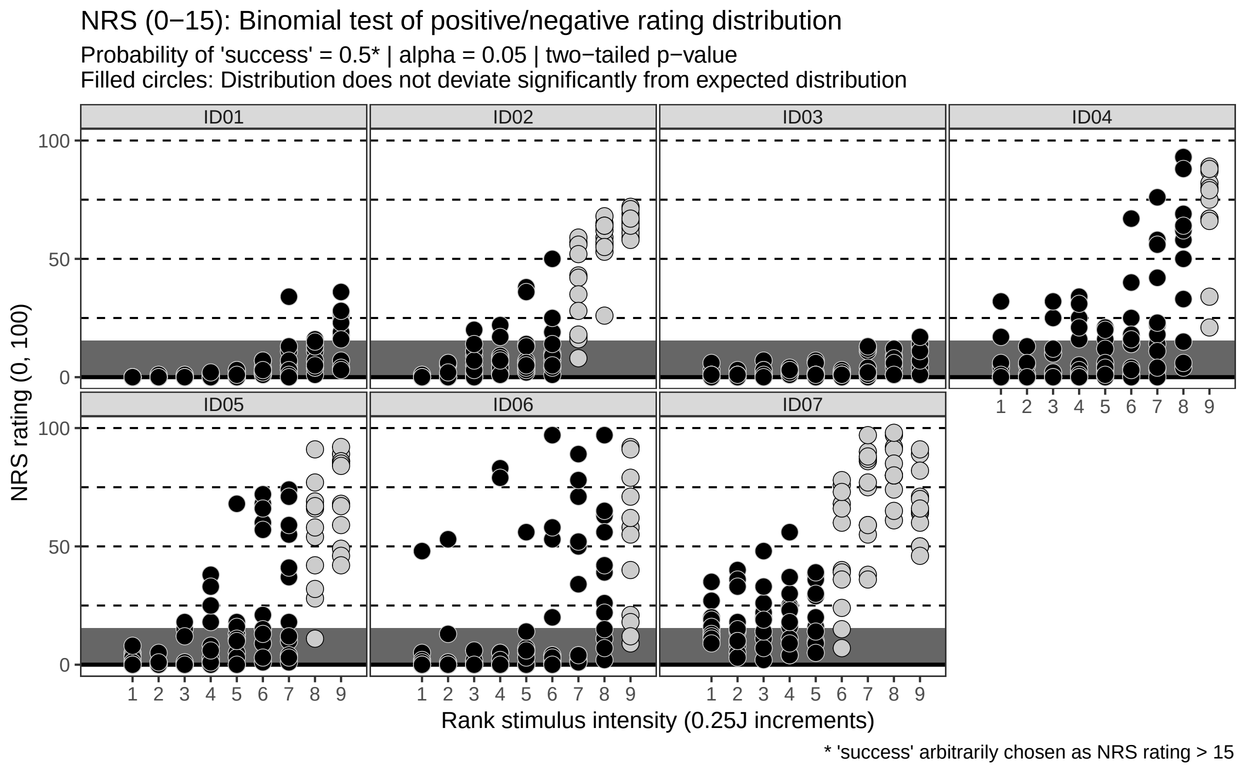

---

### SRS (zero: 0)

#### Import and inspect data

```
# Import
data_srs <- read_rds('data-cleaned/SPARS_B.rds') %>%
    # Extract trials rated using the SRS
    filter(scale == 'SRS') %>%
    # Remove <NA>
    filter(!is.na(rating))

# Rank stimulus intensity
data_srs %<>%
    group_by(PID, scale) %>%
    arrange(intensity) %>%
    mutate(intensity_rank = dense_rank(intensity)) %>%
    select(-intensity) %>%
    rename(intensity = intensity_rank) %>%
    ungroup()

# Inspect
glimpse(data_srs)
```

```
## Observations: 644
## Variables: 6
## $ PID          <chr> "ID06", "ID06", "ID06", "ID06", "ID06", "ID06", "...
## $ block_number <int> 5, 5, 5, 6, 6, 6, 7, 7, 7, 8, 8, 8, 5, 5, 5, 6, 6...
```

```
## $ trial_number <dbl> 2, 16, 26, 13, 19, 21, 1, 17, 27, 2, 4, 5, 6, 24,...
## $ scale        <chr> "SRS", "SRS", "SRS", "SRS", "SRS", "SRS", "SRS", ...
## $ rating       <dbl> -34, -99, -89, -99, -100, -99, -59, -96, -70, -92...
## $ intensity    <int> 1, 1, 1, 1, 1, 1, 1, 1, 1, 1, 1, 1, 2, 2, 2, 2, 2...
```

```
data_srs %>%
    select(intensity, rating) %>%
    skim()
```

```
## Skim summary statistics
##  n obs: 644
##  n variables: 2
##
## -- Variable type:integer ------------------------------------------------------------
##   variable missing complete   n mean   sd p0 p25 p50 p75 p100     hist
##  intensity       0      644 644    5 2.58  1   3   5   7    9
##
## -- Variable type:numeric ------------------------------------------------------------
##  variable missing complete   n   mean    sd   p0 p25 p50 p75 p100     hist
##    rating       0      644 644 -54.46 35.19 -100 -88 -63 -21    0
```

```
# Number of 0 ratings
data_srs %>%
    # Retain ratings of 0
    filter(rating == 0) %>%
    # Select columns
    select(PID, intensity, rating) %>%
    # Group by individual and intensity
    group_by(PID, intensity) %>%
    # Summarise
    summarise(zero_count = n()) %>%
    ftable(.)
```

```
##                zero_count 1 2 3 5 9 12
## PID  intensity
## ID02 5                    0 1 0 0 0  0
##      6                    0 0 0 1 0  0
##      7                    0 0 0 0 1  0
##      8                    0 0 0 0 0  1
##      9                    0 0 0 0 0  1
## ID04 5                    0 0 0 0 0  0
##      6                    0 0 0 0 0  0
##      7                    0 0 0 0 0  0
##      8                    1 0 0 0 0  0
##      9                    1 0 0 0 0  0
## ID05 5                    0 0 0 0 0  0
##      6                    0 0 0 0 0  0
##      7                    0 0 0 0 0  0
##      8                    0 0 0 0 0  0
##      9                    1 0 0 0 0  0
## ID06 5                    0 0 0 0 0  0
##      6                    0 0 0 0 0  0
##      7                    1 0 0 0 0  0
##      8                    1 0 0 0 0  0
##      9                    0 0 1 0 0  0
```

#### Binomial test

```
# Select data
data_srs %<>%
    # Select columns
    select(PID, intensity, rating)

# Nest data by PID and stimulus intensity
srs_nest <- data_srs %>%
    group_by(PID, intensity) %>%
    nest()

# Generate data
srs_nest <- srs_nest %>%
    # Add probability of success column
    mutate(prob = 0.5) %>%
    # Extract rating data from dataframe
    mutate(data_vec = map(.x = data,
                          ~ .$rating)) %>%
    # Recode rating data as categories according to whether
    # the value is less than 0 (maximum rating on srs)
    mutate(data_cat = map(.x = data_vec,
                          ~ ifelse(.x == 0,
                                   yes = 'positive',
                                   no = 'negative'))) %>%
    # Count the number of positive and negative ratings
    ## positive numbers arbitrarily listed first == 'success'
    mutate(success_count = map(.x = data_cat,
                           ~ c(length(.x[.x == 'positive']),
                               length(.x[.x == 'negative'])))) %>%
    # Conduct binomial test (two-sided)
    mutate(binomial_test = map2(.x = success_count,
                                .y = prob,
                                ~ binom.test(x = .x,
                                             p = .y,
                                             alternative = 'greater'))) %>%
    # Extract p-value from binomial_test
    mutate(binomial_p.value = map(.x = binomial_test,
                                  ~ .x$p.value %>%
                                      round(., 3))) %>%
    # Categorise p-value using a p < 0.05 threshold
    ## Significant: distribution deviates significantly
    ## from the theoretical distribution
    ## No correction for multiple comparisons
    ## (too conservative for explorartory analysis)
    mutate(significant_p.value = map(.x = binomial_p.value,
                                     ~ ifelse(.x < 0.05,
                                              yes = 'yes',
                                              no = 'no')))
```

#### Plot

```
srs_nest %>%
    # Select data columns
    select(PID, intensity, significant_p.value) %>%
```

```r
# Unnest data
unnest() %>%
# Join with original data
right_join(data_srs) %>%
# Reclass intensity as an ordered factor
mutate(intensity = factor(intensity,
                          ordered = TRUE)) %>%
# Plot
ggplot(data = .) +
aes(x = intensity,
    y = rating,
    fill = significant_p.value,
    colour = significant_p.value) +
geom_hline(yintercept = 0,
           size = 1) +
geom_hline(yintercept = -25,
           linetype = 2) +
geom_hline(yintercept = -50,
           linetype = 2) +
geom_hline(yintercept = -75,
           linetype = 2) +
geom_hline(yintercept = -100,
           linetype = 2) +
geom_point(shape = 21,
           size = 4,
           stroke = 0.3) +
scale_fill_manual(values = c('#CCCCCC', '#000000')) +
scale_colour_manual(values = c('#000000', '#CCCCCC')) +
labs(title = "SRS: Binomial test of positive/negative rating distribution",
     subtitle = "Probability of 'success' = 0.5* | alpha = 0.05 | two-tailed p-value\nFilled circle
     caption = "* 'success' arbitrarily chosen as SRS rating = 0",
     x = 'Rank stimulus intensity (0.25J increments)',
     y = 'SRS rating (-100, 0)') +
scale_x_discrete(breaks = seq(from = 1,
                              to = 9,
                              by = 1),
                 labels = sprintf('%.0f', seq(from = 1,
                                              to = 9,
                                              by = 1))) +
scale_y_continuous(limits = c(-100, 0),
                   breaks = c(-100, -50, 0),
                   labels = c(-100, -50, 0)) +
facet_wrap(~ PID, ncol = 4) +
theme(legend.position = 'none',
      panel.grid = element_blank(),
      panel.spacing = unit(0.1, 'lines'),
      strip.text = element_text(margin = margin(t = 0.1,
                                                b = 0.1,
                                                r = 1,
                                                l = 1,
                                                'lines')))
```

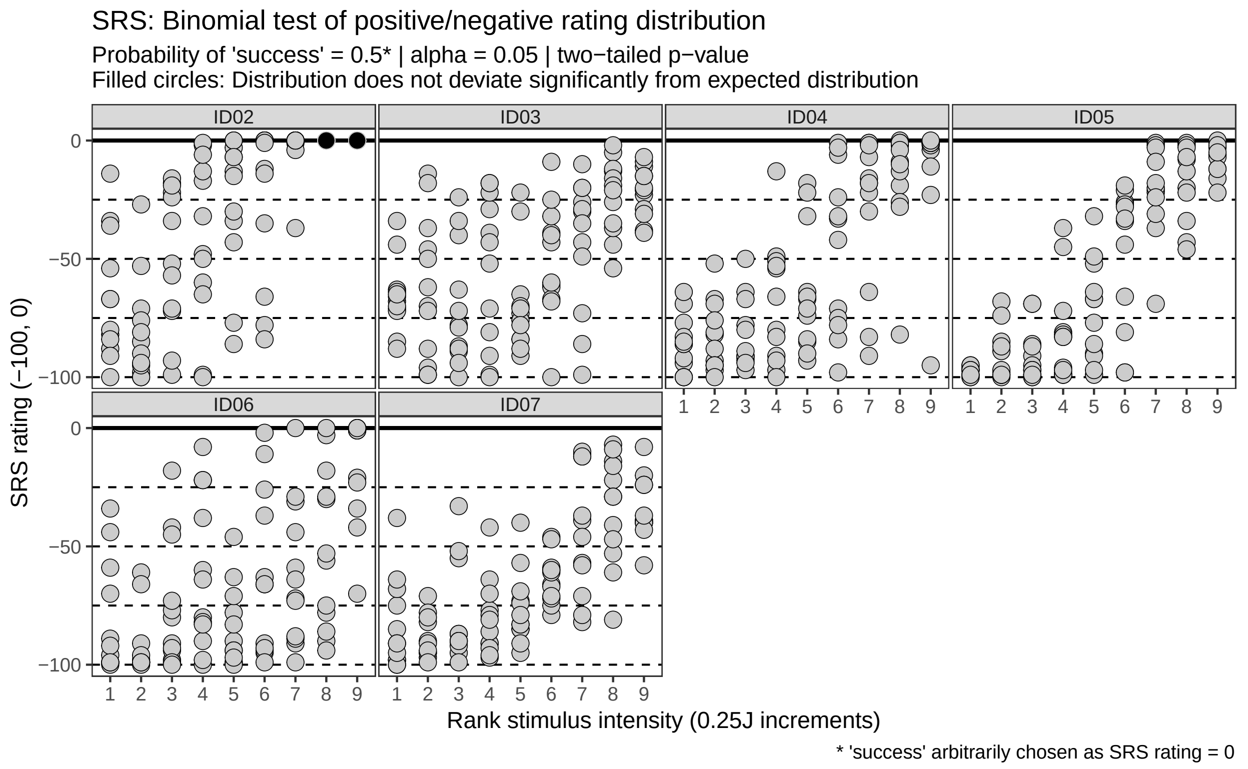

---

### SRS (zero: -15 to 0)

There is no evidential basis to repeat the SRS analysis using a -15 to 0 range as being **'positive'**, but for comparative purposes to the analyses done for the NRS, we performed the analysis of the SRS using the expanded definition of pain threshold.

#### Import and inspect data

Data already imported, inspected, and nested *(data_srs, srs_nest)*.

#### Binomial test

```
# Generate data
srs15_nest <- srs_nest %>%
    # Add probability of success column
    mutate(prob = 0.5) %>%
    # Extract rating data from dataframe
    mutate(data_vec = map(.x = data,
                          ~ .$rating)) %>%
    # Recode rating data as categories according to whether
    # the value is greater than 0 (minimum rating on NRS)
    mutate(data_cat = map(.x = data_vec,
                          ~ ifelse(.x >= -15,
                                   yes = 'positive',
                                   no = 'negative'))) %>%
    # Count the number of positive and negative ratings
    ## positive numbers arbitrarily listed first == 'success'
```

```
    mutate(success_count = map(.x = data_cat,
                               ~ c(length(.x[.x == 'positive']),
                                   length(.x[.x == 'negative'])))) %>%
    # Conduct binomial test (two-sided)
    mutate(binomial_test = map2(.x = success_count,
                                .y = prob,
                                ~ binom.test(x = .x,
                                             p = .y,
                                             alternative = 'greater'))) %>%
    # Extract p-value from binomial_test
    mutate(binomial_p.value = map(.x = binomial_test,
                                  ~ .x$p.value %>%
                                      round(., 3))) %>%
    # Categorise p-value using a p < 0.05 threshold
    ## Significant: distribution deviates significantly
    ## from the theoretical distribution
    ## No correction for multiple comparisons
    ## (too conservative for explorartory analysis)
    mutate(significant_p.value = map(.x = binomial_p.value,
                                     ~ ifelse(.x < 0.05,
                                              yes = 'yes',
                                              no = 'no')))
```

##### Plot

```
srs15_nest %>%
    # Select data columns
    select(PID, intensity, significant_p.value) %>%
    # Unnest data
    unnest() %>%
    # Join with original data
    right_join(data_srs) %>%
    # Reclass intensity as an ordered factor
    mutate(intensity = factor(intensity,
                              ordered = TRUE)) %>%
    # Plot
    ggplot(data = .) +
     aes(x = intensity,
        y = rating,
        fill = significant_p.value,
        colour = significant_p.value) +
    geom_rect(aes(ymin = 0, ymax = -15,
                  xmin = -1, xmax = 10),
              fill = '#666666',
              colour = '#666666') +
    geom_hline(yintercept = 0,
               size = 1) +
    geom_hline(yintercept = -25,
               linetype = 2) +
    geom_hline(yintercept = -50,
               linetype = 2) +
    geom_hline(yintercept = -75,
               linetype = 2) +
    geom_hline(yintercept = -100,
```

```
            linetype = 2) +
geom_point(shape = 21,
           size = 4,
           stroke = 0.3) +
scale_fill_manual(values = c('#CCCCCC', '#000000')) +
scale_colour_manual(values = c('#000000', '#CCCCCC')) +
labs(title = "SRS (0-15): Binomial test of positive/negative rating distribution",
     subtitle = "Probability of 'success' = 0.5* | alpha = 0.05 | two-tailed p-value\nFilled circle
     caption = "* 'success' arbitrarily chosen as SRS rating > -16",
     x = 'Rank stimulus intensity (0.25J increments)',
     y = 'SRS rating (-100, 0)') +
scale_x_discrete(breaks = seq(from = 1,
                              to = 9,
                              by = 1),
                 labels = sprintf('%.0f', seq(from = 1,
                                              to = 9,
                                              by = 1))) +
scale_y_continuous(limits = c(-100, 0),
                   breaks = c(-100, -50, 0),
                   labels = c(-100, -50, 0)) +
facet_wrap(~ PID, ncol = 4) +
theme(legend.position = 'none',
      panel.grid = element_blank(),
      panel.spacing = unit(0.1, 'lines'),
      strip.text = element_text(margin = margin(t = 0.1,
                                                b = 0.1,
                                                r = 1,
                                                l = 1,
                                                'lines')))
```

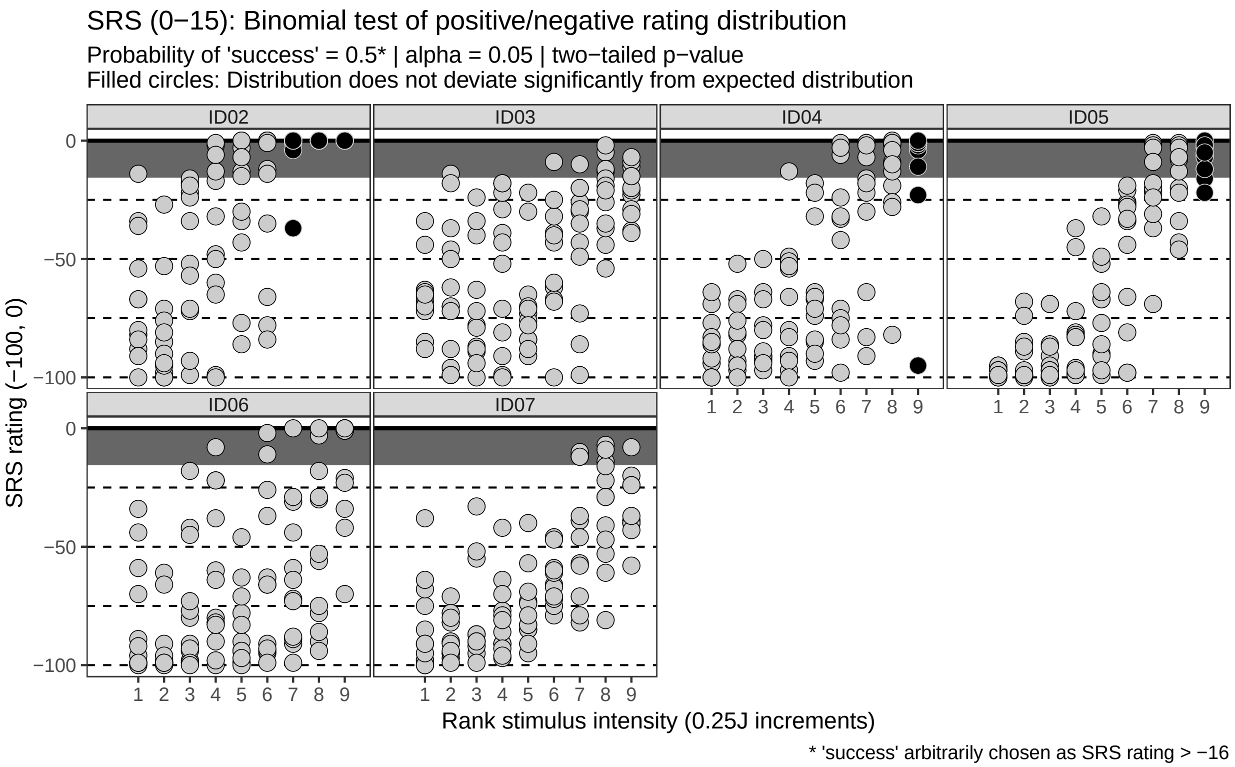

---

### Session information

```r
sessionInfo()
```

```
## R version 3.5.1 (2018-07-02)
## Platform: x86_64-pc-linux-gnu (64-bit)
## Running under: Debian GNU/Linux 9 (stretch)
##
## Matrix products: default
## BLAS: /usr/lib/openblas-base/libblas.so.3
## LAPACK: /usr/lib/libopenblasp-r0.2.19.so
##
## locale:
##  [1] LC_CTYPE=en_US.UTF-8       LC_NUMERIC=C
##  [3] LC_TIME=en_US.UTF-8        LC_COLLATE=en_US.UTF-8
##  [5] LC_MONETARY=en_US.UTF-8    LC_MESSAGES=C
##  [7] LC_PAPER=en_US.UTF-8       LC_NAME=C
##  [9] LC_ADDRESS=C               LC_TELEPHONE=C
## [11] LC_MEASUREMENT=en_US.UTF-8 LC_IDENTIFICATION=C
##
## attached base packages:
## [1] stats     graphics  grDevices utils     datasets  methods   base
##
## other attached packages:
##  [1] bindrcpp_0.2.2  skimr_1.0.3     magrittr_1.5    forcats_0.3.0
##  [5] stringr_1.3.1   dplyr_0.7.8     purrr_0.2.5     readr_1.3.0
##  [9] tidyr_0.8.2     tibble_1.4.2    ggplot2_3.1.0   tidyverse_1.2.1
```

```
##
## loaded via a namespace (and not attached):
##  [1] Rcpp_1.0.0       cellranger_1.1.0 plyr_1.8.4       pillar_1.3.1
##  [5] compiler_3.5.1   bindr_0.1.1      tools_3.5.1      digest_0.6.18
##  [9] lubridate_1.7.4  jsonlite_1.6     evaluate_0.12    nlme_3.1-137
## [13] gtable_0.2.0     lattice_0.20-35  pkgconfig_2.0.2  rlang_0.3.0.1
## [17] cli_1.0.1        rstudioapi_0.8   yaml_2.2.0       haven_2.0.0
## [21] xfun_0.4         withr_2.1.2      xml2_1.2.0       httr_1.4.0
## [25] knitr_1.21       hms_0.4.2        generics_0.0.2   grid_3.5.1
## [29] tidyselect_0.2.5 glue_1.3.0       R6_2.3.0         readxl_1.2.0
## [33] rmarkdown_1.11   modelr_0.1.2     backports_1.1.3  scales_1.0.0
## [37] htmltools_0.3.6  rvest_0.3.2      assertthat_0.2.0 colorspace_1.3-2
## [41] stringi_1.2.4    lazyeval_0.2.1   munsell_0.5.0    broom_0.5.1
## [45] crayon_1.3.4
```
