## Supplementary File 4 for "Rethinking pain threshold as a zone of uncertainty"

Does the difference in stimulus intensity between successive stimuli affect intensity rating

*12 Jan 2019*

**Session information** **21**

---

### Question

We wanted to know whether the difference in intensity between two successive stimuli predicts the rating of the response to the second stimulus.

We hypothesized that ratings of a given stimulus intensity increases as the magnitude of the difference between the stimulus and the preceding stimulus increases.

We assessed this relationship graphically by plotting:

1. Plotting the ratings at each stimulus intensity, and for each participant, corresponding with the maximum and minimum absolute difference in stimulus intensity between a given stimulus intenisty and the preceding stimulus.
2. Plotting all ratings at each stimulus intensity, and for each participant, and colour coding the data points according to the difference in stimulus intensity to the preceding stimulus for each rating.

---

### SPARS A

#### Import and inspect data

```
# Import
data_sparsA <- read_rds('data-cleaned/SPARS_A.rds')

# Inspect
glimpse(data_sparsA)
```

```
## Observations: 1,927
## Variables: 6
## $ PID          <chr> "ID01", "ID01", "ID01", "ID01", "ID01", "ID01", "...
## $ block        <chr> "A", "A", "A", "A", "A", "A", "A", "A", "A", "A",...
## $ block_order  <dbl> 4, 4, 4, 4, 4, 4, 4, 4, 4, 4, 4, 4, 4, 4, 4, 4, 4...
## $ trial_number <dbl> 79, 80, 81, 82, 83, 84, 85, 86, 87, 88, 89, 90, 9...
## $ intensity    <dbl> 3.00, 2.25, 4.00, 3.25, 2.75, 2.25, 2.75, 4.00, 2...
## $ rating       <dbl> -40, -25, 10, 2, -10, -25, -20, 10, -25, -50, -25...
```

```
data_sparsA %>%
    select(intensity, rating) %>%
    skim()
```

```
## Skim summary statistics
##  n obs: 1927
##  n variables: 2
##
## -- Variable type:numeric ------------------------------------------------------------
##   variable missing complete    n  mean    sd  p0    p25 p50  p75 p100
##  intensity       0     1927 1927  2.47  0.93   1   1.75 2.5  3.25    4
##     rating       0     1927 1927 -4.45 22.31 -50 -20    2   10      45
##      hist
##
##
```

#### Process the data

```
# Select columns
data_sparsA %<>%
    select(PID, block, trial_number, intensity, rating)

# Nest data by PID
sparsA_nest <- data_sparsA %>%
    group_by(PID) %>%
    nest()

# Group nested data by block
```

```r
sparsA_nest %<>%
   mutate(data = map(.x = data,
                    ~ .x %>%
                         group_by(block)))

# Sort each block by trial number
sparsA_nest %<>%
    mutate(data = map(.x = data,
                      ~ .x %>%
                           arrange(trial_number)))

# Calculate the absolute value of the lag one stimulus intensity difference
sparsA_nest %<>%
    # Extract intensity from 'data'
    mutate(data = map(.x = data,
                      ~ .x %>%
                          mutate(delta_intensity = abs(intensity -
                                                           lag(intensity))) %>%
                          # Remove stimulus 1 of each block (<NA>)
                          filter(!is.na(delta_intensity)))) %>%
    # Unnest dataframe
    unnest()

# Add max/min plot colour coding
sparsA_nest %<>%
    group_by(PID, intensity) %>%
    mutate(colour = case_when(
        delta_intensity == max(delta_intensity) ~ 'max',
        delta_intensity == min(delta_intensity) ~ 'min',
        delta_intensity > min(delta_intensity) &
            delta_intensity < max(delta_intensity) ~ 'other'
    )) %>%
    ungroup() %>%
    arrange(PID, intensity, delta_intensity)
```

#### Plots

##### Maximum and minimum inter-stimulus intensity change only

```r
sparsA_nest %>%
    filter(colour != 'other') %>%
    ggplot(data = .) +
    aes(x = intensity,
        y = rating,
        fill = colour) +
    geom_hline(yintercept = 0,
               size = 1) +
    geom_hline(yintercept = 25,
               linetype = 2) +
    geom_hline(yintercept = 50,
               linetype = 2) +
    geom_hline(yintercept = -25,
               linetype = 2) +
    geom_hline(yintercept = -50,
               linetype = 2) +
```

```
geom_point(shape = 21,
           size = 4,
           stroke = 0.3) +
labs(title = "SPARS A: Scatterplot of intensity ratings at each stimulus intensity\nfor the max and
     caption = "* The absolute value of the difference in intensity between successive stimuli was
     x = 'Stimulus intensity (J)',
     y = 'SPARS rating (-50, 50)') +
scale_x_continuous(breaks = seq(from = 1,
                                to = 4,
                                by = 0.5)) +
scale_y_continuous(limits = c(-50, 50),
                   breaks = c(-50, 0, 50),
                   labels = c(-50, 0, 50)) +
scale_fill_viridis_d(name = 'Inter-stimulus change (J): ',
                     option = 'C') +
facet_wrap(~ PID, ncol = 4) +
theme(legend.position = 'top',
      legend.margin = margin(t = -0.2,
                             l = 0,
                             b = -0.4,
                             r = 0,
                             unit = 'lines'),
      panel.grid = element_blank(),
      panel.spacing = unit(0.1, 'lines'),
      strip.text = element_text(margin = margin(t = 0.1,
                                                b = 0.1,
                                                r = 1,
                                                l = 1,
                                                'lines')),
      axis.text.x = element_text(angle = -90,
                                 vjust = 0.5))
```

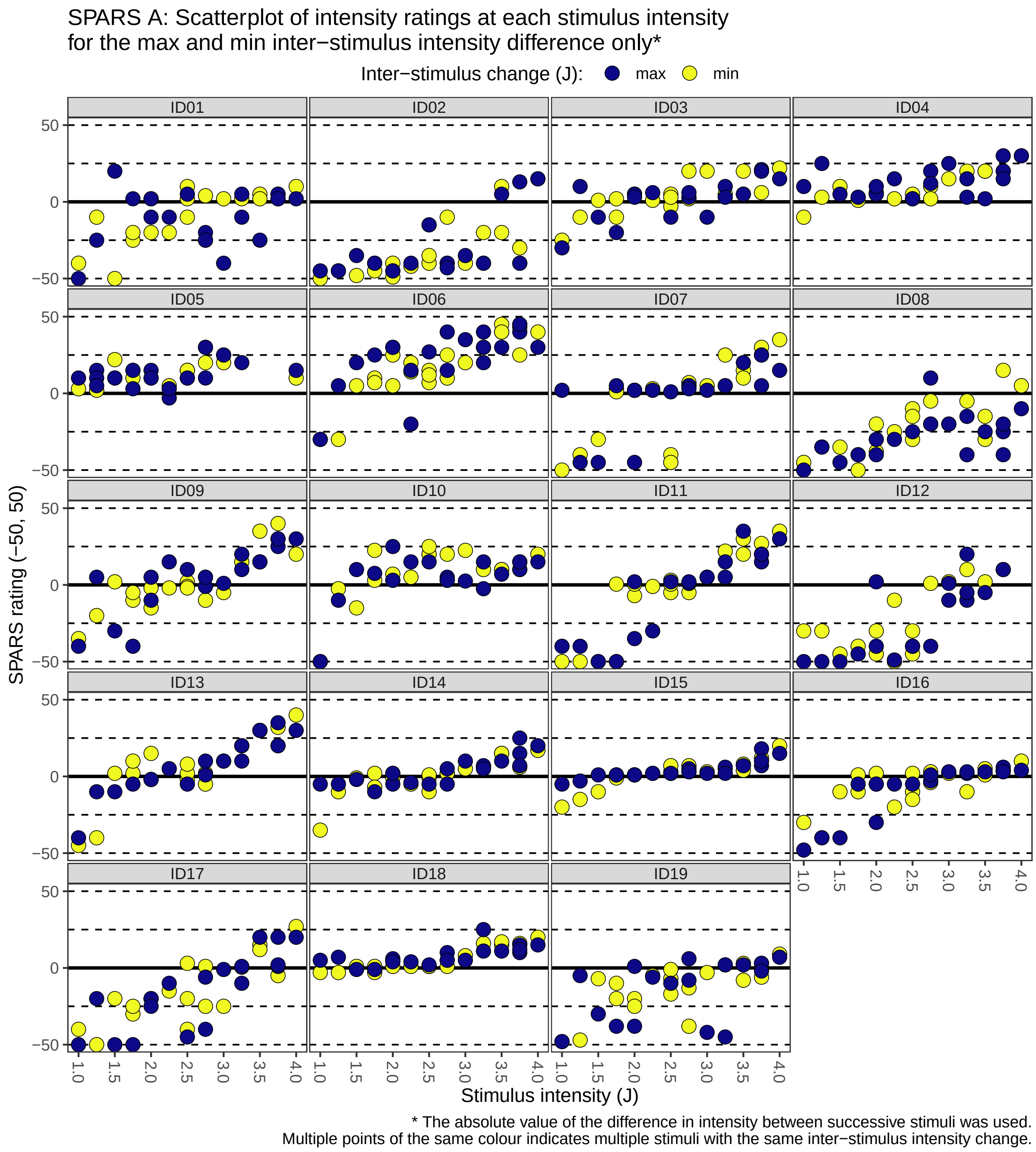

##### All inter-stimulus intensity changes

```
sparsA_nest %>%
    ggplot(data = .) +
    aes(x = intensity,
        y = rating,
        fill = delta_intensity) +
    geom_hline(yintercept = 0,
               size = 1) +
    geom_hline(yintercept = 25,
```

```
           linetype = 2) +
geom_hline(yintercept = 50,
           linetype = 2) +
geom_hline(yintercept = -25,
           linetype = 2) +
geom_hline(yintercept = -50,
           linetype = 2) +
geom_point(shape = 21,
           size = 4,
           stroke = 0.3) +
labs(title = "SPARS A: Scatterplot of intensity ratings at each stimulus intensity\nfor all inter-st
     caption = "* The absolute value of the difference in intensity between successive stimuli was u
     x = 'Stimulus intensity (J)',
     y = 'SPARS rating (-50, 50)') +
scale_x_continuous(breaks = seq(from = 1,
                                to = 4,
                                by = 0.5)) +
scale_y_continuous(limits = c(-50, 50),
                   breaks = c(-50, 0, 50),
                   labels = c(-50, 0, 50)) +
scale_fill_viridis_c(name = 'Inter-stimulus change (J): ',
                     option = 'C') +
facet_wrap(~ PID, ncol = 4) +
theme(legend.position = 'top',
      legend.margin = margin(t = -0.2,
                             l = 0,
                             b = -0.4,
                             r = 0,
                             unit = 'lines'),
      panel.grid = element_blank(),
      panel.spacing = unit(0.1, 'lines'),
      strip.text = element_text(margin = margin(t = 0.1,
                                                b = 0.1,
                                                r = 1,
                                                l = 1,
                                                'lines')),
      axis.text.x = element_text(angle = -90,
                                 vjust = 0.5))
```

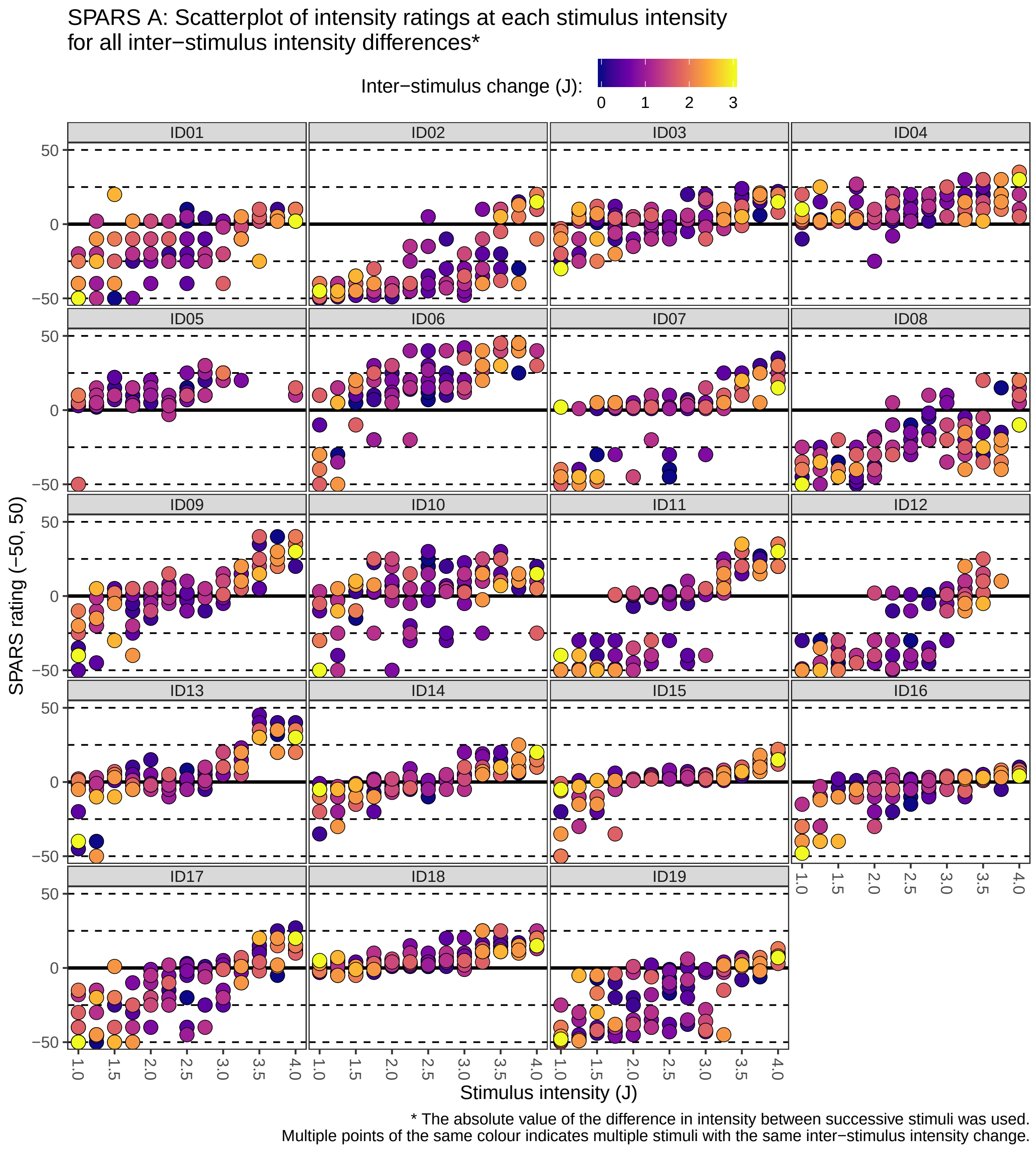

---

### SPARS B

#### Import and inspect data

```
# Import
data_sparsB <- read_rds('data-cleaned/SPARS_B.rds') %>%
    # Extract trials rated using the SPARS
```

```r
    filter(scale == 'SPARS') %>%
    # Remove <NA>
    filter(!is.na(rating))

# Rank stimulus intensity
data_sparsB %<>%
    group_by(PID, scale) %>%
    arrange(intensity) %>%
    mutate(intensity_rank = dense_rank(intensity)) %>%
    select(-intensity) %>%
    rename(intensity = intensity_rank) %>%
    ungroup()

# Inspect
glimpse(data_sparsB)
```

```
## Observations: 752
## Variables: 6
## $ PID          <chr> "ID06", "ID06", "ID06", "ID06", "ID06", "ID06", "...
## $ block_number <int> 1, 1, 1, 2, 2, 2, 3, 3, 3, 4, 4, 4, 1, 1, 1, 2, 2...
## $ trial_number <dbl> 4, 6, 27, 9, 13, 20, 20, 24, 27, 4, 18, 22, 5, 16...
## $ scale        <chr> "SPARS", "SPARS", "SPARS", "SPARS", "SPARS", "SPA...
## $ rating       <dbl> -49, 2, -6, 3, -20, -2, -31, 2, -5, -8, -23, 14, ...
## $ intensity    <int> 1, 1, 1, 1, 1, 1, 1, 1, 1, 1, 1, 1, 2, 2, 2, 2, 2...
```

```r
data_sparsB %>%
    select(intensity, rating) %>%
    skim()
```

```
## Skim summary statistics
##  n obs: 752
##  n variables: 2
##
## -- Variable type:integer -----------------------------------------------------------
##   variable missing complete   n mean   sd p0 p25 p50 p75 p100     hist
##  intensity       0      752 752    5 2.58  1   3   5   7    9
##
## -- Variable type:numeric -----------------------------------------------------------
##  variable missing complete   n  mean    sd  p0 p25 p50 p75 p100     hist
##    rating       0      752 752 -8.83 23.46 -50 -26  -4   5   50
```

#### Process the data

```r
# Select columns
data_sparsB %<>%
    select(PID, block_number, trial_number, intensity, rating)

# Nest data by PID
sparsB_nest <- data_sparsB %>%
    group_by(PID) %>%
    nest()

# Group nested data by block
sparsB_nest %<>%
   mutate(data = map(.x = data,
                     ~ .x %>%
```

```r
                        group_by(block_number)))

# Sort each block by trial number
sparsB_nest %<>%
    mutate(data = map(.x = data,
                      ~ .x %>%
                        arrange(trial_number)))

# Calculate the absolute value of the lag one stimulus intensity difference
sparsB_nest %<>%
    # Extract intensity from 'data'
    mutate(data = map(.x = data,
                      ~ .x %>%
                        mutate(delta_intensity = abs(intensity -
                                                     lag(intensity))) %>%
                        # Remove stimulus 1 of each block (<NA>)
                        filter(!is.na(delta_intensity)))) %>%
    # Unnest dataframe
    unnest()

# Add max/min plot colour coding
sparsB_nest %<>%
    group_by(PID, intensity) %>%
    mutate(colour = case_when(
        delta_intensity == max(delta_intensity) ~ 'max',
        delta_intensity == min(delta_intensity) ~ 'min',
        delta_intensity > min(delta_intensity) &
            delta_intensity < max(delta_intensity) ~ 'other'
    )) %>%
    ungroup() %>%
    arrange(PID, intensity, delta_intensity)
```

#### Plots

##### Maximum and minimum inter-stimulus intensity change only

```r
sparsB_nest %>%
    filter(colour != 'other') %>%
    ggplot(data = .) +
    aes(x = intensity,
        y = rating,
        fill = colour) +
    geom_hline(yintercept = 0,
               size = 1) +
    geom_hline(yintercept = 25,
               linetype = 2) +
    geom_hline(yintercept = 50,
               linetype = 2) +
    geom_hline(yintercept = -25,
               linetype = 2) +
    geom_hline(yintercept = -50,
               linetype = 2) +
    geom_point(shape = 21,
               size = 4,
               stroke = 0.3) +
```

```
labs(title = "SPARS A: Scatterplot of intensity ratings at each stimulus intensity\nfor the max and
     caption = "* The absolute value of the difference in intensity between successive stimuli was u
     x = 'Rank stimulus intensity (0.25J increments)',
     y = 'SPARS rating (-50 to 50)') +
scale_x_continuous(breaks = 1:9) +
scale_y_continuous(limits = c(-50, 50),
                   breaks = c(-50, 0, 50),
                   labels = c(-50, 0, 50)) +
scale_fill_viridis_d(name = 'Inter-stimulus change (J): ',
                     option = 'C') +
facet_wrap(~ PID, ncol = 4) +
theme(legend.position = 'top',
      legend.margin = margin(t = -0.2,
                             l = 0,
                             b = -0.4,
                             r = 0,
                             unit = 'lines'),
      panel.grid = element_blank(),
      panel.spacing = unit(0.1, 'lines'),
      strip.text = element_text(margin = margin(t = 0.1,
                                                b = 0.1,
                                                r = 1,
                                                l = 1,
                                                'lines')))
```

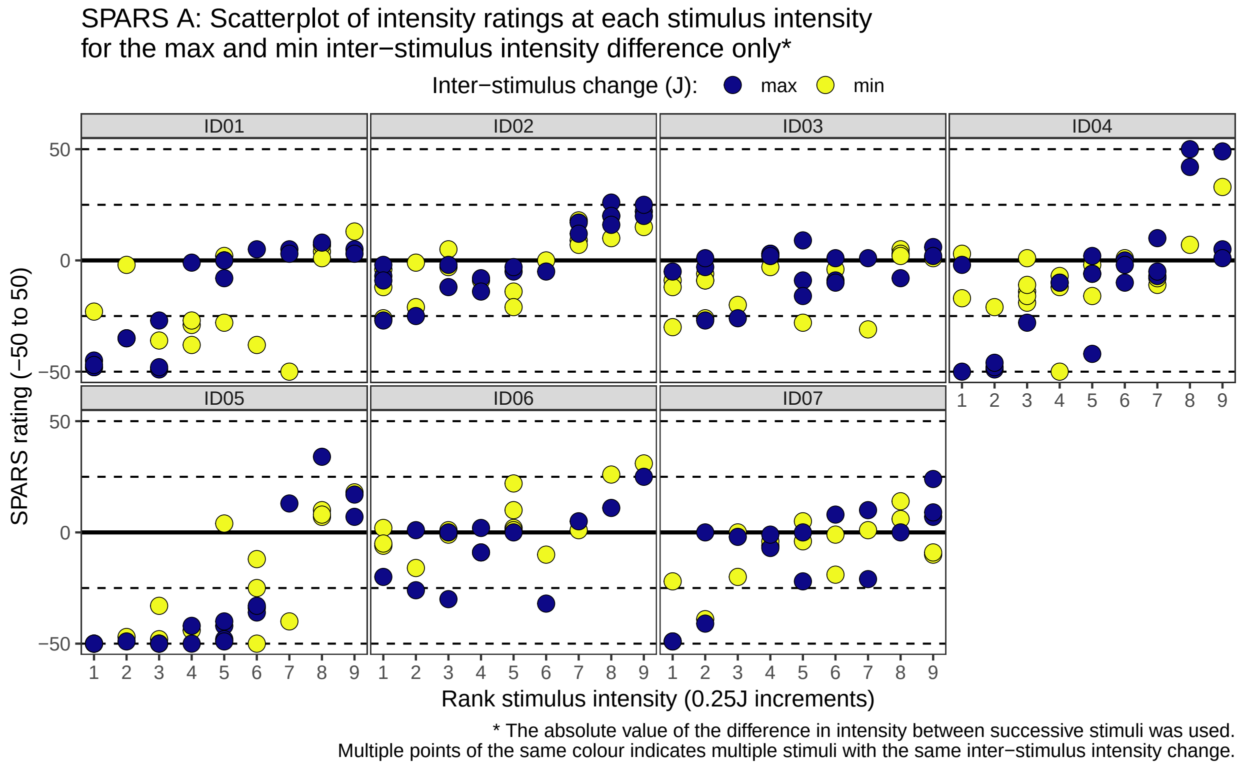

##### All inter-stimulus intensity changes

```
sparsB_nest %>%
    ggplot(data = .) +
    aes(x = intensity,
        y = rating,
        fill = delta_intensity) +
    geom_hline(yintercept = 0,
               size = 1) +
    geom_hline(yintercept = 25,
               linetype = 2) +
    geom_hline(yintercept = 50,
               linetype = 2) +
    geom_hline(yintercept = -25,
               linetype = 2) +
    geom_hline(yintercept = -50,
               linetype = 2) +
    geom_point(shape = 21,
               size = 4,
               stroke = 0.3) +
    labs(title = "SPARS B: Scatterplot of intensity ratings at each (rank) stimulus intensity\nfor all i
         caption = "* The absolute value of the difference in intensity between successive stimuli was u
         x = 'Rank stimulus intensity (0.25J increments)',
         y = 'SPARS rating (-50, 50)') +
    scale_x_continuous(breaks = 1:9) +
    scale_y_continuous(limits = c(-50, 50),
                       breaks = c(-50, 0, 50),
                       labels = c(-50, 0, 50)) +
    scale_fill_viridis_c(name = 'Inter-stimulus change (J): ',
                         option = 'C') +
    facet_wrap(~ PID, ncol = 4) +
    theme(legend.position = 'top',
          legend.margin = margin(t = -0.2,
                                 l = 0,
                                 b = -0.4,
                                 r = 0,
                                 unit = 'lines'),
          panel.grid = element_blank(),
          panel.spacing = unit(0.1, 'lines'),
          strip.text = element_text(margin = margin(t = 0.1,
                                                    b = 0.1,
                                                    r = 1,
                                                    l = 1,
                                                    'lines')))
```

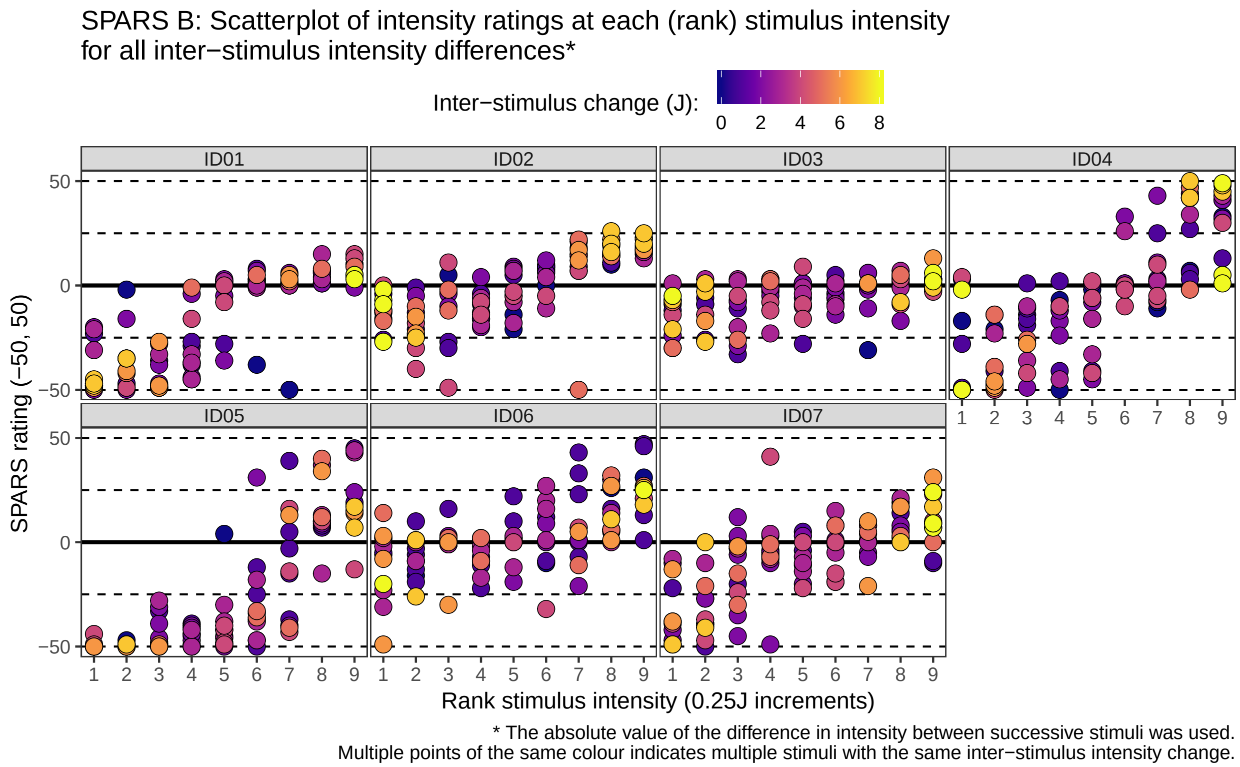

---

### NRS

#### Import and inspect data

```
# Import
data_nrs <- read_rds('data-cleaned/SPARS_B.rds') %>%
    # Extract trials rated using the SPARS
    filter(scale == 'NRS') %>%
    # Remove <NA>
    filter(!is.na(rating))

# Rank stimulus intensity
data_nrs %<>%
    group_by(PID, scale) %>%
    arrange(intensity) %>%
    mutate(intensity_rank = dense_rank(intensity)) %>%
    select(-intensity) %>%
    rename(intensity = intensity_rank) %>%
    ungroup()

# Inspect
glimpse(data_nrs)
```

```
## Observations: 753
## Variables: 6
## $ PID          <chr> "ID06", "ID06", "ID06", "ID06", "ID06", "ID06", "...
## $ block_number <int> 9, 9, 9, 10, 10, 10, 11, 11, 11, 12, 12, 12, 9, 9...
```

```
## $ trial_number <dbl> 7, 9, 26, 4, 9, 27, 2, 4, 12, 4, 7, 10, 5, 6, 27,...
## $ scale        <chr> "NRS", "NRS", "NRS", "NRS", "NRS", "NRS", "NRS", ...
## $ rating       <dbl> 5, 2, 0, 0, 0, 0, 0, 1, 0, 0, 0, 48, 1, 0, 53, 0,...
## $ intensity    <int> 1, 1, 1, 1, 1, 1, 1, 1, 1, 1, 1, 1, 2, 2, 2, 2, 2...
```

```
data_nrs %>%
    select(intensity, rating) %>%
    skim()
```

```
## Skim summary statistics
##  n obs: 753
##  n variables: 2
##
## -- Variable type:integer -----------------------------------------------------------
##   variable missing complete   n mean   sd p0 p25 p50 p75 p100     hist
##  intensity       0      753 753    5 2.59  1   3   5   7    9
##
## -- Variable type:numeric -----------------------------------------------------------
##  variable missing complete   n  mean    sd p0 p25 p50 p75 p100     hist
##    rating       0      753 753 19.63 26.82  0   1   5  28   98
```

#### Process the data

```
# Select columns
data_nrs <- data_nrs %>%
    select(PID, block_number, trial_number, intensity, rating)

# Nest data by PID
nrs_nest <- data_nrs %>%
    group_by(PID) %>%
    nest()

# Group nested data by block
nrs_nest %<>%
   mutate(data = map(.x = data,
                    ~ .x %>%
                        group_by(block_number)))

# Sort each block by trial number
nrs_nest %<>%
    mutate(data = map(.x = data,
                      ~ .x %>%
                          arrange(trial_number)))

# Calculate the absolute value of the lag one stimulus intensity difference
nrs_nest %<>%
    # Extract intensity from 'data'
    mutate(data = map(.x = data,
                      ~ .x %>%
                          mutate(delta_intensity = abs(intensity -
                                                           lag(intensity))) %>%
                          # Remove stimulus 1 of each block (<NA>)
                          filter(!is.na(delta_intensity)))) %>%
    # Unnest dataframe
    unnest()
```

```
# Add max/min plot colour coding
nrs_nest %<>%
    group_by(PID, intensity) %>%
    mutate(colour = case_when(
        delta_intensity == max(delta_intensity) ~ 'max',
        delta_intensity == min(delta_intensity) ~ 'min',
        delta_intensity > min(delta_intensity) &
            delta_intensity < max(delta_intensity) ~ 'other'
    )) %>%
    ungroup() %>%
    arrange(PID, intensity, delta_intensity)
```

#### Plots

##### Maximum and minimum inter-stimulus intensity change only

```
nrs_nest %>%
    filter(colour != 'other') %>%
    ggplot(data = .) +
    aes(x = intensity,
        y = rating,
        fill = colour) +
    geom_hline(yintercept = 0,
               size = 1) +
    geom_hline(yintercept = 25,
               linetype = 2) +
    geom_hline(yintercept = 50,
               linetype = 2) +
    geom_hline(yintercept = 75,
               linetype = 2) +
    geom_hline(yintercept = 100,
               linetype = 2) +
    geom_point(shape = 21,
               size = 4,
               stroke = 0.3) +
    labs(title = "SPARS A: Scatterplot of intensity ratings at each stimulus intensity\nfor the max and
         caption = "* The absolute value of the difference in intensity between successive stimuli was
         x = 'Rank stimulus intensity (0.25J increments)',
         y = 'NRS rating (0 to 100)') +
    scale_x_continuous(breaks = 1:9) +
    scale_y_continuous(limits = c(0, 100),
                       breaks = c(0, 50, 100),
                       labels = c(0, 50, 100)) +
    scale_fill_viridis_d(name = 'Inter-stimulus change (J): ',
                         option = 'C') +
    facet_wrap(~ PID, ncol = 4) +
    theme(legend.position = 'top',
          legend.margin = margin(t = -0.2,
                                 l = 0,
                                 b = -0.4,
                                 r = 0,
                                 unit = 'lines'),
          panel.grid = element_blank(),
          panel.spacing = unit(0.1, 'lines'),
```

```
strip.text = element_text(margin = margin(t = 0.1,
                                          b = 0.1,
                                          r = 1,
                                          l = 1,
                                          'lines')))
```

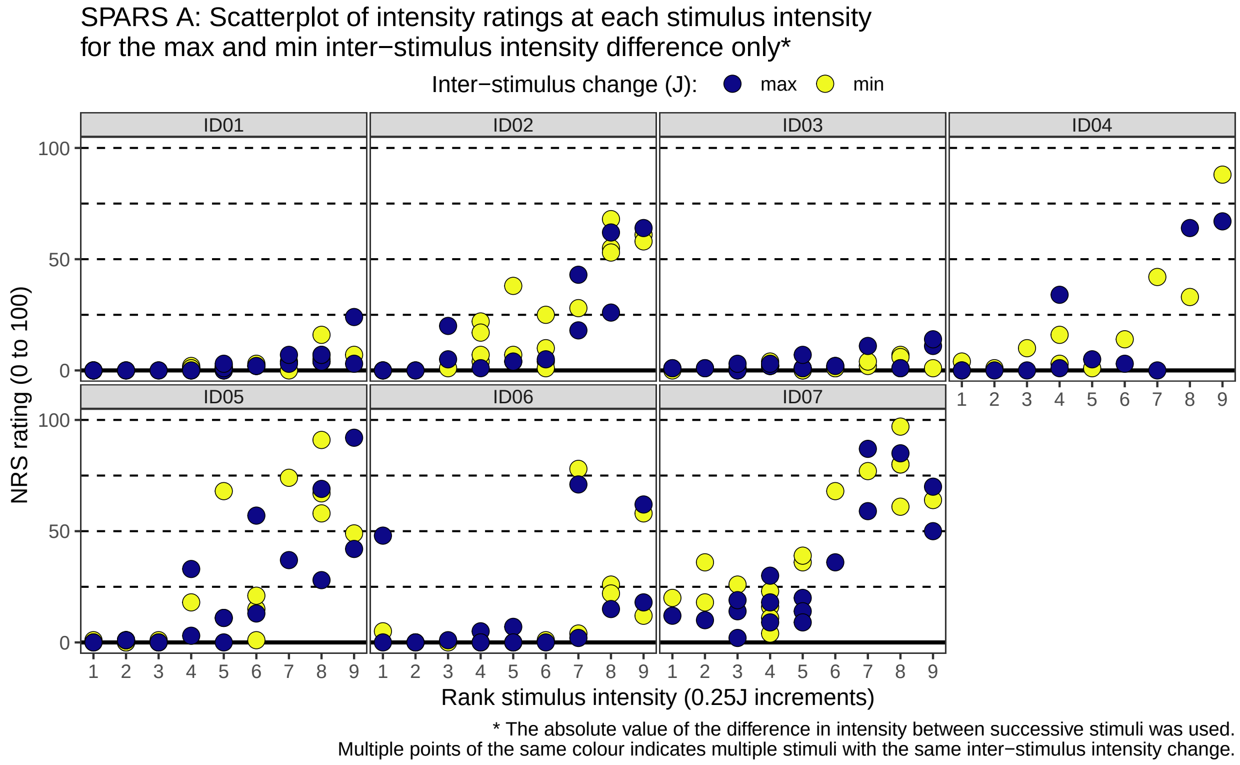

##### All inter-stimulus intensity changes

```
nrs_nest %>%
    ggplot(data = .) +
    aes(x = intensity,
        y = rating,
        fill = delta_intensity) +
    geom_hline(yintercept = 0,
               size = 1) +
    geom_hline(yintercept = 25,
               linetype = 2) +
    geom_hline(yintercept = 50,
               linetype = 2) +
    geom_hline(yintercept = 75,
               linetype = 2) +
    geom_hline(yintercept = 100,
               linetype = 2) +
    geom_point(shape = 21,
               size = 4,
               stroke = 0.3) +
    labs(title = "SPARS A: Scatterplot of intensity ratings at each stimulus intensity\for all inter-sti
         caption = "* The absolute value of the difference in intensity between successive stimuli was u
```

```
     x = 'Rank stimulus intensity (0.25J increments)',
     y = 'NRS rating (0 to 100)') +
scale_x_continuous(breaks = 1:9) +
scale_y_continuous(limits = c(0, 100),
                   breaks = c(0, 50, 100),
                   labels = c(0, 50, 100)) +
scale_fill_viridis_c(name = 'Inter-stimulus change (J): ',
                     option = 'C') +
facet_wrap(~ PID, ncol = 4) +
theme(legend.position = 'top',
      legend.margin = margin(t = -0.2,
                             l = 0,
                             b = -0.4,
                             r = 0,
                             unit = 'lines'),
      panel.grid = element_blank(),
      panel.spacing = unit(0.1, 'lines'),
      strip.text = element_text(margin = margin(t = 0.1,
                                                b = 0.1,
                                                r = 1,
                                                l = 1,
                                                'lines')))
```

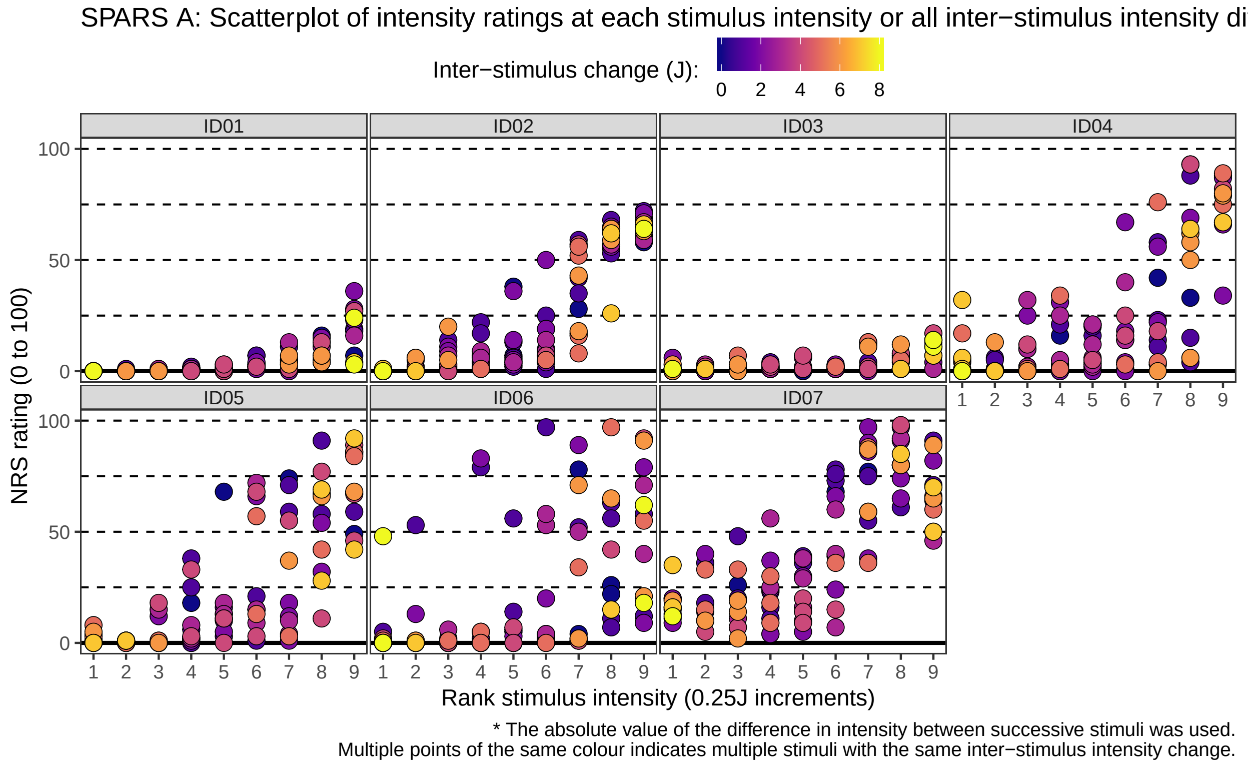

---

### SRS

#### Import and inspect data

```
# Import
data_srs <- read_rds('data-cleaned/SPARS_B.rds') %>%
    # Extract trials rated using the SPARS
    filter(scale == 'SRS') %>%
    # Remove <NA>
    filter(!is.na(rating))

# Rank stimulus intensity
data_srs %<>%
    group_by(PID, scale) %>%
    arrange(intensity) %>%
    mutate(intensity_rank = dense_rank(intensity)) %>%
    select(-intensity) %>%
    rename(intensity = intensity_rank) %>%
    ungroup()

# Inspect
glimpse(data_srs)
```

```
## Observations: 644
## Variables: 6
## $ PID          <chr> "ID06", "ID06", "ID06", "ID06", "ID06", "ID06", "...
## $ block_number <int> 5, 5, 5, 6, 6, 6, 7, 7, 7, 8, 8, 8, 5, 5, 5, 6, 6...
## $ trial_number <dbl> 2, 16, 26, 13, 19, 21, 1, 17, 27, 2, 4, 5, 6, 24,...
## $ scale        <chr> "SRS", "SRS", "SRS", "SRS", "SRS", "SRS", "SRS", ...
## $ rating       <dbl> -34, -99, -89, -99, -100, -99, -59, -96, -70, -92...
## $ intensity    <int> 1, 1, 1, 1, 1, 1, 1, 1, 1, 1, 1, 1, 2, 2, 2, 2, 2...
```

```
data_srs %>%
    select(intensity, rating) %>%
    skim()
```

```
## Skim summary statistics
##  n obs: 644
##  n variables: 2
##
## -- Variable type:integer ----------------------------------------------------------
##   variable missing complete   n mean   sd p0 p25 p50 p75 p100     hist
##  intensity       0      644 644    5 2.58  1   3   5   7    9
##
## -- Variable type:numeric ----------------------------------------------------------
##  variable missing complete   n   mean    sd   p0 p25 p50 p75 p100     hist
##    rating       0      644 644 -54.46 35.19 -100 -88 -63 -21    0
```

#### Process the data

```
# Select columns
data_srs <- data_srs %>%
    select(PID, block_number, trial_number, intensity, rating)

# Nest data by PID
srs_nest <- data_srs %>%
```

```
    group_by(PID) %>%
    nest()

# Group nested data by block
srs_nest %<>%
   mutate(data = map(.x = data,
                    ~ .x %>%
                        group_by(block_number)))

# Sort each block by trial number
srs_nest %<>%
    mutate(data = map(.x = data,
                      ~ .x %>%
                        arrange(trial_number)))

# Calculate the absolute value of the lag one stimulus intensity difference
srs_nest %<>%
    # Extract intensity from 'data'
    mutate(data = map(.x = data,
                      ~ .x %>%
                        mutate(delta_intensity = abs(intensity -
                                                     lag(intensity))) %>%
                        # Remove stimulus 1 of each block (<NA>)
                        filter(!is.na(delta_intensity)))) %>%
    # Unnest dataframe
    unnest()

# Add max/min plot colour coding
srs_nest %<>%
    group_by(PID, intensity) %>%
    mutate(colour = case_when(
        delta_intensity == max(delta_intensity) ~ 'max',
        delta_intensity == min(delta_intensity) ~ 'min',
        delta_intensity > min(delta_intensity) &
            delta_intensity < max(delta_intensity) ~ 'other'
    )) %>%
    ungroup() %>%
    arrange(PID, intensity, delta_intensity)
```

#### Plots

##### Maximum and minimum inter-stimulus intensity change only

```
srs_nest %>%
    filter(colour != 'other') %>%
    ggplot(data = .) +
    aes(x = intensity,
        y = rating,
        fill = colour) +
    geom_hline(yintercept = 0,
               size = 1) +
    geom_hline(yintercept = -25,
               linetype = 2) +
    geom_hline(yintercept = -50,
               linetype = 2) +
```

```
geom_hline(yintercept = -75,
           linetype = 2) +
geom_hline(yintercept = -100,
           linetype = 2) +
geom_point(shape = 21,
           size = 4,
           stroke = 0.3) +
labs(title = "SRS: Scatterplot of intensity ratings at each (rank) stimulus intensity\nfor the max a
     caption = "* The absolute value of the difference in intensity between successive stimuli was 
     x = 'Rank stimulus intensity (0.25J increments)',
     y = 'SRS rating (-100 to 0)') +
scale_x_continuous(breaks = 1:9) +
scale_y_continuous(limits = c(-100, 0),
                   breaks = c(-100, -50, 0),
                   labels = c(-100, -50, 0)) +
scale_fill_viridis_d(name = 'Inter-stimulus change (J): ',
                     option = 'C') +
facet_wrap(~ PID, ncol = 4) +
theme(legend.position = 'top',
      legend.margin = margin(t = -0.2,
                             l = 0,
                             b = -0.4,
                             r = 0,
                             unit = 'lines'),
      panel.grid = element_blank(),
      panel.spacing = unit(0.1, 'lines'),
      strip.text = element_text(margin = margin(t = 0.1,
                                                b = 0.1,
                                                r = 1,
                                                l = 1,
                                                'lines')))
```

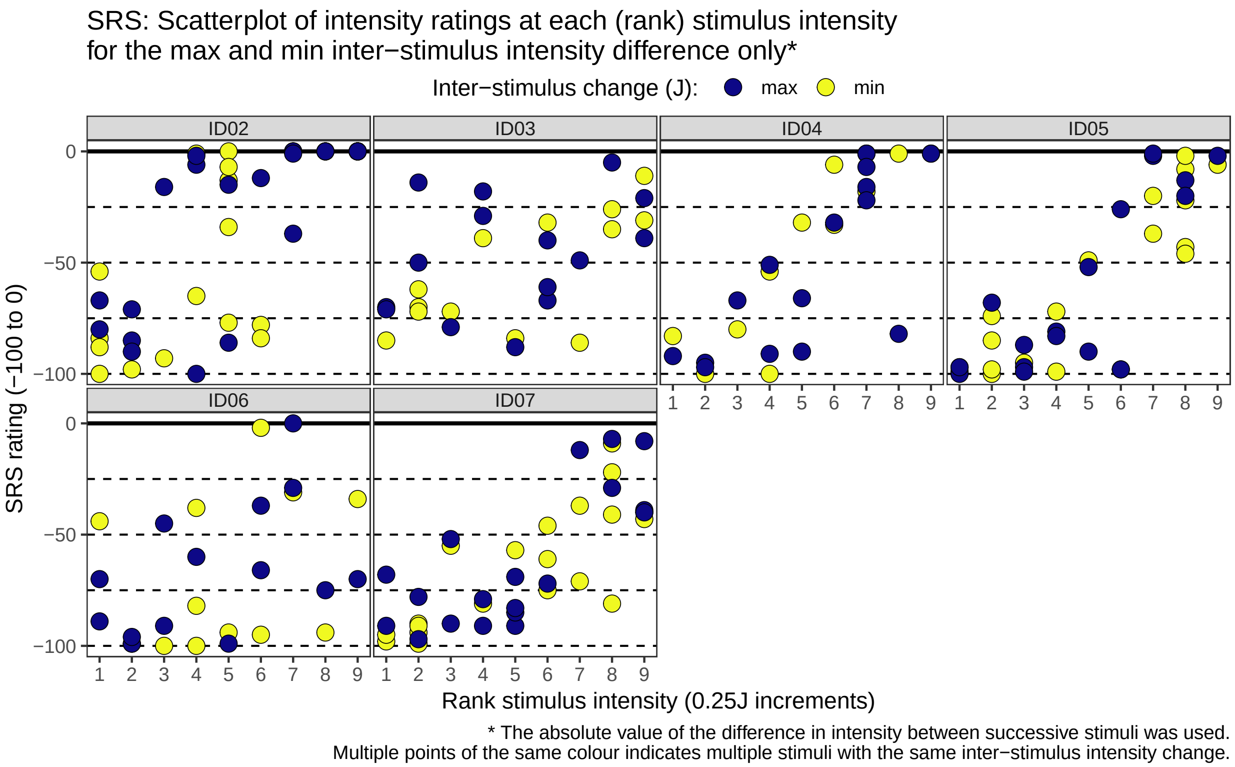

##### All inter-stimulus intensity changes

```
srs_nest %>%
    ggplot(data = .) +
    aes(x = intensity,
        y = rating,
        fill = delta_intensity) +
    geom_hline(yintercept = 0,
               size = 1) +
    geom_hline(yintercept = -25,
               linetype = 2) +
    geom_hline(yintercept = -50,
               linetype = 2) +
    geom_hline(yintercept = -75,
               linetype = 2) +
    geom_hline(yintercept = -100,
               linetype = 2) +
    geom_point(shape = 21,
               size = 4,
               stroke = 0.3) +
    labs(title = "SRS: Scatterplot of intensity ratings at each (rank) stimulus intensity\nfor all inte
         caption = "* The absolute value of the difference in intensity between successive stimuli was u
         x = 'Rank stimulus intensity (0.25J increments)',
         y = 'SRS rating (-100 to 0)') +
    scale_x_continuous(breaks = 1:9) +
    scale_y_continuous(limits = c(-100, 0),
                       breaks = c(-100, -50, 0),
                       labels = c(-100, -50, 0)) +
    scale_fill_viridis_c(name = 'Inter-stimulus change (J): ',
```

```
                           option = 'C') +
  facet_wrap(~ PID, ncol = 4) +
  theme(legend.position = 'top',
        legend.margin = margin(t = -0.2,
                                l = 0,
                                b = -0.4,
                                r = 0,
                                unit = 'lines'),
        panel.grid = element_blank(),
        panel.spacing = unit(0.1, 'lines'),
        strip.text = element_text(margin = margin(t = 0.1,
                                                   b = 0.1,
                                                   r = 1,
                                                   l = 1,
                                                   'lines')))
```

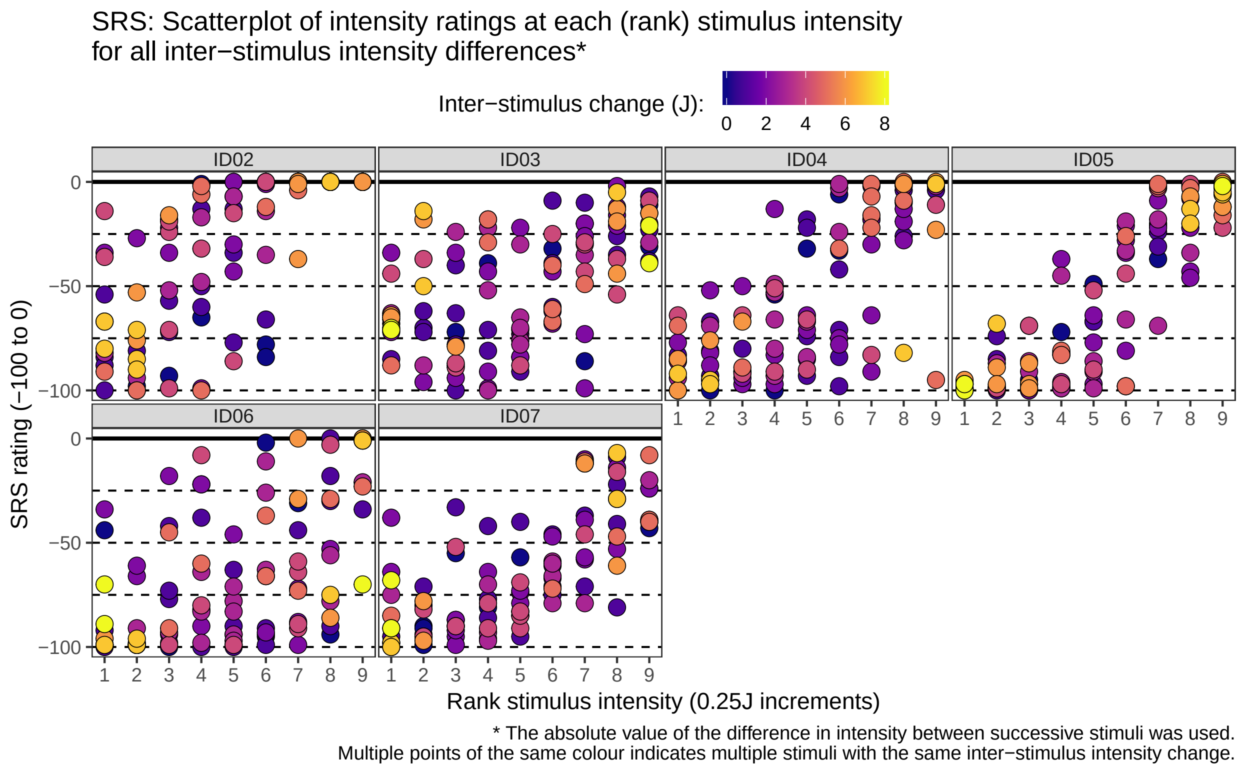

---

### Session information

```
sessionInfo()
```

```
## R version 3.5.1 (2018-07-02)
## Platform: x86_64-pc-linux-gnu (64-bit)
## Running under: Debian GNU/Linux 9 (stretch)
##
## Matrix products: default
## BLAS: /usr/lib/openblas-base/libblas.so.3
```

```
## LAPACK: /usr/lib/libopenblasp-r0.2.19.so
##
## locale:
##  [1] LC_CTYPE=en_US.UTF-8       LC_NUMERIC=C
##  [3] LC_TIME=en_US.UTF-8        LC_COLLATE=en_US.UTF-8
##  [5] LC_MONETARY=en_US.UTF-8    LC_MESSAGES=C
##  [7] LC_PAPER=en_US.UTF-8       LC_NAME=C
##  [9] LC_ADDRESS=C               LC_TELEPHONE=C
## [11] LC_MEASUREMENT=en_US.UTF-8 LC_IDENTIFICATION=C
##
## attached base packages:
## [1] stats     graphics  grDevices utils     datasets  methods   base
##
## other attached packages:
##  [1] bindrcpp_0.2.2  skimr_1.0.3     magrittr_1.5    forcats_0.3.0
##  [5] stringr_1.3.1   dplyr_0.7.8     purrr_0.2.5     readr_1.3.0
##  [9] tidyr_0.8.2     tibble_1.4.2    ggplot2_3.1.0   tidyverse_1.2.1
##
## loaded via a namespace (and not attached):
##  [1] tidyselect_0.2.5  xfun_0.4          haven_2.0.0
##  [4] lattice_0.20-35   colorspace_1.3-2  generics_0.0.2
##  [7] htmltools_0.3.6   viridisLite_0.3.0 yaml_2.2.0
## [10] rlang_0.3.0.1     pillar_1.3.1      glue_1.3.0
## [13] withr_2.1.2       modelr_0.1.2      readxl_1.2.0
## [16] bindr_0.1.1       plyr_1.8.4        munsell_0.5.0
## [19] gtable_0.2.0      cellranger_1.1.0  rvest_0.3.2
## [22] evaluate_0.12     labeling_0.3      knitr_1.21
## [25] broom_0.5.1       Rcpp_1.0.0        scales_1.0.0
## [28] backports_1.1.3   jsonlite_1.6      hms_0.4.2
## [31] digest_0.6.18     stringi_1.2.4     grid_3.5.1
## [34] cli_1.0.1         tools_3.5.1       lazyeval_0.2.1
## [37] crayon_1.3.4      pkgconfig_2.0.2   xml2_1.2.0
## [40] lubridate_1.7.4   assertthat_0.2.0  rmarkdown_1.11
## [43] httr_1.4.0        rstudioapi_0.8    R6_2.3.0
## [46] nlme_3.1-137      compiler_3.5.1
```
